# Transcriptional Inactivation of TP53 and the BMP Pathway Mediates Therapy-induced Dedifferentiation and Metastasis in Prostate Cancer

**DOI:** 10.1101/2021.04.14.439569

**Authors:** Hyunho Han, Yan Wang, Josue Curto, Sreeharsha Gurrapu, Sara Laudato, Alekya Rumandla, Goutam Chakraborty, Xiaobo Wang, Hong Chen, Yan Jiang, Dhiraj Kumar, Emily Caggiano, Boyu Zhang, Yan Ji, Sankar N. Maity, Min Hu, Shanshan Bai, Ana Aparicio, Eleni Efstathiou, Christopher J. Logothetis, Nicholas Navin, Nora Navone, Yu Chen, Filippo G. Giancotti

**Author notes:** These authors contributed equally (Hyunho Han, Yan Wang). These authors contributed equally and are listed in alphabetical order (Josue Curto, Sreeharsha Gurrapu, Sara Laudato, Alekya Rumandla).

## Abstract

Unsupervised clustering and deconvolution analysis identifies a novel subtype of M-CRPC endowed with hybrid epithelial/mesenchymal (E/M) and luminal progenitor-like traits (Mesenchymal and Stem-like PC, MSPC). Analysis of patient datasets and mechanistic studies indicate that MSPC arises as a consequence of therapy-induced lineage plasticity. AR blockade instigates two separate and complementary processes: 1) transcriptional silencing of *TP53* and hence acquisition of hybrid E/M and stem-like traits; and 2) inhibition of the BMP signaling, which promotes resistance to the pro-apoptotic and anti-proliferative effects of AR inhibition. The drug-tolerant prostate cancer cells generated through reprogramming are rescued by neuregulin and generate metastases in mice. Combined inhibition of HER2/3 and AR or mTORC1 exhibit efficacy in preclinical models of mixed ARPC/MSPC or MSPC, respectively. These results identify a novel subtype of M-CRPC, trace its origin to therapy-induced lineage plasticity, and reveal its dependency on HER2/3 signaling.

## Introduction

Lineage plasticity, commonly encompassing dedifferentiation and transdifferentiation, plays a crucial role in tumor progression to metastasis and resistance to oncogene-targeted therapies (Boumahdi and de Sauvage, 2020; Gupta et al., 2019). A particular form of plasticity, neuroendocrine transformation, promotes resistance to EGFR inhibitors in lung cancer and to AR inhibitors in prostate cancer. In these cancers, inactivation of the tumor suppressor genes *TP53* and *RB1* promotes neuroendocrine transformation, downregulating expression of the initial oncogenic driver and activating alternative survival and proliferation pathways (Quintanal-Villalonga et al., 2020). In contrast, exposure to BRAF and MEK kinase inhibitors induces a drug-tolerant state in melanoma by directly reprogramming highly proliferative MITF^high^-AXL^low^ tumor cells into MITF^low^-AXL^high^ quiescent tumor cells (Arozarena and Wellbrock, 2019). A similar form of adaptive resistance has been observed in a variety of cancer cell lines exposed to multiple types of antimitotic therapy and has been associated with the acquisition of mesenchymal traits (Hangauer et al., 2017). The mechanisms that enable the epiclones of drug-tolerant cancer cells to reenter into the proliferative cell cycle are not known.

Prostate adenocarcinoma typically progresses from a hormone deprivation-sensitive stage to a castration-resistant and metastatic stage (CRPC), which becomes rapidly recalcitrant to therapeutic intervention (Attard et al., 2016; Logothetis et al., 2013). Several mechanisms of resistance to androgen receptor (AR) blockade have been proposed, including amplification and/or mutation of the *AR*, overexpression of the V7 splice variant of the AR, cooption of AR signaling by the Glucocorticoid Receptor (GR), and acquisition of mutations or activation of signaling pathways that alleviate the need for AR signaling, rendering tumor cells AR-independent (Watson et al., 2015). Next-generation AR inhibitors, such as enzalutamide and abiraterone, have improved patient survival but have not eradicated the disease (de Bono et al., 2011; Scher et al., 2012). About 30% of patients exhibit primary resistance to these agents and almost all responders eventually develop secondary resistance, highlighting the importance of identifying clinically relevant and targetable mechanisms of resistance to profound AR blockade.

Several observations suggest that exposure to next-gen anti-AR agents promotes the emergence of AR-independent populations of tumor cells, ultimately leading to treatment failure. A fraction of castration-resistant tumors exhibits variegated histology and a continuum of neuroendocrine characteristics, culminating in cases indistinguishable from *de novo* neuroendocrine or small cell prostate cancer (NEPC) (Beltran et al., 2011; Epstein et al., 2014). These tumors are often clinically aggressive and characterized by both bone and visceral metastases (Aggressive Variant Prostate Cancer, AVPC) (Aparicio et al., 2016). Experiments in LNCaP-AR cells and mouse models have indicated that inactivation of *TP53* and *RB1* or inactivation of *TP53* and exposure to abiraterone can convert *PTEN*-null prostate adenocarcinoma into therapy-resistant NEPC (Ku et al., 2017; Mu et al., 2017; Zou et al., 2017). In addition, direct transformation assays with luminal progenitors indicate that a combination of oncogenic mutations, including loss of *PTEN*, *TP53* and *RB1*, can drive the development of NEPC (Park et al., 2018). These findings suggest that, instead of directly inducing adaptive resistance, exposure to therapy favors the expansion of neoplastic clones that have acquired mutations leading to neuroendocrine transformation.

Recent studies have suggested that AR inactivation can promote the development of an AR-independent subtype of M-CRPC devoid of neuroendocrine traits (AR pathway-negative, NE-negative or Double Negative Prostate Cancer, DNPC) (Bluemn et al., 2017). In spite of this advance, the origin and nature of AR-independent tumor cells in many cases of M-CRPC remains unclear. In particular, it is unknown if the tumor cells in these cancers are cancer stem cells or if they have transdifferentiated to an alternative cell fate. Additional questions of whether they exist at the time of diagnosis or arise from therapy-induced lineage plasticity, and if this reprogramming can occur in the absence of *TP53* and *RB1* mutations are also pertinent. Finally, it is important to determine if the AR-independent tumor cells are intrinsically more malignant and metastatic compared to androgen driven tumor cells. We have here endeavored to address these questions by transcriptionally profiling M-CRPC datasets and tracing the origin of a newly defined subtype with mesenchymal and stem cell traits (MSPC) to therapy-induced lineage plasticity and transcriptional silencing of *TP53* and the BMP pathway.

## Results

### PAM Analysis Reveals Three Intrinsic Transcriptional Subtypes of Prostate Cancer

To identify the transcriptional subtypes of M-CRPC, we performed unsupervised clustering on the SU2C-PCF dataset (Robinson et al., 2015). Partition Around Medoids (PAM) and Principal Coordinates Analysis (PCoA) identified 4 clusters of samples, which correlated to a significant extent with metastatic site, presumably reflecting organ site-specific stromal gene expression or the effect of local tumor microenvironment on cancer cell gene expression (Figure S1A). To eliminate the effect of extrinsic tumor cell mechanisms, we applied the same approach to a large panel of CRPC <u>P</u>DXs, <u>c</u>ell lines, and tumor <u>o</u>rganoids (PCO samples; n=94) (Figure S1B) and identified three clusters of instrinsic tumor cell transcriptional programs, which were validated as stable and robust by using Consensus Clustering and the Proportion of Ambiguously Clustered pairs (PAC) metric (Figures 1A left and S1C). Non-linear methods, such as UMAP or tSNE, also identified three clusters (unpublished data). Pairwise comparison of the gene expression programs of the three clusters led to the definition of 2,424 genes (FDR < 0.05 and fold change > 2), which retained an intact clustering ability when compared to the whole transcriptome (Figure 1A middle, Supplementary Table 1). Gene Set Enrichment Analysis (GSEA) identified the top signatures that define the two coordinates of the PCoA (Figure 1B) and revealed that cluster 1 is dominated by the expression of AR target and lipid oxidation genes, cluster 2 by genes involved in immune suppression, fatty acid biosynthesis, and TGF-β signaling, and cluster 3 by genes associated with neuronal differentiation and negative regulation of SOX9 (Figure S1D). We concluded that unsupervised clustering enables the separation of CRPC PCOs into three robust and stable transcriptional subtypes.

**Figure 1.**
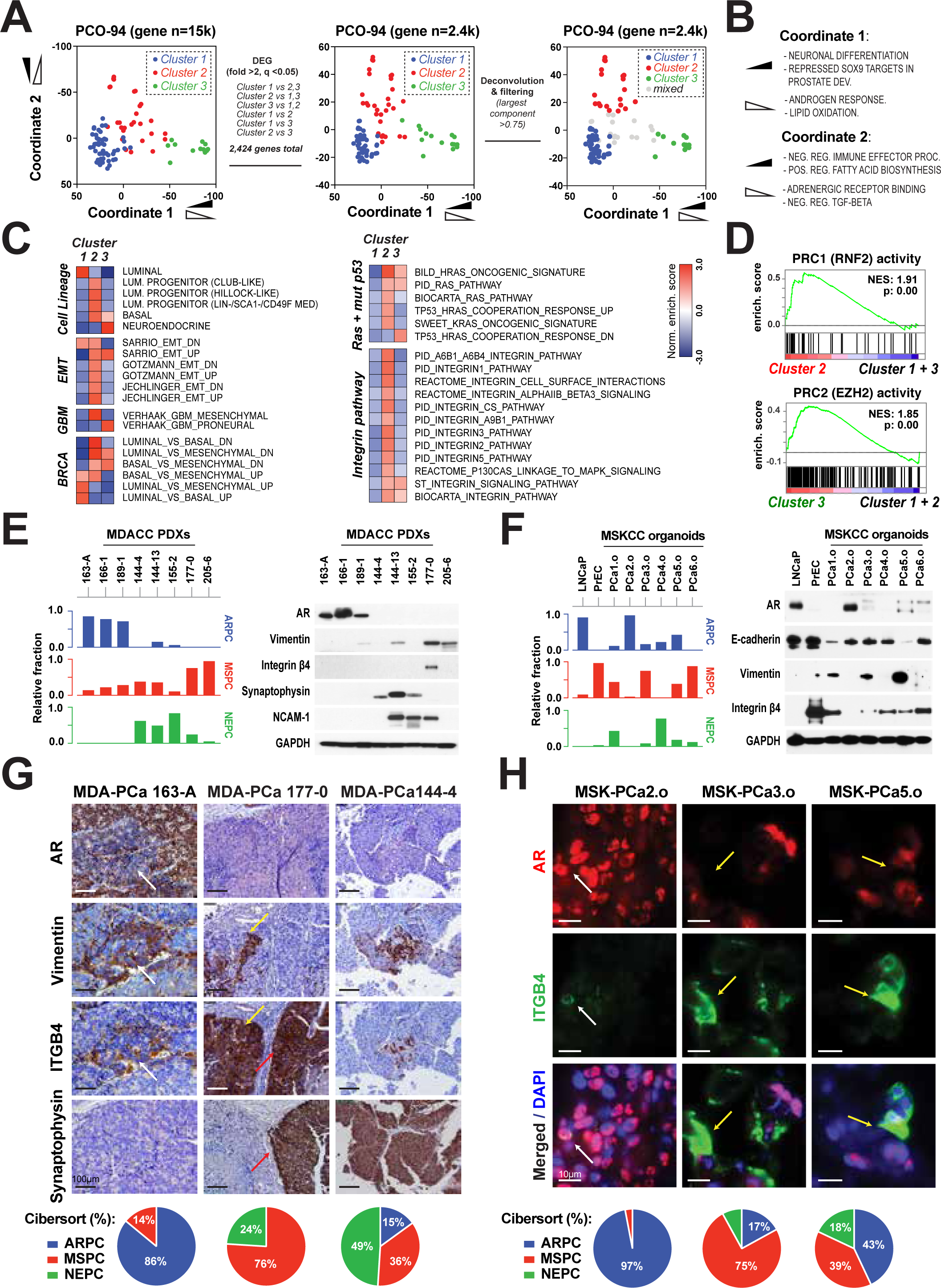
Prostate Cancer Experimental Models Transcriptomic Subtyping. (A) Left: principal coordinate analysis (PCoA) plot of the PCO-94 dataset clustered by portioning around medoids (PAM) method. Middle: PCoA plot of the reduced PCO-94 dataset with repeating PAM clustering. Right: PCoA plot of the reduced PCO-94 dataset, with purity annotation by deconvolution analysis. Purity was defined by the estimated fraction of each cluster types in a sample (“mixed” if the largest fraction < 75%). (B) Principal coordinates 1, 2 from the PCO-93 PCoA plot and top correlating gene sets among the C2 “curated”, the C5 “Gene Ontology” and the H “hallmark” gene sets (from the mSigDB collections). Ranking by Pearson correlation “r” of ssGSEA scores and the coordinate values. Black-filled angle: negative correlation. Blank angle: positive correlation (detailed results are shown in Figure S1D). (C) GSEA Normalized Enrichment Score (NES) heatmap of three clusters, comparing one cluster versus the rest. Results were shown together categorically. Cell Lineage: normal human prostate epithelial population defined by single cell RNA sequencing (Henry et al., 2018). EMT (Epithelial to Mesenchymal Transition): Breast cancer cell line MCF 10A undergoing EMT (Sarrio et al., 2008); Hepatocyte MMH-RT response to TGF-beta (Gotzmann et al., 2006); Ras-transformed mammary epithelial cell EpH4 response to TGF-beta (Jechlinger et al., 2003). GBM (Glioblastoma Multiforme): gene expression-based molecular subtypes of GBM (Verhaak et al., 2010). BRCA (Breast Cancer): gene expression-based molecular subtypes of BRCA (Charafe-Jauffret et al., 2006). “Ras + mut p53” gene sets were collected by using the key word “Ras”, “Integrin pathway” genesets by “Integrin” from the MSigDB. (D) Polycomb Repressive Complex (PRC) 1 and 2 activities. GSEA was performed in comparison of cluster 2 versus the rest (top) or cluster 3 versus the rest (bottom). PRC1 activity gene set was from our group’s publication, “genes upregulated by RNF2” (Su et al., 2019). For PRC2 activity, PRC2_EZH2_UP.V1_UP (M2737) from the MSigDB was used (Bracken et al., 2006). (E and F) Predicted relative fractions (0 to 1) of ARPC (cluster 1 renamed, blue color), MSPC (cluster 2, red) and NEPC (cluster 3, green) (bar graph) and representative protein expressions (immunoblot) in MDACC PDXs (E) and MSKCC organoids (F). (G and H) Immunohistochemistry stainings (G) and Immunofluorescence stainings (H) of AR, ITGB4 and Vimentin, Synaptophysin in three PDXs MDA-PCa-163-A, 177-0, and 144-4 and organoids PCa-2, PCa-3 and PCa-5. Representative region of heterogeneity are shown. Scale bar = 100um (G); 10um (H). Pie chart (below) presents the relative fractions (0 to 100%) of ARPC, MSPC and NEPC.

We examined in detail the gene expression programs active in each PCO cluster and their relationship to previously defined subtypes (Supplementary Table 2). Consistent with their AR pathway activity, cluster 1 PCOs are enriched for signatures associated with primary prostate adenocarcinoma, prostate luminal differentiation, and luminal breast cancer, and they thus correspond or overlap with ARPC. In contrast, cluster 3 PCOs express signatures associated with prostate neuroendocrine cells and the proneural subtype of Glioblastoma Multiforme (GBM) (Figures 1C left and S1E), display high PRC2 (EZH2) activity (Figure 1D), and include all the PDXs derived from small cell carcinomas (Tzelepi et al., 2012), suggesting that they represent NEPC. Finally, considering the absence of AR pathway activity or neuroendocrine traits in cluster 2 samples, we posited that these samples may correspond to or overlap with DNPC (Bluemn et al., 2017). Intriguingly, cluster 2 PCOs are enriched for luminal progenitor signatures, which are shared by ‘club-like’ and ‘hillock-like’ cells in the prostate and the respiratory tract (Henry et al., 2018; Sackmann Sala et al., 2017). Moreover, they express several relevant cancer signatures associated with the EMT, the mesenchymal subtype of GBM, and basal-like breast cancer (Figure 1C, left). Based on their transcriptional profiles, we defined cluster 1 as AR-pathway positive Prostate Cancer (ARPC), cluster 2 as Mesenchymal and Stem-like PC (MSPC), and cluster 3 as Neuro-Endocrine PC (NEPC). Further analysis indicated that MSPC is enriched for signatures activated during oncogene-dependent prostate tumorigenesis in mouse models (Acevedo et al., 2007; Azare et al., 2007; Liu et al., 2009) and signatures reflective of RAS and mutant TP53 pathway activation and integrin signaling (Figure 1C, right and S1E). Foreshadowing their immunosuppressive nature, cluster 2 samples also displayed an enrichment of the PRC1 (RNF2) signature, which we previously linked to both stemness and immune evasion in DNPC (Su et al., 2019), as well as several immune signaling and inflammatory signatures (Figures 1D and S1E).

Transcriptional subtyping has revealed intratumoral heterogeneity in glioblastoma and breast cancer PCOs (Bierie et al., 2017; Neftel et al., 2019; Wagner et al., 2019; Wang et al., 2017). To examine if the prostate cancer PCOs harbor intratumoral heterogeneity, we used CIBERSORT deconvolution analysis of transcriptional data (Newman et al., 2015). The results revealed that, although the majority of samples in each cluster consisted of tumor cells endowed with a homogeneous transcriptional profile (>75% pure), 20% of the samples mapped at the boundaries between the three clusters and consisted of admixtures of cancer cells with distinct transcriptional profiles (<75% pure) (Figures 1A right and S1F). All 8 cell lines showed high purity (Figure S1G). Early and late passage organoid cultures and PDXs maintained a similar transcriptional composition (Figure S1H). Moreover, organoid-derived xenografts (ODXs) maintained the same transcriptional composition of the tumor organoids from which they were generated, suggesting that propagation *in vitro* or *in vivo* does not favor the emergence and dominance of a specific transcriptional state (Figure S1I). Similar conclusions were reached independently of whether the transcriptional profiles were generated from DNA microarray or RNA sequencing data (Figure S1J). Thus, although the majority of PCO samples can be grouped into three major subtypes, based on the transcriptomes of the majority of their constituent cells, a fraction of samples are admixtures of tumor cells characterized by two or three of the identified transcriptional states.

To validate these findings, we examined the expression of AR and luminal differentiation markers, mesenchymal and stem cell markers, and NE markers in a subset of MDACC PDXs and MSKCC organoids, which had been subjected to CIBERSORT deconvolution (Figure 1E and 1F, left). Immunoblotting documented prominent expression of the AR in samples containing a significant ARPC component (>50%) and synaptophysin in those consisting of a significant NEPC component (>50%). Moreover, it detected robust expression of the mesenchymal marker vimentin and/or the prostate stem cell marker ITGB4 (Yoshioka et al., 2013) in samples with a predominant MSPC component (>50%) (Figure 1E and F, right). Intriguingly, immunohistochemical staining of PDXs indicated that the NE and mesenchymal or stem cell markers were restricted to subpopulations of tumor cells, consistent with intratumoral heterogeneity (Figure 1G). In addition, immunofluorescent staining of tumor organoids revealed a mutually exclusive expression of AR and the stem cell marker ITGB4. Although ARPC organoids contained predominantly AR^+^ tumor cells, they also exhibited scattered ITGB4^+^ tumor cells. In contrast, MSPC organoids displayed the opposite pattern of expression. Organoids that could not be readily classified as ARPC or MSPC comprised similar proportions of AR^+^ and ITGB4^+^ tumor cells, again consistent with intratumoral heterogeneity (Figure 1H). Finally, CIBERSORT deconvolution indicated that a fraction of LuCaP PDXs and organoids consist of admixtures of two or three of the transcriptional subtypes (Figure S1H). Therefore, the PDXs and organoids consist of different proportions of tumor cells expressing ARPC, MSPC, or NEPC transcriptional programs.

### MSPC Exhibits Genetic Alterations Similar to ARPC and is Associated with Advanced Stage, Poor Prognosis and AR Pathway Inhibition

To test the discriminatory power of instrinsic tumor cell transcriptional subtyping in patients, we used the 2,424 genes differentially expressed across the three PCO clusters to perform clustering and deconvolution on the metastatic samples from the SU2C-PCF (Robinson et al., 2015), FHCRC (Kumar et al., 2016), and UCSF datasets (Quigley et al., 2018). Through this approach, we identified three distinct clusters of M-CRPC samples expressing the ARPC, MSPC, or NEPC transcriptional signature (Figure S2A). Notably, analysis of the prevalence of MSPC amongst subtypes of M-CRPC indicated an increase from 19% in the oldest dataset (FHCRC) to 36% in the most recent dataset (UCSF), suggesting that exposure to second-generation AR inhibitors, such as enzalutamide and abiraterone, may contribute to the emergence of MSPC. After filtering out mixed samples by deconvolution (<60% pure), we generated subtype-specific gene expression heatmaps of the three patient datasets (Figure 2A, Supplementary Table 3). Examination of the prevalence of each subtype at metastatic sites revealed an enrichment of ARPC in lymph node and bone metastases, MSPC in bone and liver metastasis, and NEPC specifically in liver metastasis, suggesting that each subtype is characterized by a preferential pattern of metastatic colonization (Figure 2B).

**Figure 2.**
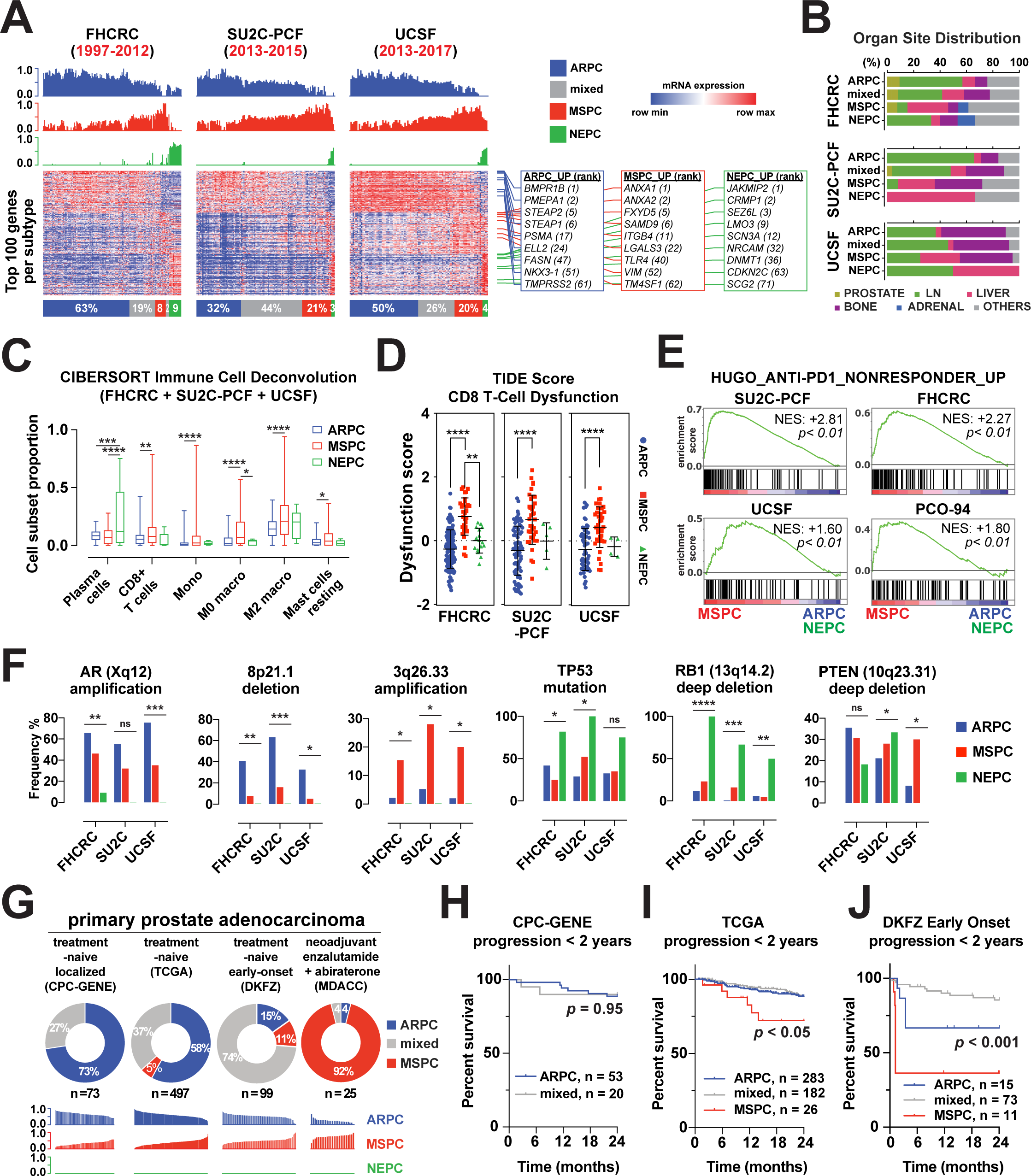
Transcriptomic Subtypes of Metastatic Castration-Resistant Prostate Cancer and Their Characteristics. (A) (Top) ARPC, MSPC and NEPC relative fraction bar graphs in human M-CRPC datasets, reported from FHCRC (Kumar et al., 2016), SU2C-PCF (Robinson et al., 2015) and UCSF (Quigley et al., 2018). Bracketed n: years samples collected. Sample data are aligned with the heatmap column order. (Bottom) Top 100 upregulated genes expressions per subtype. Subtypes were assigned by the largest relative fraction components from deconvolution analysis (“mixed” if the largest fraction < 0.6). Columns were grouped by subtype and sorted by hierarchical clustering. (right lower) selected upregulated genes per subtype. bracketed n: mRNA expression ranks, calculated by lowest max false discovery rate (FDR max) from the three datasets. (B) Tissue site distribution by M-CRPC subtype. (C) CIBERSORT immune cell population deconvolution analysis on M-CRPC. Immune cell population deconvolution analysis (LM22) on M-CRPC data was performed as previously described (Su et al., 2019). Pure ARPC, MSPC and NEPC samples defined in panel A were merged in this presentation. P value calculated by One-way ANOVA. All other immune cell population distributions were not significantly different. Box and whiskers plot (min to max). (D) Tumor Immune Dysfunction and Exclusion (TIDE) scoring analysis (Jiang et al., 2018). CD8^+^ T-Cell Dysfunction score of each subtype is shown. P value calculated by One-way ANOVA. Multiple comparison by Dunnett’s test. (E) Enrichment plots of Anti-PD-1 therapy nonresponder geneset in MSPC versus the other subtypes. GSEA performed in three M-CRPC datasets and one PCO-94 dataset. Gene set was generated by selecting genes upregulated in metastatic melanoma anti-PD-1 nonresponders compared to responders (Mann-Whitney test p < 0.01) (Hugo et al., 2016). NES = normalized enrichment score. (F) Signature genetic alterations of ARPC, MSPC and NEPC, in three M-CRPC datasets. AR amplification, chromosome 8p21.1 deletion (shallow and deep), chromosome 3q26.33 amplification, PTEN and RB1 deletions, and TP53 mutations are shown. P value calculated by chi-square test. ns = not significant; *: P ≤ 0.05; **: P ≤ 0.01; ***: P ≤ 0.001; ****: P ≤ 0.0001. (G) Subtype classification of primary prostate adenocarcinoma. Deconvolution and purity analysis (cut-off value: 0.6) same as used in M-CRPC (Panel A). Below is relative fraction prediction provided by CIBERSORT analysis. Note that NEPC is zero, likely due to histologic criteria and the rarity of primary *de novo* neuroendocrine carcinoma in prostate. n = number of samples with RNA expression data. (H-J) Kaplan-Meier plots of progression-free survival, in CPC-GENE (H), TGCA (I), and DKFZ early onset (J). Log-rank (Mantel-Cox) test for comparison of survival curves. Progression was defined as biochemical recurrence after primary therapy in each study. In DKFZ, PCA034 was excluded from analysis (T01,2,4,6 classified as mixed, T03,5 as ARPC). All other samples assignments were consistent within a patient. Survival data acquired from cBioPortal.org.

We next directly examined the relationship between our classification based on spontaneous clustering of transcriptional programs and the most recent molecular classification based on the expression of gene sets identified through biomarker analysis (AR/PSA and SYP/CHGA) (Bluemn et al., 2017). As anticipated, we found that MSPC largely overlaps with double negative prostate cancer (DNPC), which is negative for AR/PSA and SYP/CHGA protein expression (Figure S2B). Parenthetically, we note that the extent of overlap between MSPC and DNPC is similar to that observed between the basal-like intrinsic subtype of breast cancer and triple-negative breast cancer (75-80%) (Foulkes et al., 2010). However, MSPC samples comprise a larger group of metastases as compared to marker and gene set-defined DNPC samples. In fact, they also include metastases previously classified as AR^+^NE^-^ or AR^-^NE^+^ (Figure S2B). We recently found that PRC1 promotes immune suppression and neoangiogenesis in bone metastases by inducing recruitment of M2 macrophages and regulatory T cells (Su et al., 2019). Consistent with the widespread activation of PRC1 in MSPC (Figure 1D), CIBERSORT deconvolution revealed an enrichment of myeloid cells, including M2 macrophages, in MSPC across the three M-CRPC datasets. Although this analysis revealed that CD8^+^ T cells are also enriched in MSPC (Figure 2C), TIDE scoring (Tumor Immune Dysfunction and Exclusion) (Jiang et al., 2018) indicated that these T cells are highly dysfunctional in spite of elevated intratumoral interferon-gamma (IFNL) activity (Figures 2D and S2C). Consistently, GSEA indicated that MSPC is enriched for an anti-PD-1 nonresponder gene expression signature (Figure 2E, Supplementary Table 4) (Hugo et al., 2016). These results corroborate the conclusion that MSPC is characterized by a distinctly immunosuppressive microenvironment as compared to ARPC or NEPC.

Analysis of the most frequently mutated cancer genes indicated that *PTEN* deletions and mutations and *ERG* fusions are common in all three transcriptional subtypes of M-CRPC. In contrast, *AR* and *FOXA1* amplifications and mutations are common in ARPC and MSPC, but not NEPC. Conversely, *TP53* and *RB1* deletions and mutations were considerably enriched in NEPC (Figures 2F and S2D), in agreement with its postulated origin (Ku et al., 2017; Mu et al., 2017). To identify the oncogenic alterations that may distinguish ARPC and MSPC, we examined chromosomal structure variations. In agreement with the observation that lipid oxidation is one of the top pathways enriched in ARPC (Figure S1D), we found that this subtype is characterized by frequent deletion of Chromosome (Chr) 8p21.1 (both shallow and deep), which inactivates multiple genes involved in lipid metabolism and resistance to anti-cancer drugs (Cai et al., 2016; Xue et al., 2012). In contrast, the amplification of Chr 3q26.33 was moderately enriched in MSPC patient datasets (10-20% across datasets), organoids, and cell lines (Figures 2F, S2D, and S2E). This chromosomal segment comprises three oncogenes, *SOX2*, *FXR1, and PRKCI*, which could contribute to MSPC (Bass et al., 2009; Justilien et al., 2014; Qian et al., 2015). Finally, as anticipated, we noted a large enrichment of the homozygous deletion of Chr 13q14.2, which comprises *RB1*, in NEPC (Figures 2F and S2D). These findings suggest that MSPC is closely related to ARPC, whereas NEPC is distinguished by *TP53* and *RB1* mutations.

To examine if MSPC can originate at the primary site, we performed deconvolution analysis on the transcriptomes of three independent primary prostate adenocarcinoma datasets comprising samples of varying clinical characteristics. In addition to the TCGA dataset, which consists of treatment naïve prostate cancers of all T stages, Gleason scores, and patient ages, we examined the CPC-GENE dataset, which includes treatment naïve nonaggressive localized prostate cancers (Gleason score ≤ 7) (Fraser et al., 2017), and the DKFZ dataset, which comprises early-onset treatment naïve prostate cancers (patient age < 55) (Gerhauser et al., 2018). Although the CPC-GENE dataset did not contain pure MSPC samples (0%), the TCGA and, even more so, the DKFZ dataset comprised increasing proportions of MSPC (5% and 12%, respectively) (Figure 2G). RPPA analysis confirmed decreased expression of the *AR* and increased expression of the top-ranked MSPC gene *ANXA1* in primary MSPC samples (Figure S2G). There was no NEPC component in any of the datasets, consistent with the exclusion of small cell neuroendocrine histology. Intriguingly, we found that primary MSPC samples are enriched for *TP53* mutations (31-54%) and *PTEN* deletions (39%), but not *SPOP* and *FOXA1* mutations or *RB1* deletions (Figures S2F and S2H). Moreover, primary MSPC samples were in general more advanced than primary ARPC or mixed samples in terms of Gleason score, pathologic T stage, and N stage (Figure S2I-K). Accordingly, primary MSPC cases were associated with a significantly worse progression-free survival as compared to other cases (Figure 2H-J). These findings suggest that MSPC can arise during the evolution of primary prostate cancer and lead to accelerated progression to dissemination and metastasis.

To examine if MSPC can arise as a consequence of therapeutical blockade of the AR, we applied deconvolution analysis to the transcriptional profiles of a MDACC dataset consisting of localized high risk prostate cancers (MDACC dataset, Gleason score ≥ 8 or clinical stage ≥ T2b), which had not responded to neoadjuvant treatment with enzalutamide, abiraterone, and leuprolide for 24 weeks (Efstathiou et al., 2016). Strikingly, the large majority of these resistant tumors consisted of pure MSPC (92%). Although the patients in this trial did not undergo a pretreatment biopsy, the dominance of pure MSPC in the resistant tumors as compared to its rarity in the historical control group suggests that MSPC may arise as a consequence of exposure to AR inhibitors.

### Enzalutamide Induces Dedifferentiation to a Therapy Resistant Hybrid Epithelial/Mesenchymal and Stem Cell Fate Resembling MSPC

To examine the effect of AR blockade on prostate cancer luminal differentiation, we treated LNCaP cells with 10 μM enzalutamide and conducted transcriptional analysis over a 2 week period. LNCaP cells harbor a deletion of *PTEN* but are *TP53* and *RB1* proficient (Li et al., 1997; Mu et al., 2017; van Bokhoven et al., 2003). As anticipated, enzalutamide inhibited cell growth and provoked apoptosis of an increasing proportion of LNCaP cells at each of the time points examined (Figure S3A). However, about 20% of tumor cells survived 14 days of drug treatment and remained viable over an additional period of 2 weeks, suggesting that they had become drug-tolerant. RNA-seq followed by PCoA and GSEA indicated that LNCaP cells exposed to enzalutamide downregulate signaling through mTORC1 and exit the cell cycle (Figure S3B). Superimposition of the timeseries of transcriptomic profiles of persister LNCaP cells onto the PCoA plot of ARPC, MSPC and NEPC PCOs revealed that the LNCaP cells progressively change their transcriptional program from ARPC to MSPC over 2 weeks of enzalutamide treatment (Figure 3A). As anticipated, this conversion was associated with the downregulation of AR target genes and cell cycle genes and the upregulation of EMT and stemness genes (Figure 3B). The drug-tolerant LNCaP cells exhibited diminished levels of the AR and of the epithelial tight junction marker ZO-1. Decreased levels of the AR in enzalutamide-treated cells had been noted in earlier studies and attributed to diminished mRNA expression and protein stability (Kuruma et al., 2013; Tran et al., 2009). In addition, although the LNCaP cells treated with enzalutamide maintained high levels of E-cadherin, they acquired expression of the mesenchymal proteins vimentin and fibronectin and the prostate stem cell marker ITGB4 (β4 integrin) (Figure 3B, C). Notably, prior studies have linked expression of ITGB4 to the hybrid epithelial/mesenchymal (E/M) state associated with stemness in breast cancer (Bierie et al., 2017). These observations suggest that the LNCaP cells become tolerant to enzalutamide by acquiring hybrid E/M and stem-like traits and switching their fate from ARPC to MSPC.

**Figure 3.**
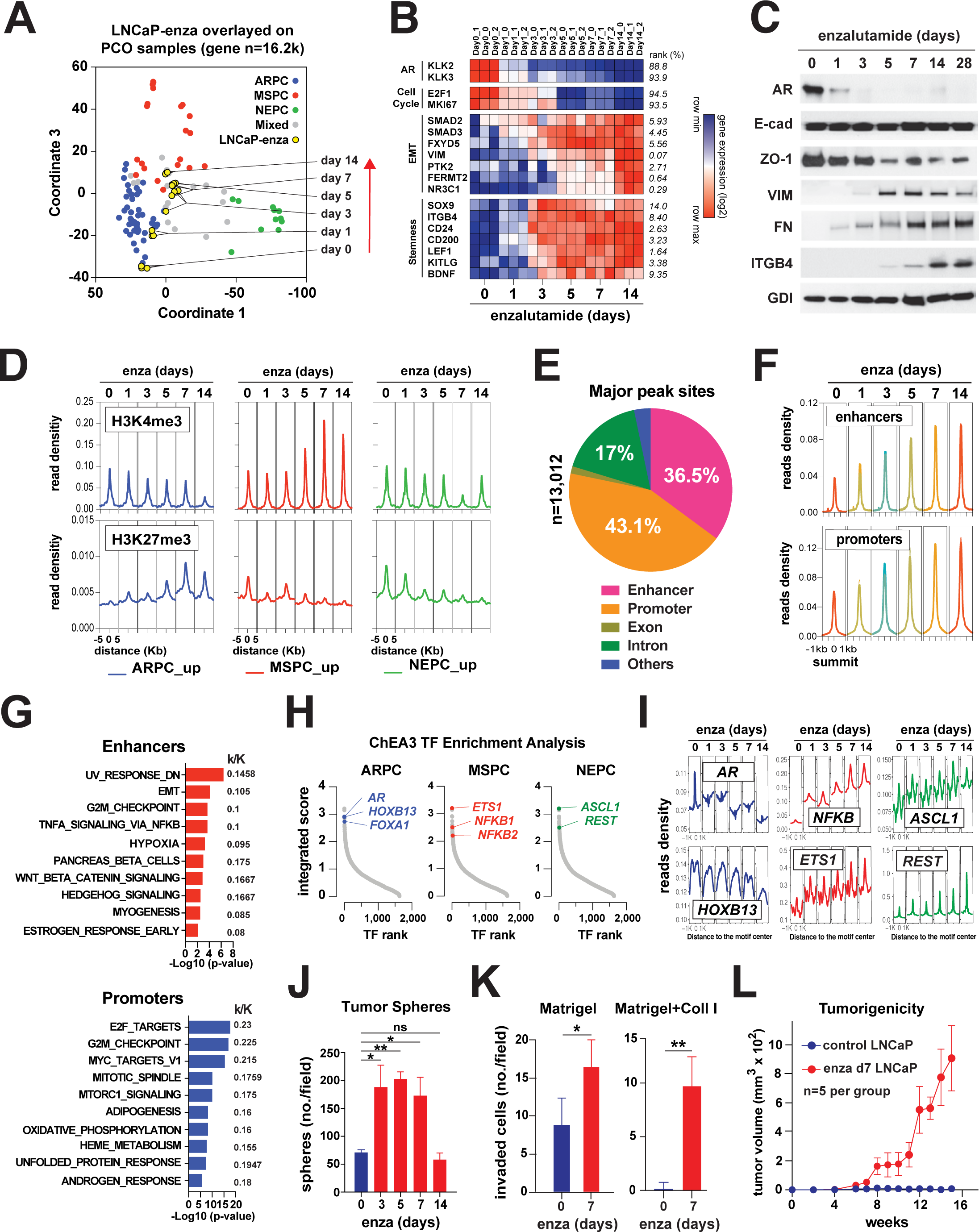
Prostate Cancer Reprogramming to Mesenchymal and Stem-like State by Androgen Receptor Blockade. (A) PCoA plot of LNCaP enzalutamide time series RNA-seq data merged with PCO-94 dataset. Three replicates per time point. Note that the general distribution of PCO-94 samples remains (compared to Figure 1A) while previous coordinate 2 is now coordinate 3 (shown as Y-axis here). The new coordinate 2 represents cell cycle and mTOR activity, which showed highest correlation with coordinate 2 values (shown in Figure S3B). (B) Gene expression heatmap. Categorized into AR (target genes), Cell Cycle, EMT, and Stemness. Rank (%, right, italic) by Pearson correlation coefficient with drug incubation time point values (0-14, days). (C) AR, EMT and stemness markers protein expression by Western Blot in LNCaP enzalutamide time-series. (D) Promoter histone marks shift. H3K4me3 (upper) and H3K27me3 (lower) peaks of ARPC, MSPC and NEPC upregulated genes (defined in Figure 1A) in LNCaP enzalutamide time series histone marker Chip-sequencing. Distance = distance from transcription start site. (E-G) LNCaP enzalutamide time series ATAC-seq data. ATAC = Assay for Transposase-Accessible Chromatin using sequencing. (E) Pie plot showing classification of “major” peak sites (defined in Figure S3G). (F) Enhancer sites and promoter regions in “major” peak summit. Average of duplicates. (G) The hallmark gene set enrichments for “major” enhancer sites peaks and promoter regions peaks. k/K: overlapping n (k) of genes per total n (K) of genes in each geneset. (H) Transcription factor (TF) enrichment analysis using ChEA3 (Keenan et al., 2019) in ARPC, MSPC and NEPC upregulated genes. (I) ARPC, MSPC and NEPC TFs binding motif accessibility shift in LNCaP cells treated with enzalutamide. (J-L) Functional Assays. (J) Number of spheres (per 3,000 cells) from LNCaP cells treated with enzalutamide. (K) Migration (matrigel matrix) and invasion (matrigel + collagen matrix) assays of LNCaP cells with or without 7-day enzalutamide pretreatment. Student’s t-test. (L) Castration-resistant tumor growth *in vivo*. LNCaP cells (0.3 x 10^6) with or without 7-day enzalutamide pretreatment were mixed with 50% matrigel and implanted subcutaneously in castrated mice. Tumor growth was monitored for 16 weeks. Fiver mice per group.

To examine the generality of these findings, we tested the VCaP cells and the MDA PCa-163-A PDX model, which was established from a treatment naïve patient (Aparicio et al., 2016) and displays a predominant ARPC phenotype (Figure 1E and 1G). In agreement with their high sensitivity to the drug (Tran et al., 2009), the majority of VCaP cells underwent cell death within 5 days of treatment with 0.2 μM enzalutamide. However, the cells persisting throughout drug exposure downregulated expression of the AR and the AR target gene TMPRSS2 but maintained the expression of E-cadherin and upregulated that of vimentin and ITGB4, consistent with the acquisition of hybrid E/M and stemness traits (Figure S3C and S3D). Prostate adenocarcinoma cells explanted from the MDA PCa-163-A PDX model underwent a similar phenotypic conversion *in vitro* (Figure S3E). In addition, exposure to enzalutamide of mice bearing MDA PCa-163-A PDX tumors induced similar changes, including a downregulation of AR and an upregulation of ITGB4 and vimentin, *in vivo* (Figure S3F). These observations indicate that enzalutamide induces mesenchymal and stem-like traits in multiple prostate adenocarcinoma models.

To examine the nature of the phenotypic conversion from ARPC to MSPC, we conducted an integrated epigenomic analysis on LNCaP cells treated with enzalutamide over a 2 week period. ChIP-seq indicated that exposure to the drug decreases the deposition of the repressive histone mark H3K27me3 and increases the deposition of the activation mark H3K4me3 on the promoters of MSPC marker genes. It also induces reciprocal changes in the deposition of the two marks on the promoters of ARPC marker genes (Figure 3D). Parenthetically, the promoters of NEPC marker genes gradually lost H3K27me3 during enzalutamide treatment but did not acquire higher levels of H3K4me3, suggesting that they had become poised for activation (Figure 3D). To examine the genome-wide landscape of chromatin accessibility and its association with gene expression, we conducted ATAC-seq and GSEA. PCA and annotation of the major peaks of chromatin accessibility pointed to a progressive opening of the chromatin at enhancers and promoters in cells treated with enzalutamide (Figures 3E, F, and S3G-I). GSEA indicated that the regulatory regions becoming hyperaccessible in response to the drug control the expression of batteries of genes linked to stem cell activity, cell signaling, cell fate, cell cycle, the response to androgen, and other functions (Figure 3G). In consonance with the observed phenotypic conversion, the chromatin accessibility of the enhancers and promoters of EMT and stemness genes increased substantially during exposure to enzalutamide (Figure S3J). Furthermore, ChIP-seq indicated that the deposition of the repressive mark H3K27me3 decreased and that of the activation mark H3K4me3 increased at promoters of protypical mesenchymal and stemness genes (Figure S3K). Finally, although AR inhibition coordinately increased the accessibility of the enhancers and the transcription of genes within mesenchymal, inflammatory, and stem cell signatures, it increased the accessibility but decreased the activity of the promoters of “ANDROGEN_RESPONSE”, “E2F_TARGETS”, “G2M_CHECKPOINT”, “MTORC1 SIGNALING”, and “WNT_BETA_CATENIN_SIGNALING” genes (Figure S3L). This latter observation is consistent with the model that hyperaccessible regions can facilitate the recruitment of activator or repressor complexes depending on mass action and chromatin context (Klemm et al., 2019).

To identify the top transcription factors (TFs) dominating the ARPC, MSPC and NEPC landscape, we used ChEA3 TF Enrichment Analysis (Keenan et al., 2019). We found that the ETS1 and NFKB1/2 cistromes dominate the transcriptional program of MSPC. In contrast and as anticipated, the AR, FOXA1, and HOXB13 cistromes were prevalent in ARPC and the ASCL1 and REST cistromes in NEPC (Figure 3H, Supplementary Table 5) (Kron et al., 2017; Labrecque et al., 2019; Park et al., 2018; Pomerantz et al., 2015; Pomerantz et al., 2020). In addition, whereas enzalutamide decreased the chromatin accessibility of the DNA binding motifs of ARPC-specific TFs AR, FOXA1 and HOXB13, it increased that of MSPC-specific TFs NFκB and ETS1. However, the drug also increased the accessibility of the DNA binding motifs for ASCL1 and REST, which do not dominate the MSPC transcriptional program (Figure 3H and I). In fact, ssGSEA indicated that enzalutamide treatment downregulates the expression of ASCL1 and REST target genes. Therefore, LNCaP cells exposed to enzalutamide exhibit coordinated and dynamic changes in chromatin accessibility and transcriptional activity culminating in the downregulation of the AR-driven cistrome and the induction of a transcriptional program dominated by ETS1 and NFKB1/2.

To examine the functional consequences of the drug-tolerant state induced by enzalutamide, we examined the neoplastic traits of LNCaP cells persisting after drug treatment. The cells that were exposed to enzalutamide for 1 week exhibited a robust increase in tumorsphere formation, migration, and invasion in vitro compared to untreated cells (Figure 3J-L). This effect decreased during the second week of treatment possibly due to consolidation of proliferative quiescence (Figure 3J). Moreover, the LNCaP cells that were treated with enzalutamide for 1 week were able to produce subcutaneous tumors in castrated mice after 1 month of latency, whereas the parental controls were not tumorigenic under these conditions (Figure 3L). Thus, although the enzalutamide tolerant LNCaP cells are slowly cycling or quiescent in vitro, they manifest oncogenic traits associated with prostate cancer stem cells, including high self-renewal capacity *in vitro* and castration resistance *in vivo*.

### Single Cell Analysis Delineates the Trajectory from ARPC to MSPC

To further examine the nature of the phenotypic and functional conversion of enzalutamide-treated LNCaP cells, we performed single cell RNA sequencing (scRNA-seq) at various time points over a 7 day period. Graph-based clustering defined six closely related cell clusters (Figures 4A and S4A). Pseudotime analysis suggested that the cells exposed to enzalutamide transit from cluster 5 through cluster 1 and 2 to cluster 4. Cluster 4 finally morphs into cluster 3 and to a lower extent into cluster 6 (Figure 4A). Transcriptional analysis indicated that cluster 5 and 1 cells, which are adjacent in high dimensional space, exhibit AR signaling and MYC activity but are distinguished because of the expression of cell cycle genes in cluster 5 cells. In contrast, cluster 2 and cluster 4 cells show inflammatory and quiescence traits. Finally, whereas cluster 6 cells have mesenchymal features, cluster 3 cells display hybrid E/M and stem cell traits (Figure S4A). Consistent with the pseudotime trajectory, analysis of individual timepoint data indicated that cluster 5 cells, which are characterized by the expression of G2/M cell cycle genes, disappear following enzalutamide treatment. In contrast, cluster 3 cells with hybrid E/M and stem cell traits increase from 7% to 40% and become the predominant cluster over 7 days of treatment (Figure 4B). Corroborating the transition from cluster 5 to cluster 3 deduced from the pseudotime analysis, re-analysis of the ChIP-seq data indicated that cluster 5 genes are deactivated and repressed, whereas cluster 3 genes are de-repressed and activated during the time course of enzalutamide treatment (Figure S4B).

**Figure 4.**
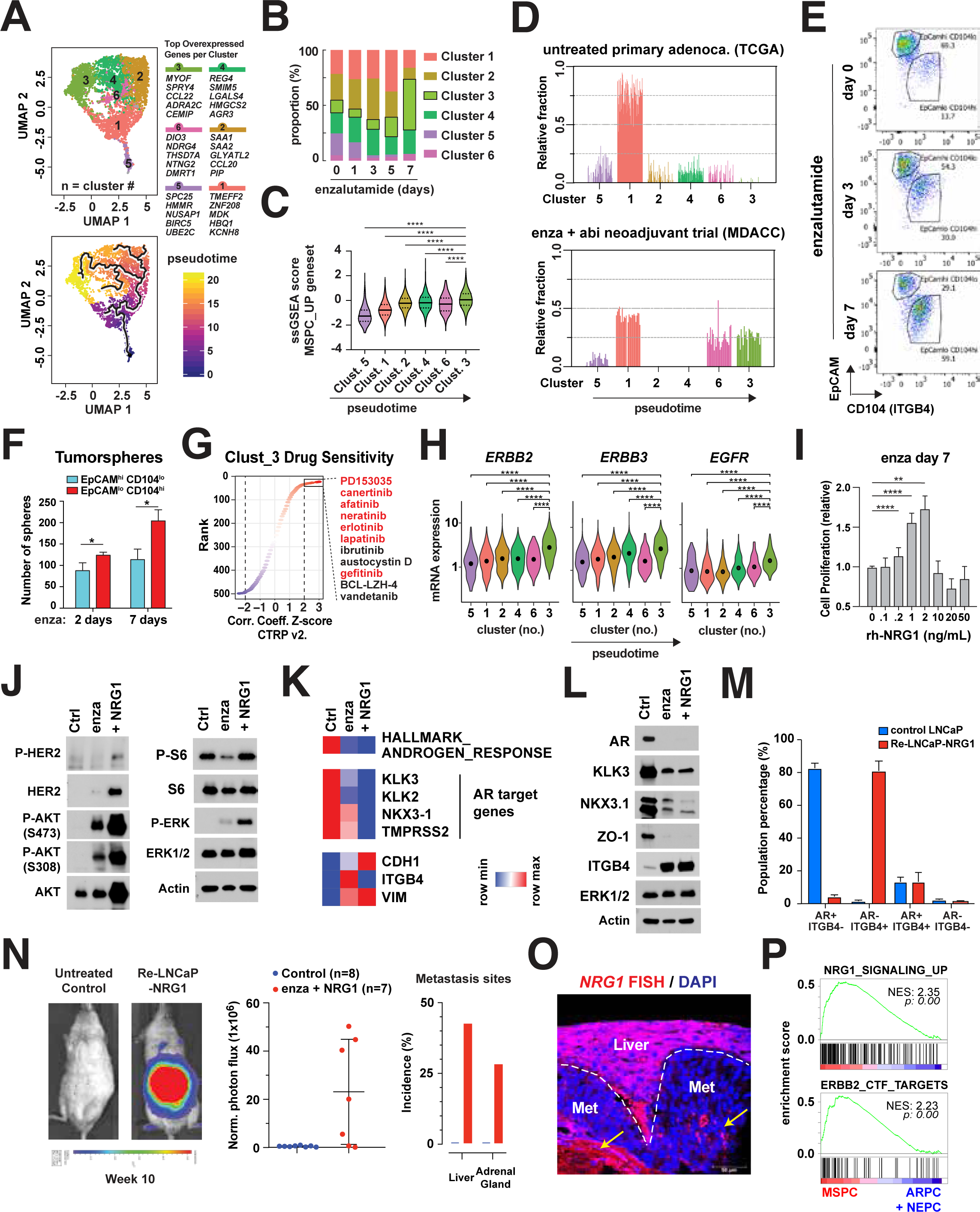
ERBB2 as reactivation cue in reprogrammed LNCaP cells. (A-D) Single cell RNA-seq analysis of LNCaP reprogramming to mesenchymal and stem-like state. (A) (Top) UMAP plot of LNCaP enzalutamide timeseries (day 0, 1, 3, 5, 7 merged). Clusters identified by graph-based clustering. Two clusters of low UMI (gray) excluded from further analysis. (Bottom) Pseudotime analysis and cell fate trajectory by Monocle 3. Gene expression characteristics with trajectory order is summarized in Figure S4A. (B) Clusters proportion in each time point. (C) ssGSEA score of MSPC_UP geneset in each cluster. P value calculated by One-way ANOVA. Multiple comparison by Dunnett’s test. (D) Cluster 1 to 6 relative fraction by deconvolution analysis in primary prostate adenocarcinoma datasets. (E) FACS analysis of LNCaP reprogramming to mesenchymal and stem-like state. Two populations defined by EpCAM and CD104 (Integrin β4): EpCAM^HIGH^CD104^LOW^ and EpCAM^LOW^CD104^HIGH^ (F) Tumorsphere forming capacity of EpCAM ^LOW^CD104^HIGH^ population. Student’s t-test. (G) Inferred drug sensitivity of cluster 3. The extent of correlation between cytotoxic effects of each compound (data from the Cancer Therapeutics Response Portal (CTRP) v2) and cell line cluster 3 ssGSEA score. X axis: z-scored Pearson’s correlation coefficients. Y axis: coefficients rank in compounds. (H) Violin plot showing ERBB family gene expressions in cluster 1 to 6. P value calculated by One-way ANOVA. Multiple comparison by Dunnett’s test. (I) Cell proliferation (O.D.450 by CCK-8 assay) of LNCaP cells of 7 day enzalutamide treatment followed by varying concentration of recombinant human NRG1-β1 (RhNRG1-β1). P value calculated by One-way ANOVA. Multiple comparison by Dunnett’s test. (J) Western blot of phosphorylated Her2 (Y1196), Akt (S473, T308), ERK1/2 (T202 and Y204) and S6 (S235/236) and their total protein levels of LNCaP cells untreated or treated with enzalutamide or enzalutamide plus RhNRG1-β1. β-Actin was used as β loading control. (K and L) Gene expression heatmap (K) of androgen response ssGSEA score, and Western blot (L) of the AR targets and prototypical hybrid E/M markers of LNCaP cells untreated or treated with enzalutamide or enzalutamide plus RhNRG1-β1. β-Actin was used as loading control. (M) Bar graphs of AR^+/-^ and ITGB4^+/-^ populations proportion in control untreated LNCaP cells or re-LNCaP-NRG1 cells identified by coimmunofluoresence staining (representative images in Figure S4F). (N and O) Metastatic capacity of reprogrammed LNCaP cells. (N) Representative bioluminescence images (left) and normalized photon flux (middle) of mice 10 weeks after intracardiac injection of re-LNCaP-NRG1 cells. (right) Incidences (%) of macroscopic metastasis in liver and adrenal gland. (O) NRG1 RNA FISH in reprogrammed LNCaP cell liver metastasis. Costained with DAPI. (P) Enrichment plot of ERBB2/3 activation signatures in SU2C-PCF MSPC subtype versus the rest. NES = normalized enrichment score.

The changes in gene expression occurring during the pseudotime trajectory largely recapitulated the gradual shift from luminal adenocarcinoma to MSPC deduced from bulk RNA-seq data (Figure S4C). Consistently, single sample GSEA indicated that the cluster 3 LNCaP cells are enriched for the MSPC genes, including HALLMARK EMT, HEDGEHOG_SIGNALING, KRAS_SIGNALING_UP, and Hillock-like Luminal Progenitor signatures (Figures 4C and S4D). However, in contrast to MSPC PCOs, cluster 3 LNCaP cells were depleted of E2F_TARGETS, G2M_CHECKPOINT and DNA_REPAIR signatures, presumably because enzalutamide induces cell cycle arrest and inhibits the expression of DNA repair genes (Li et al., 2017) (Figure S4D). Furthermore, cluster 3 cells were enriched of genes overexpressed by embryonic diapause-like dormant pluripotent stem cells, associated with MYC inactivation and tumor cell persistence against treatment (Boroviak et al., 2015; Dhimolea et al., 2021; Scognamiglio et al., 2016). To examine the clinical relevance of the transition from cluster 5 to cluster 3, we re-examined the transcriptional profiles of the nonresponders in the MDACC neoadjuvant enzalutamide + abiraterone dataset (Efstathiou et al., 2016). Importantly, we found that the representation of the gene expression programs of clusters 5 and 1 decreased and that of clusters 4 and 2 disappeared in the resistant tumors as compared to untreated reference tumors. Conversely, the representation of the gene expression programs of cluster 3 and, to a somewhat lower degree, that of cluster 6 increased in the persistent tumors (Figure 4D). Collectively, the gene expression programs associated with cluster 3 and cluster 6 were expressed by 50% of the tumor cells in these tumors. These results suggest that primary prostate adenocarcinomas acquire the gene expression program associated with cluster 3 and cluster 6 in response to enzalutamide.

To monitor the phenotypic transformation of LNCaP cells treated with enzalutamide, we used FACS analysis. Consistent with the observation that cluster 3 cells exhibit the lowest expression of EpCAM and the highest expression of ITGB4 as compared to other clusters (Figure S4E), FACS analysis indicated that the EpCAM^HIGH^ ITGB4^LOW^ cells are gradually depleted as the EpCAM^LOW^ ITGB4^HIGH^ cells emerge and become predominant during drug treatment (Figure 4E). Notably, EpCAM^LOW^ ITGB4^HIGH^ cells isolated by FACS at either day 2 or day 7 formed a higher number of tumorspheres as compared to EpCAM^HIGH^ ITGB4^LOW^ cells, corroborating the association of cluster 3 with stemness (Figure 4F). These results indicate that profound inhibition of AR signaling reprograms prostate adenocarcinoma cells to a hybrid E/M and stemness fate in prostate adenocarcinoma cells *in vitro* and in patients’ tumors *in vivo*.

### Enzalutamide Persistent Cells are Dependent on HER2/3 Signaling and Re-enter the Cell Cycle in Response to NRG1

To examine if the hybrid E/M and stem-like persister state is associated with newly acquired dependencies, we inferred the drug sensitivities of cancer cell lines expressing cluster 3 genes by examining the Cancer Therapeutics Response Portal v2 (Supplementary Table 4) (Rees et al., 2016). Strikingly, this analysis predicted that cluster 3 cancer cells are exquisitely sensitive to HER1 and 2 kinase inhibitors (Figure 4G). Corroborating this finding, scRNAseq indicated that cluster 3 LNCaP cells express elevated levels of ERBB2 and ERBB3 and, to a smaller extent, EGFR (Figure 4H). These findings suggest that HER2/3 kinase signaling is specifically activated during enzalutamide-induced reprogramming and in MSPC patient samples.

Based on these results, we reasoned that activation of HER2/3 kinase could rescue the reprogrammed LNCaP cells from growth arrest. Indeed, physiological amounts of recombinant human neuregulin-β1 (NRG1) enabled the reprogrammed LNCaP cells to proliferate in the presence of enzalutamide (Figure 4I). Higher doses of NRG1 did not promote this process, presumably because they inhibit HER2/3 dimerization (Yarden and Pines, 2012). In contrast, EGF, which activates HER1 and HER1/2 dimers, and FGF, which activates FGF receptors, did not promote the proliferation of enzalutamide tolerant LNCaP cells (Figure S4F). The reprogrammed and rescued LNCaP cells (heretofore re-LNCaP-NRG1) exhibited robust activation of HER2 and the Ras-ERK and PI3K-mTOR pathways (Figure 4J). In addition, they maintained hybrid E/M traits and did not display signs of restoration of AR signaling as a population (Figure 4K and 4L). However, immunofluorescent staining indicated that, similar to the enzalutamide persister cells, they contained a fraction of AR^+^ ITGB4^+^ cells (Figures 4M and S4G). Parenthetically, we note that cluster 4 cells, the immediate precursor of cluster 3 cells, express AR and elevated levels of HER3 (Figure 4H) and, hence, could also potentially respond to NRG1. This supposition is consistent with the realization that microenvironmental NRG1 promotes anti-androgen resistance in AR^+^ luminal adenocarcinoma models (Zhang et al., 2020).

To examine the metastatic potential of the re-LNCaP-NRG1 cells, we conducted intracardiac injection experiments. In contrast to parental LNCaP cells, the re-LNCaP-NRG1 cells produced macroscopic metastases in the liver and, less frequently, in the adrenal glands of uncastrated mice (Figure 4N). RNA-FISH pointed to robust expression of NRG1 by normal hepatocytes and stromal cells in the tumor microenvironment but not by metastatic tumor cells (Figures 4O; arrows point to stromal cells). Notably, about 80% of metastatic tumor cells were AR negative or low, but the remainder exhibited moderate or strong nuclear accumulation of AR (Figure S4H). Since the re-LNCaP-NRG1 cells injected in mice contained a similar fraction of AR^+^ cells, we presume that these cells expanded in castrated mice in response to adrenal androgens (Mostaghel et al., 2019). These findings indicate that HER2/3 signaling rescues the enzalutamide persister cells from growth arrest and promotes their capacity to colonize metastatic sites in response to NRG1 produced by elements of the local microenvironment.

To confirm the relevance of NRG1 and HER2 signaling to human prostate cancer, we examined the SU2C-PCF dataset. GSEA demonstrated that NRG1-HER2 signaling is substantially activated in MSPC but not ARPC or NEPC in the SU2C-PCF dataset (Figure 4P). In addition, RPPA analysis of the TCGA dataset indicated that, although cluster 3 gene expression is not predominant among primary prostate cancer samples (Figure 4D), it strongly correlates with the phosphorylation and activation of HER1 or 2 kinase and components of downstream RAS-ERK, PI3K-mTOR, and JAK2-STAT3 pathways (Figure S4I, J). These observations suggest that HER2/3 signaling is activated to a substantially higher level in MSPC as compared to ARPC or NEPC.

### Inactivation of BRCA1 and E2F1 Induces Transcriptional Silencing of TP53 at the Onset of Reprogramming

To identify the oncogenic pathways involved in the transition from ARPC to MSPC, we ranked the 189 oncogenic signatures curated in the GSEA database according to their enrichment in MSPC as compared to ARPC. Intriguingly, P53_DN.V1_UP reflective of inactivation of TP53 was the top signature enriched in MSPC in the PCO-94 dataset. In addition, it ranked amongst the top 10 signatures enriched in this subtype in the SU2C-PCF, FHCRC, and UCSF datasets (Figures 5A, B and S5A). Since the *TP53* gene is not mutated at a higher rate in MSPC as compared to ARPC (Figures 2F and S2D), we considered the possibility that inhibition of AR signaling induces transcriptional silencing of wild type *TP53* in MSPC. Consistently, we found that AR activity (ssGSEA score of HALLMARK_ANDROGEN_RESPONSE) strongly correlates with the expression of *TP53* mRNA in wild type *TP53* samples from the SU2C-PCF (no mutation or deletion, n=58, Spearman r=0.54, P<0.001) and FHCRC datasets (n=72, Spearman r=0.36, P=0.002, Figure S5B). Analysis of this correlation in each subtype did not yield statistically significant results presumably because of the paucity of samples. These results suggest that *TP53* is broadly inactivated in MSPC, either as a result of genetic mutation or transcriptional silencing, the latter event possibly arising from attenuation of AR signaling.

**Figure 5.**
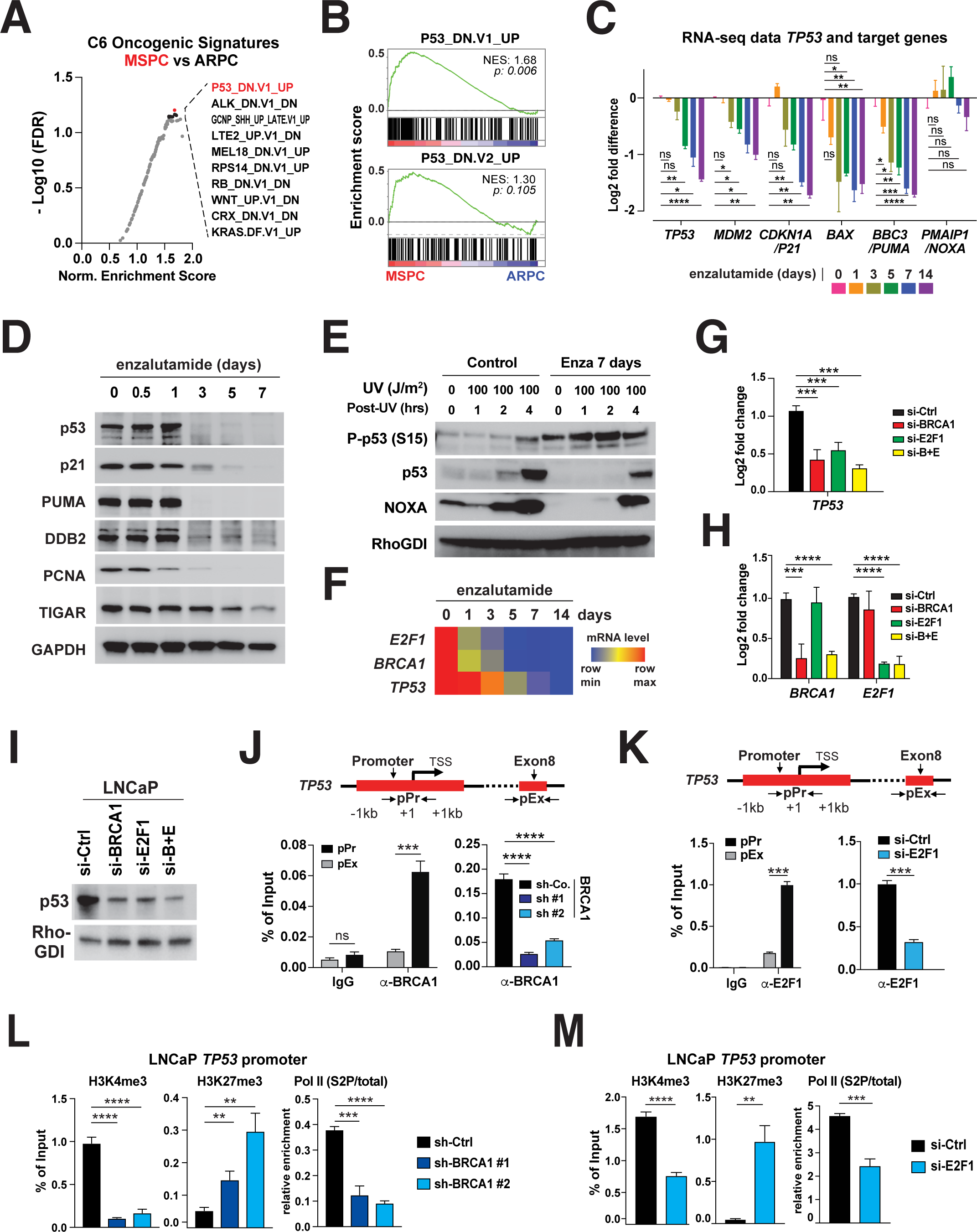
Drug-induced TP53 transcriptional inactivation mediated by BRCA1 and E2F1. (A) Top 10 Genesets enriched in MSPC vs ARPC from the PCO-94 dataset. Parental gene sets are the C6 Oncogenic Signatures (n=189) of the MSigDB Collections. (B) Enrichment plots of P53_DN.V1_UP and P53_DN.V2_UP genesets in MSPC vs ARPC from the PCO-94 dataset. (C) Log2 transformed fold differences of the TP53 and selected target genes FPKM values from RNA-seq of LNCaP cells treated with 10µM enzalutamide for 0, 1, 3, 5, 7 or 14 days. 2-Way ANOVA, multiple comparisons test by Dunnett’s method. *adjusted p<0.05; **<0.01; ***<0.001; ****<0.0001; ns = not significant. (D) P53 and its regulatory targets expression by Western Blot in LNCaP enzalutamide time-series (E) P53 and its regulatory targets expression by Western Blot in LNCaP enzalutamide time-series, combined with of UV ray treatments (by indicated time points) (F) Heatmap of ranked normalized z-scores of the *E2F1*, *BRCA1* and *TP53 mRNA* FPKM values from RNA-seq of LNCaP cells treated with 10µM enzalutamide for 0, 1, 3, 5, 7 or 14 days. (G and H) Relative levels of *BRCA1*, *E2F1* (G) and *TP53* (H) mRNA in LNCaP cells transfected with annotated siRNAs for 48hrs. GAPDH served as internal control. (I) Western blot showing p53 protein levels in LNCaP cells transfected with annotated siRNAs for 96hrs. Rho-GDI served as a loading control. **(J)** Enrichment of *BRCA1* and immunoglobulin G (IgG) control on the *TP53* gene from ChIP-qPCR in LNCaP cells (left); Occupancy of *BRCA1* on the *TP53* promoter from ChIP-qPCR in sh-control and sh-*BRCA1* LNCaP cells (right); error bars indicate mean ± SD. **p<0.01, ***p<0.005, ****p<0.001. **(K)** Enrichment of *E2F1* and immunoglobulin G (IgG) control on the *TP53* gene from ChIP-qPCR in LNCaP cells (left); Occupancy of *E2F1* on the *TP53* promoter from ChIP-qPCR in si-control and si-*E2F1* LNCaP cells (right); error bars indicate mean ± SD. **p<0.01, ***p<0.005, ****p<0.001. (L and M) Enrichment of H3K4me3, H3K27me3 and relative occupancy of RNA Pol II Ser2p on the *TP53* promoter from ChIP-qPCR in sh*BRCA1* (H) or si*E2F1* (I) LNCaP cells; error bars denote mean ± SD. **p<0.01, ***p<0.005, ****p<0.001.

To examine if AR blockade leads to transcriptional inactivation of *TP53*, we examined the expression of *TP53* and *TP53* target genes in LNCaP cells treated with enzalutamide. Strikingly, exposure to the drug caused a rapid decline in the expression of *TP53*, *MDM2*, *CDKN1A(p21)*, BAX, *BBC3(PUMA)* (Figure 5C). *TP53* mRNA and protein levels declined sharply during the first 3 days of enzalutamide treatment and more gradually during the remainder of the 14 days time course (Figure 5C, D). Additional lists of p53 downstream target genes whose expression decreased in similar degree are shown in Supplementary Figure 5C. We confirmed the inactivation of p53 by UV-induced p53 activation test and qPCR of its target genes (Figure 5E, supplementary Figure 5D). To dissect the mechanism leading to transcriptional inactivation of TP53, we identified transcription factors able to bind to the *TP53* promoter by using the ENCODE_CHIP database and selected those which were upregulated or downregulated by more than two-fold within 24 hours of treatment and did not rebound thereafter. BRCA1 and E2F1 were the only potential transcriptional regulators of TP53 which satisfied these criteria (Figure 5F). Moreover, silencing of *BRCA1* or *E2F1* caused a substantial repression of *TP53,* and simultaneous silencing of both *BRCA1* and *E2F1* exerted an even larger inhibitory effect, leading to near loss of protein expression (Figure 5G-I). These results identify BRCA1 and E2F1 as upstream regulators of TP53 in LNCaP cells.

To corroborate the hypothesis that *BRCA1* and *E2F1* directly regulate the expression of TP53, we conducted ChIP-qPCR experiments. The results indicated that both transcription factors bind directly to the *TP53* promoter in LNCaP cells (Figure 5J and 5K). Moreover, doxycycline-induced expression of shRNAs targeting *BRCA1* caused a significant decrease in the activation marks P-S2-Pol II and H3K4me3 and, reciprocally, an increase in the repression mark H3K27me3 associated with the *TP53* promoter, confirming that *BRCA1* positively regulates *TP53* (Figure 5L). To extend these results, we analyzed prostate cancer cells from *Pten*^PC/PC^ mice. Silencing of *Brca1* also deactivated and repressed the *Trp53* promoter and decreased p53 expression in these cells (Figure S5E and S5F). Enzalutamide treatment caused a similar deactivation and repression of the *Trp53* promoter, suggesting that the extent of downregulation of *Brca1* induced by pharmacological inhibition of the AR is sufficient to inactivate the *Trp53* promoter (Figure S5G). Finally, silencing of *E2F1* resulted in a decrease in the activation marks P-S2-Pol II and H3K4me3 and an increase in the repressive mark H3K27me3 at the *TP53* promoter in LNCaP cells, confirming that *E2F1* also positively regulates the *TP53* promoter (Figure 5M). These results suggest that *BRCA1* and *E2F1* jointly control the expression of *TP53* in prostate cancer cells.

### Inactivation of TP53 but not BRCA1 Drives Dedifferentiation and Acquisition of Stemness Traits

To determine whether inactivation of TP53 is sufficient to mediate reprogramming to MSPC in response to enzalutamide, we silenced TP53 in LNCaP cells. TP53 inactivation caused downregulation of ZO-1, but not E-cadherin, and induced expression of vimentin, fibronectin, and ITGB4 (Figures 6A), as observed in enzalutamide-treated cells (Figures 3C, S3C, and S3E). Moreover, silencing of TP53 induced expression of EMT and stemness signatures overlapping with those induced by enzalutamide (Figure 6B). However, in contrast to enzalutamide, silencing of TP53 did not downregulate the AR or AR signaling (Figure 6A and 6B). Functional analysis indicated that the TP53-silenced cells formed a larger number of tumorspheres in suspension and tumor organoids in 3D Matrigel as compared to control cells (Figures 6C and 6D). Intriguingly, although NRG1 did not increase the capacity of TP53-silenced cells to form tumorspheres or organoids (unpublished data), it induced them to invade *in vitro* to a substantially larger extent as compared to control cells (Figure 6E). Silencing of *Trp53* induced hybrid E/M traits and invasion in response to NRG1 without suppressing AR expression also in prostate cancer cells from *Pten*^PC/PC^ mice (Figure 6F-6H). These findings suggest that inactivation of TP53 induces reprogramming to a hybrid E/M and stem-like state and promotes invasion in response to NRG1 without downregulating AR signaling.

**Figure 6.**
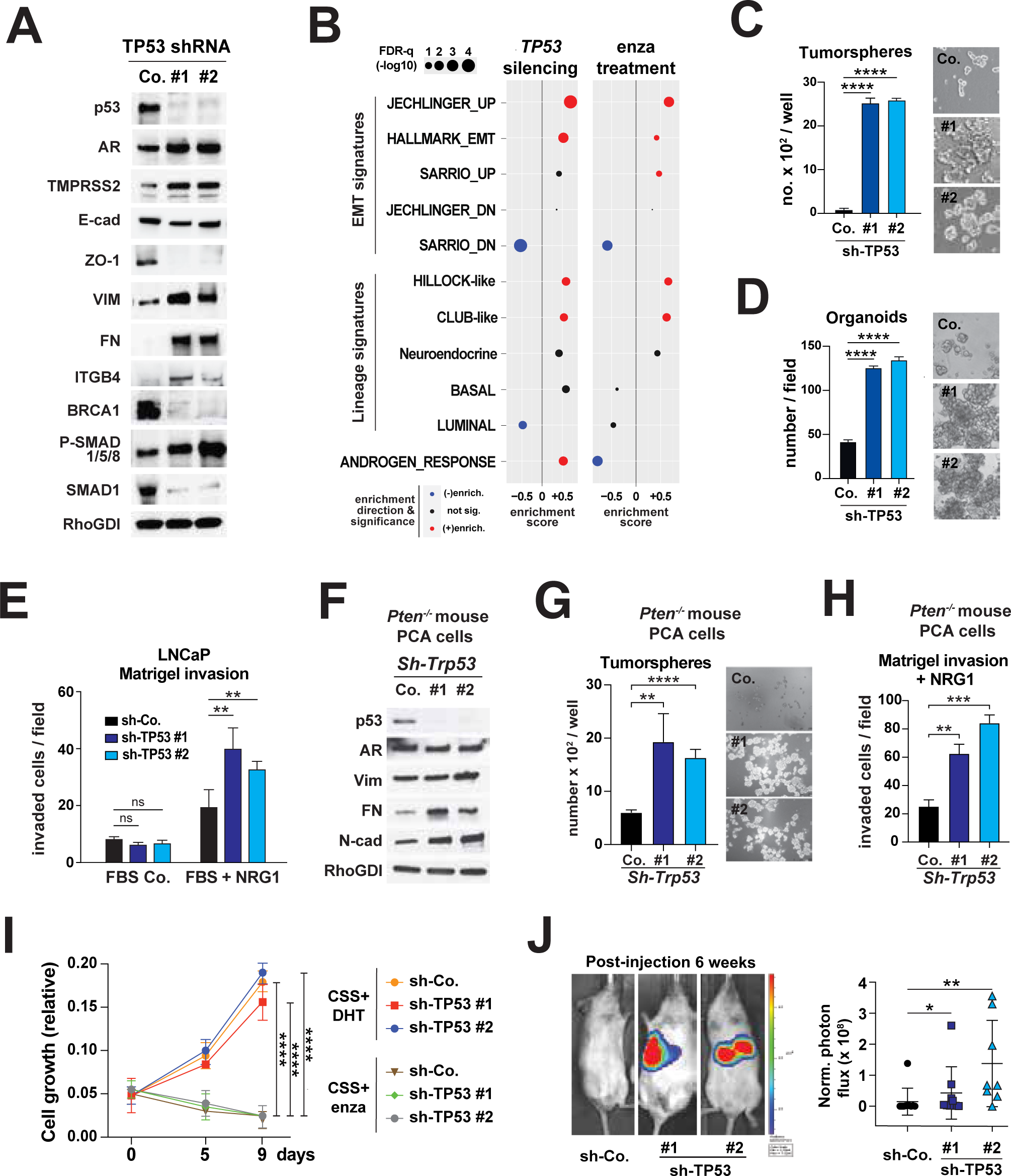
TP53 inactivation induces hybrid E/M and stem features, not anti-androgen resistance. (A) Western blot of selected AR, AR-target genes, EMT, stamness and BMP signaling markers in LNCaP cells stably transduced with the indicated hairpins. RhoGDI served as loading control. (B) ClusterProfiler dot plot showing EMT signatures enriched (top) or stemness signatures enriched (bottom) by the genes differentially expressed in control compared to *TP53* silenced condition (left), or in control compared to enzalutamide 7days treatment condition (right). (C) Quantification (left) and representative images (right) of control and *TP53*-silenced LNCaP cells subjected to sphere assay at day 10. The indicated cells were plated in triplicate in 24 well ultra-low attachment plates at a seeding density of 3,000 cells/well. Error bars, mean±SD of triplicate experiments, **** p<0.0001 two-tailed Student’s t-test. (D) Quantification (left) and representative images (right) of control and *TP53*-silenced LNCaP cells subjected to organoid formation assay. 100 –10,000 dissociated cells were plated into wells of ultra-low attachment 96 well plates. Organoid number per field (left) and diameter in micrometer (right) were counted and measured at day 10. Scale bar = 100 μm. Error bars, mean±SD of triplicate experiments, **** p<0.0001 two-tailed Student’s t-test. (E) Quantifications of *TP53*-silenced LNCaP cells subjected to matrigel invasion assay in response to NRG1. Indicated cells were plated in triplicates in 24 well Matrigel coated chambers at a seeding density of 10000 cells/well and counted 96 hrs after seeding with and without NRG as attractant. The error bars represent the SD of triplicate experiments. (F) Western blot of p53, AR, and selected EMT markers in Pten-P8^(-/-)^ cells stably transduced with the *Trp53*-indicated hairpins. Rho-GDI served as loading control. (G) Quantification (left panel) and representative images (right panel) of control and *Trp53*-silenced Pten-P8^(-/-)^ cells subjected to sphere assay at day 10. The indicated cells were plated in triplicate in 24 well ultra-low attachment plates at a seeding density of 3,000 cells/well. Error bars, mean±SD of triplicate experiments, **p<0.01, ***p<0.005, ****p<0.001 two-tailed Student t test. (H) Quantifications of *Trp53*-silenced Pten-P8^(-/-)^ cells subjected to matrigel invasion assay in response to recombinant human NRG1. Indicated cells were plated in triplicates in 24 well Matrigel coated chambers at a seeding density of 10000 cells/well and counted 96 hrs after seeding. Error bars, mean±SD of triplicate experiments, **p<0.01, ***p<0.005, ****p<0.001 two-tailed Student t test. (I) Cell proliferation assay of TP53-silenced LNCaP cells treated with CSS + DHT (upper) or CSS + Enza 10µM (lower) for the indicated time points (days). (J) Representative images (upper) and quantification of luciferase counts (lower) of male NGS mice at 6 weeks after injected i.c. with 3.0 x 10^5^ LNCaP cells expressing the indicated constructs. error bars denote mean ± SD. *p<0.01, **p<0.005.

We had shown that BRCA1 and E2F1 cooperate to induce expression of TP53 (Figure 5G-I). Notably, silencing of TP53 led to a profound downregulation of BRCA1, revealing a potential positive feedback loop, whereby TP53 function is required for expression of BRCA1 (Figure 6A). Inactivation of BRCA1 and the ensuing defect in the assembly of DNA repair foci containing HES1 and NUMB drives reprogramming of luminal breast adenocarcinoma to basal-like triple-negative breast cancer (Wang et al., 2019). To examine the possibility that inactivation of BRCA1 contributes to reprogramming to MSPC in addition to or independently from inactivation of TP53, we used doxycycline-inducible shRNAs to silence BRCA1 in LNCaP cells (Figure S6A). Stable silencing of BRCA1 did not lead to downregulation of TP53 protein, suggesting that this event may require concurrent stable silencing of E2F1 (Figure 5I). As anticipated, inactivation of BRCA1 induced broad transcriptional changes, including the downregulation of several DNA repair and cell cycle signatures (Figure S6B), and it attenuated cell proliferation (unpublished data). However, it did not induce EMT and stemness traits measured by sphere formation (Figure S6C). In fact, BRCA1 silencing led to the opposite effect in LNCaP cells as compared to mammary adenocarcinoma cells (Wang et al., 2019): it downregulated EMT and stemness signatures and upregulated mesenchymal-to-epithelial transition (MET) and stem cell differentiation signatures (Figure S6B). In addition, although silencing of TP53 restored the ability of BRCA1-silenced cells to proliferate *in vitro*, as anticipated from early studies in knock-out mice (Hakem et al., 1996), it suppressed the ability of TP53-deficient cells to form tumorspheres (Figures S6D and unpublished data). We concluded that inactivation of TP53 is sufficient to induce hybrid E/M and stem cell traits without a patent contribution from the associated deficiency in BRCA1.

Consistent with its apparent lack of effect on AR signaling (Figure 6A and 6B), inactivation of TP53 did not increase the sensitivity of LNCaP cells to androgen or decrease their sensitivity to enzalutamide, suggesting that loss of TP53 does not contribute to enzalutamide resistance (Figure 6I). To test the metastatic capacity of TP53 silenced cells, we injected them intracardially in noncastrated NSG mice and found that they generate macrometastases in the liver and adrenal glands, whereas control cells do not (Figure 6J). These findings indicate that inactivation of TP53 promotes dedifferentiation to a hybrid E/M and stem-like state and induces metastatic competency. However, it does not contribute to enzalutamide resistance.

### Inhibition of BMP Signaling Sustains AR-independent Survival and HER2/3-dependent Proliferation

Since BMP-SMAD signaling promotes the differentiation of basal stem cells in several stratified epithelia, including the prostatic epithelium (Mishina et al., 1995; Mou et al., 2016), we asked if blockade of the AR results in inhibition of BMP signaling and if the latter event contributes to the acquisition of E/M and stemness traits and enzalutamide resistance. To examine these hypotheses, we generated a signature reflective of BMP receptor inactivation in LNCaP cells by selecting the genes modulated by the selective BMP-R1 inhibitor LDN193189 (Hao et al., 2010) in both control and TP53-silenced LNCaP cells (Figure 7A). Notably, ssGSEA indicated that the LDN193189_RESPONSE score was inversely correlated with the HALLMARK_ANDROGEN_RESPONSE score in metastatic samples from the FHCRC, SU2C-PCF, and UCSF datasets (Figures 7B and S7A). Treatment of LNCaP cells with enzalutamide resulted in rapid and coordinated inhibition of AR and BMP signaling, suggesting that the transcriptional activity of the AR sustains activation of the BMP pathway (Figure 7C). Further analysis indicated that enzalutamide downregulates the expression of the two type I BMP receptors BMP-R1A and B, the coreceptors RGMA and B, and the BMP-responsive SMADs and it suppresses the phosphorylation and activation of BMP-responsive SMADs (Figures 7D, 7E, and S7B; note that the *SMAD9* gene encodes SMAD8). Stable silencing of the AR induced similar changes in the expression of BMP pathway components (Figure S7C, GSE22483). Conversely, the synthetic androgen R1881 induced the expressions of *BMPR1A, BMR1B, NEO1, SMAD1, SMAD5, and SMAD9* (Figure S7D). Notably, *BMPR1B* was the BMP pathway component most profoundly induced by AR signaling, consistent with its identification as the top gene expressed in ARPC (Figure 2B). These results indicate that AR signaling controls, either directly or indirectly, the expression of several BMP pathway components and thereby sustains BMP signaling, explaining why blockade of the AR results in inhibition of BMP signaling.

**Figure 7.**
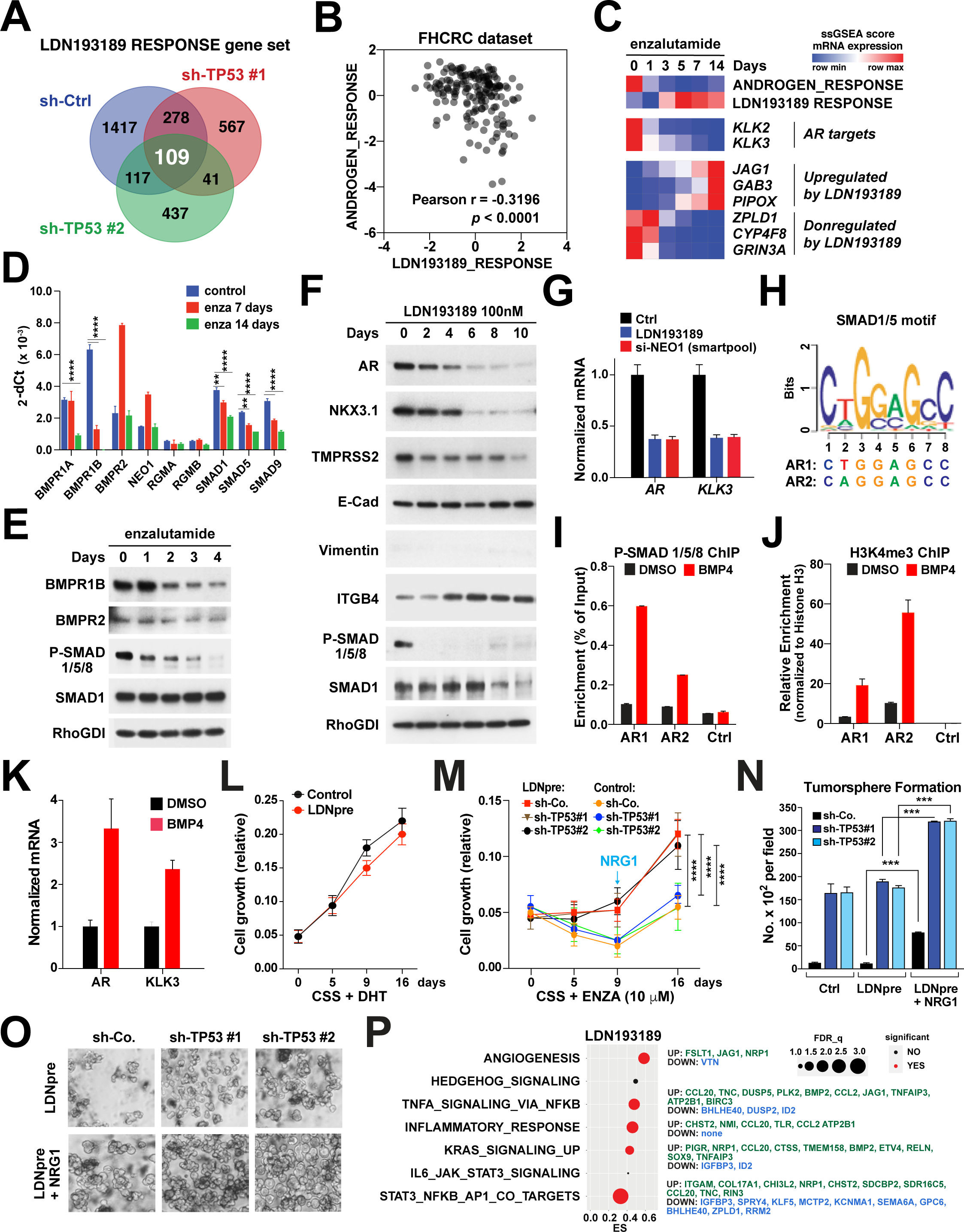
Inactivation of BMP-Smad signaling induces anti-androgren resistance. (A) Venn diagram of differently expressed genes in LNCaP shCtrl and shTP53 #1 and #2 cells treated with or without LDN193189 100nM for 8 days. The overlapping 109 genes were designated as “LDN193189 RESPONSE” geneset. (B) Scatter plot showing correlation of ssGSEA scores of the HALLMARK_ANDROGEN_RESPONSE and LDN193189 response in the FHCRC dataset. (C) Heatmap of average ssGSEA scores of the HALLMARK_ANDROGEN_RESPONSE and LDN193189 response in LNCaP cells treated with enzalutamide 10µM for 0 to 14 days. Expression levels of genes representing each geneset is also shown. Three replicates per condition. (D) mRNA expressions (normalized to 18S) of LNCaP cells treated with enzalutamide (enza) 10µM for 0, 7 and 14 days. ** p<0.01, **** p<0.001. Student’s t*-*test. (E) Immunoblotting of BMP signaling components in LNCaP cells treated with enzalutamide for 0-4 days. (F) Immunoblotting of AR, AR targets and EMT/Stemness markers in LNCaP cells treated with LDN193189 100nM for 0-10 days. (G) AR and KLK3 mRNA expressions measured by qPCR in serum starved LNCaP cells either treated with LDN193189 50nM or NEO1 siRNA (SMART pool). (H) Schematic view of SMAD1/5 binding motif on androgen receptor (AR) promoter (AR1 and AR2), used in panel 6I and 6J. (I and J) ChIP analysis. LNCaP cells grown in androgen depleted media with 2% charcoal stripped serum were treated for 8 hours with BMP4 100 ng/mL or DMSO. Cells were crosslinked and processed for ChIP using phosphorylated SMAD1/5/8 antibody (left) and H3K4me3 antibody (right). Real-time PCR quantification of immunoprecipitated 2 distal sites in AR promoter that both contain SMAD1/5 motif (refer to panel 6H) and negative control site in AR exon2 are shown (Mean ± SD, n=3). (K) AR and KLK3 mRNA expressions measured by qPCR in serum starved LNCaP cells either treated for 24hrs with BMP4 100 ng/mL or DMSO as control. (L) Relative cell growth of LNCAP control or LDN193189 (pretreated for 8 days) in the conditions of CSS+DHT for additional 16 days. (M) Relative cell growth of LnCAP shCo., shTP53 #1, shTP53 #2 cells grown in control or LDN193189 (pretreated for 8 days) and in the conditions of CSS+ENZA (10µM) for additional 16 days. (N and O) Quantification (N) and representative images (O) of control and TP53-silenced LNCaP cells subjected to sphere assay at day 10. The indicated cells were pretreated with or without LDN193189 100nM and/or rhNRG1 for 8 days. Then the cells were plated in triplicate in 24 well ultra-low attachment plates at a seeding density of 3,000 cells/well. Error bars, mean±SD of triplicate experiments, **** p<0.0001 two-tailed Student’s t-test. (P) Clusterprofiler dot plot showing NF-κB and STAT3 transcription factor signatures enriched using gene set enrichment analysis (GSEA) for genes differentially expressed in control compared to LDN193189 treatment condition. X-axis title “ES” represents the GSEA enrichment score. Y-axis represents the name of the signatures. Dot size represents the -log10 (FDR_q_value + 0.001). Dot color represents the significance (left); For each significant signature, the top differential expression genes including upregulated and downregulated genes are listed (right).

To investigate the consequences of inhibition of BMP signaling, we treated LNCaP cells with LDN193189 over a period of 10 days. Although the compound did not inhibit cell proliferation (Figure S7E), it suppressed the expression of AR, KLK3, TMPRSS2, and the luminal lineage transcription factor NKX3.1 and upregulated the expression of the stem cell marker ITGB4. However, it did not promote the expression of vimentin (Figure 7F and 7G). Genetic or antibody-mediated inactivation of the essential BMP coreceptor NEO1 inhibited BMP signaling and downregulated the AR and the AR target gene KLK3 as efficiently as LDN193189 (Figures 7G, S7F-H). GSEA confirmed that LDN193189 inhibits AR signaling without inducing mesenchymal traits and revealed that the compound induces an enrichment of club-like luminal progenitor and neuroendocrine signatures (Figure S7I and S7J). In addition, functional assays indicated that an 8 day pretreatment with LDN193189 does not increase the capacity of TP53-silenced LNCaP cells to form tumorspheres (Figures 7N). These results indicate that inhibition of the BMP-SMAD pathway downregulates AR and AR signaling and provokes partial dedifferentiation and lineage infidelity, without inducing patent mesenchymal traits or stemness.

To identify the mechanism leading to downregulation of the AR, we inspected the promoter of the AR and found two canonical SMAD1/5 binding motifs (Figure 7H). ChIP qPCR indicated that BMP promotes binding of activated P-SMAD1/5/8 to both motifs and simultaneously enriches the activation mark H3K4me3 in the surrounding chromatin (Figure 7I and 7J). In addition, BMP promoted transcription of *AR* and *KLK3* (Figure 7K). These findings indicate that BMP signaling induces binding of the BMP-responsive P-SMADs to the promoter of the *AR* and directly controls transcription of the gene. These results suggest that enzalutamide downregulates the expression of the AR at least in part because it inhibits BMP signaling.

Since LDN193189 inhibited AR expression and signaling but did not inhibit proliferation, we surmised that it activated alternative pathways for survival and proliferation. We therefore asked if inhibition of BMP signaling contributes to enzalutamide resistance. LNCaP cells pretreated with LDN193189 grew in response to androgen almost as efficiently as control cells, suggesting that they remain androgen responsive for proliferation in spite of diminished levels of AR (Figure 7L). Intriguingly, however, the cells pretreated with LDN193189 survived in the presence of doses of enzalutamide up to 25 μM, whereas control cells did not (Figure S7M). Furthermore, the cells pretreated with LDN193189 and surviving in the presence of 10 μM enzalutamide started to proliferate in response to NRG1 and robustly expanded in the continuous presence of the drug (Figure 7M). Silencing of TP53 did not improve the performance of LDN193189-treated LNCaP cells in the presence of enzalutamide, confirming that inactivation of TP53 does not contribute to antiandrogen inhibitor resistance (Figures 7M and S7M). These findings indicate that inhibition of BMP signaling promotes cell survival in the face of enzalutamide and enables subsequent expansion in response to NRG1.

We next examined the effect of inhibition of BMP signaling on stemness traits. Pretreatment with LDN193189 enhanced the capacity of control and TP53-silenced cells to form tumorspheres in the presence but not the absence of NRG1 (Figure 7N and 7O). Notably, the TP53-silenced cells formed a significantly higher number of spheres if they were pretreated with LDN193189 and then exposed to NRG1 during the assay as compared to those that had not been pretreated, suggesting that inhibition of BMP signaling and inactivation of TP53 cooperate to sustain self-renewal in response to NRG1 (Figure 7N and 7O). Furthermore, the TP53-silenced LNCaP cells generated more organoids than control cells when pretreated with LDN193189 and cultured in androgen-depleted medium. NRG1 further promoted organoids formation both in TP53-silenced and control cells that had been pretreated with LDN193189, suggesting that inhibition of BMP signaling and inactivation of TP53 cooperate to sustain proliferation and aberrant luminal differentiation in the absence of exogenous androgen (Figure S7K and S7L). Therefore, inhibition of BMP signaling promotes anti-androgen resistant survival and enhances NRG1-driven stemness.

These findings raised the issue of mechanism. How does inhibition of the BMP-SMAD pathway exert these effects? GSEA indicated that KRAS_SIGNALING_UP and inflammatory signatures dependent on NF-κB or JAK-STAT3 signaling were amongst the top 10 Hallmark signatures induced by LDN193189 or enzalutamide (Figure 7P and S7N). The top genes within the signatures upregulated by LDN193189 included proteins potentially involved in prostate cancer therapy resistance and metastasis, such as the Notch ligand JAG1 (Domingo-Domenech et al., 2012), the lineage transcription factor SOX9 (Nouri et al., 2020), the inhibitor of apoptosis protein BIRC3 (Silke and Vucic, 2014), and the cytokine CCL2 (Su et al., 2019). In addition, CCL20 has been implicated in chemotherapy resistance in breast cancer (Chen et al., 2018). Mechanistically, BIRC3 activates NF-κB (Silke and Vucic, 2014) and CCL2 activates JAK-STAT3 signaling (Izumi et al., 2013). Together with AP1, which is induced by RAS signaling, NF-κB and STAT3 coordinate regulatory networks involved in chronic inflammation (Taniguchi and Karin, 2018). Consistently, LDN193189 also induced phosphorylation of the STAT3 protein and increased the expression of the STAT3_NFKB_AP1_CO_TARGETS signature (Figure 7P and S7O), which constitutes a core inflammatory signature in several cancer types (Ji et al., 2019). These observations suggest the possibility that inhibition of BMP signaling promotes anti-androgen resistance through the coordinated action of several target genes and the chronic inflammatory network that they control.

### Neratinib-based Combinations Exert Preclinical Efficacy in MSPC Xenograft Models

MSPC is characterized by prominent activation of RAS and PI3K signaling (Fig. 1C). In a fraction of DNPC cases, the activation of these signaling pathways is presumed to arise from upstream activation of FGFRs (Bluemn et al., 2017). To examine the potential sensitivity of MSPC to inhibition of HER2/3, we clustered the MSPC samples from the SU2C-PCF dataset according to their inferred sensitivity to either HER or FGFR inhibitors. We found that about half of the MSPC samples are predicted to be sensitive to the HER1/2 inhibitor lapatinib and the pan-HER inhibitor neratinib, but not to the FGFR inhibitors AZD4547 and nintedanib, and vice versa (Figure 8A). Indeed, correlation analysis demonstrated an inverse correlation between the predicted sensitivity of MSPC cases to the two types of inhibitors. Moreover, the anticipated sensitivity of these cases to HER inhibitors was positively correlated with elevated levels of expression of HER1-3 (Figures 8A). Examination of the FHCRC and UCSF datasets yielded similar results (Figures S8A). These observations raise the possibility that a substantial fraction of MSPC cases may be sensitive to pan-HER inhibitors, such as neratinib. It also suggests that AR inhibitor-induced prostate cancer cell reprogramming provides a window of opportunity for combination therapy with HER2/3 inhibitors or those targeting the downstream PI3K-AKT-TOR signaling and RAS-RAF-ERK signaling (Figure 8B).

**Figure 8.**
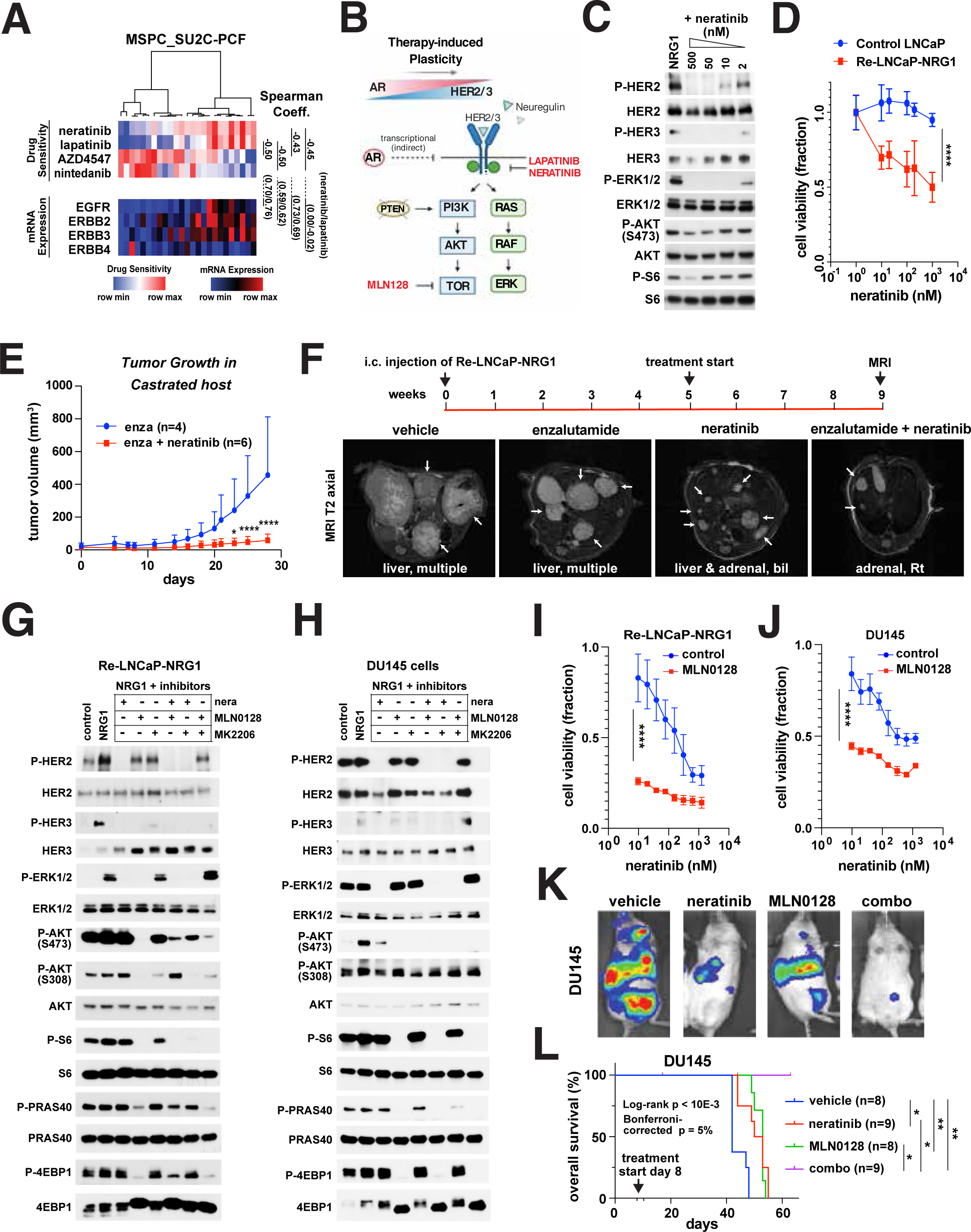
ERBB2/3 and AR inhibitors combination to block the rise of MSPC. (A) Predicted Drug sensitivity heatmap of MSPC samples from the SU2C-PCF dataset. mRNA expressions of ERBB1-4 are shown together. Spearman correlation coefficients between neratinib/lapatinib and FGF inhibitors AZD4547/nintedanib sensitivity scores or the mRNA expressions are shown (right). (B) Schematic representation of drug-induced lineage plasticity from AR-dependent to HER2/3-dependent state of prostate cancer cell and adjuvant therapies targeting HER2/3 and the downstream pathways. (C) Immunoblotting of HER2/3, ERK1/2, AKT and S6 protein phosphorylation upon different neratinib dosage in re-LNCaP-NRG1 cells. (D) Cell viability dose response curve of LNCaP control and re-LNCaP-NRG1 cells to neratinib. (E) Subcutaneous tumor growth in castrated male NSG mice of re-LNCaP-NRG1 cells treated with daily enzalutamide 10mg/kg and/or neratinib 40mg/kg. (F) Representative MRI images of re-LNCaP-NRG1 metastatic tumors in castrated male NSG mice treated with daily enzalutamide 10mg/kg and/or neratinib 40mg/kg for 4 weeks. (G and H) Immunoblotting of HER2/3, ERK1/2, AKT and mTOR kinase pathway protein phosphorylation upon MLN0128 100nM or MK2206 0.5µM short term treatment (4 hours) in re-LNCaP-NRG1 cells (F) or DU145 cells (G). (I and J) Cell viability dose response curves of re-LNCaP-NRG1 cells (H) or DU145 cells(I) upon MLN1028 50nM and/or varying dose of neratinib. (K) Representative bioluminescence images of DU145 metastatic tumors in castrated male NSG mice treated with daily neratinib 40mg/kg and/or MLN0128 0.3mg/kg for 4 weeks. (L) Kaplan-Meier survival curves of mice of DU145 cells intracardiac injection and treated with daily neratinib 40mg/kg and/or MLN0128 0.3mg/kg.

To study the preclinical efficacy of neratinib, we first examined its ability to inhibit HER2/3 signaling in LNCaP cells reprogrammed with enzalutamide. Nanomolar concentrations of neratinib (10-50 nM) efficiently blocked NRG1-driven activation of HER2/3 and ERK in these cells (Figure 8C). However, they did not suppress activation of AKT and mTOR, as anticipated from the *PTEN* mutant status of these cells (Li et al., 1997). Furthermore, neratinib and lapatinib inhibited the proliferation of re-LNCaP-NRG1 cells to a substantially larger extent as compared to that of control LNCaP cells (Figures 8D and S8B). In contrast, the mTOR inhibitor MLN0128, pan-FGFR inhibitor AZD4547 or EGFR-specific inhibitor erlotinib did not produce such a differential effect. AZD4547 and erlotinib did not demonstrate discernable activity at doses lower than 1 μM in both types of cells and MLN0128 efficiently suppressed the proliferation of both types of cells at 100 nM (Figure S8B). These results confirm the dependency of re-LNCaP cells on HER2/3 signaling.

To test the potential efficacy of neratinib *in vivo*, we first conducted primary tumorigenesis experiments. The results revealed that, in spite of its inability to inhibit PI3K signaling, neratinib suppresses the capacity of re-LNCaP-NRG1 cells to produce subcutaneous tumors in castrated and enzalutamide-treated mice (Figure 8E). Since we had observed a certain degree of restoration of AR signaling in the metastases seeded by reprogrammed LNCaP cells (Figure S4G), we tested the preclinical efficacy of neratinib in combination with enzalutamide in the metastatic setting. Castrated mice were inoculated intracardially with re-LNCaP-NRG1 cells and treated with neratinib, enzalutamide, or the combination starting at 5 weeks. Notably, the combination reduced the size of individual metastases and overall metastatic burden to a significant extent as compared to vehicle control, whereas neither drug displayed statistically significant overall efficacy as a single agent (Figure 8F and S8C). Although no treatment regimen reduced the total number of metastases, enzalutamide and the combination reduced the size of individual metastases and metastatic burden, suggesting that tumor cells dependent on AR signaling contribute to metastatic expansion *in vivo* (Figure 8F and S8C). These results suggest that neratinib may increase the efficacy of enzalutamide in ARPC and mixed ARPC/MSPC metastases.

Given the frequent inactivation of *PTEN* in M-CRPC, we asked if inhibition of AKT or mTOR kinase increases the efficacy of neratinib in re-LNCaP-NRG1 cells and DU145 cells, both of which are classified as MSPC. In preliminary dosing experiments, the AKT inhibitor MKK2206 blocked AKT phosphorylation at S473 at 0.5-1 μM and caused a paradoxical overactivation of ERK in re-LNCaP-NRG1 cells (Figure S8D). We attribute this activation of ERK to the release of the negative feedback that AKT exerts on receptor tyrosine kinase expression and signaling (Chandarlapaty et al., 2011). In contrast, the mTOR kinase inhibitors MLN0128 and AZD8055 blocked phosphorylation of S6 and AKT at Ser 473 at similarly low nanomolar concentrations (Figure S8E).

Having identified optimal concentrations of the drugs, we conducted a detailed analysis of the effect of neratinib in combination with either MKK2206 or MLN0128 in both re-LNCaP-NRG1 cells and DU145 cells. As a single agent, the mTOR kinase inhibitor MLN0128 inhibited the activation of AKT (measured by phosphorylation of its target PRAS40) and the activation of mTOR (measured by phosphorylation of S6 and 4EBP1) more profoundly as compared to the AKT inhibitor MK2206 (Figure 8G and 8H). Moreover, MLN0128 interfered with the activation of ERK in reprogrammed and rescued LNCaP but not in DU145 cells. Since this effect was limited to the former cells and was followed at 1 and 2 days by an overactivation of ERK (Figure S8F), we did not investigate it further. Importantly, the neratinib and MLN0128 combination was superior to the other combinations in effectively and durably blocking the activation of ERK, AKT, and mTOR in both types of cells (Figures 8G, 8H and S8F). These findings indicate that neratinib and MLN0128 profoundly inhibit mitogenic signaling through both the RAS-ERK and PI3K-mTOR pathways in *PTEN* mutant MSPC cells.

To examine the capacity of neratinib in combination with MLN0128 to inhibit proliferation, we kept MLN0128 at 50 nM and titrated neratinib. Notably, nanomolar doses of neratinib substantially increased the capacity of 50 nM MLN0128 to inhibit the proliferation of re-LNCaP-NRG1 and DU145 cells (Figure 8I and 8J). Neratinib modestly increased the sensitivity of LNCaP control cells to MLN0128 (Figure S8G). In reciprocal experiments, nanomolar doses of MLN0128 increased the capacity of neratinib to inhibit the proliferation of DU145 cells (Figure S8H). Since the DU145 cells do not express the AR and possess stable MSPC traits, we used these cells to test the preclinical efficacy of neratinib in combination with MLN0128. We injected the tumor cells intracardially in castrated mice and commenced drug treatments at day 8, when tumor cells are estimated to have already seeded target organs and resisted initial attrition (Giancotti, 2013). Intriguingly, neratinib singly and in combination with MLN0128 effectively reduced metastatic burden (Figures 8K and S8I). Interestingly, the combination reduced metastatic burden more uniformly as compared to neratinib alone and increased survival to a larger extent as compared to each single agent (Figure 8L and S8J). These results document the preclinical efficacy of neratinib, both alone and in combination with MLN0128, in MSPC.

## Discussion

Understanding the biology and designing effective therapies for M-CRPC remains a formidable challenge, given the significant inter and intratumoral genetic heterogeneity of advanced stage disease (Boutros et al., 2015; Kishan et al., 2020; Kumar et al., 2016; Quigley et al., 2018; Robinson et al., 2015). Our results indicate that M-CRPC can be stratified according to three intrinsic transcriptional subtypes: ARPC, MSPC, and NEPC, which largely overlap with the AR^+^ (ARPC), AR^-^/NE^-^ (DNPC), and NE^+^ (NEPC) subtypes previously defined using restricted gene sets based on combinations of immunohistochemical markers (Bluemn et al., 2017). Notably, the gene expression programs of ARPC and NEPC resemble those of normal luminal and neuroendocrine cells, respectively. In contrast and unexpectedly, MSPC is enriched with luminal progenitor and EMT signatures (Henry et al., 2018; Sackmann Sala et al., 2017; Smith et al., 2015; Wang et al., 2009). Furthermore, whereas ARPC exhibits transcriptional similarity to luminal breast cancer, MSPC resembles basal-like breast cancer (Perou et al., 2000; Sorlie et al., 2001); and MSPC and NEPC share transcriptional characteristics with the two nonclassical subtypes of Glioblastoma Multiforme (GBM): mesenchymal and proneural (Verhaak et al., 2010). Since the transcriptional profiles of the three subtypes of M-CRPC mirror developmental programs of the normal gland that may be shared in other organs, we presume that they have arisen from the effect of oncogenic transformation on the epigenetic landscape of the cell of origin, as hypothesized for the intrinsic subtypes of other cancer types (Gupta et al., 2019).

Several mechanisms link mesenchymal and stemness programs, such as those active in MSPC, to immune evasion and immunosuppression (Dongre and Weinberg, 2019; Naik et al., 2018). Recent studies have shown that the Polycomb Repressor Complex 1 (PRC1) is overactive in DNPC and promotes metastasis by coordinating stemness and immunosuppression (Su et al., 2019). PRC1 exerts this effect by inducing expression of CCL2, which enhances self-renewal and promotes recruitment of M2-like Myeloid Derived Suppressor Cells (MDSCs). Accordingly, agents that target PRC1 or MDSCs substantially improve the efficacy of double immune checkpoint therapy (ICT) in models of advanced metastatic prostate cancer (Lu et al., 2017; Su et al., 2019). In agreement with the overlap of MSPC with DNPC, MSPC exhibits a higher PRC1 activity and a higher number of M2-like MDSCs and dysfunctional T cells as compared to ARPC and NEPC. Moreover, it is enriched for the metastatic melanoma anti-PD-1 nonresponders gene expression signature (Hugo et al., 2016). These results indicate that MSPC may be particularly refractory to ICT because of high PRC1 activity and infiltration of MDSCs. They further suggest that agents targeting MDSCs should be tested specifically in this subtype.

Leveraging the discriminatory power of intrinsic tumor cell profiling and deconvolution analysis, we found a remarkable and unanticipated degree of intratumoral transcriptional heterogeneity in M-CRPC. Although most PDXs, organoids, and patient metastases consist predominantly of tumor cells of one intrinsic subtype, a substantial fraction contains an admixture of ARPC and MSPC cells. In fact, the large majority of M-CRPC samples can be classified as ARPC, ARPC/MSPC, or MSPC, with the mixed tumors ranging in composition from almost pure ARPC to almost pure MSPC. Similarly, it has been proposed that many GBMs are comprised of admixtures of subtypes reflective of distinct cellular lineages (Neftel et al., 2019). Intriguingly, ARPC and MSPC harbor a similar repertoire of oncogenic mutations, including AR mutations and amplifications, but the proportion of mixed ARPC/MSPC and pure MSPC cases has increased substantially in recent datasets (to 20-21% and 26-44%, respectively), possibly reflecting the widespread use of second-generation AR inhibitors over the last decade. In contrast, the intrinsic subtype of NEPC occurs infrequently and is almost invariably associated with small cell histology and concurrent mutation of *TP53* and deletion of *RB1*.

Transcriptional profiling indicated that the large majority of localized primary adenocarcinomas can be categorized as ARPC or mixed ARPC/MSPC, and the infrequent occurrence of MSPC correlates with elevated pathological grade and reduced time to progression and death, suggesting that MSPC may arise at the primary site as an aggressive variant. We presume that previous attempts to transcriptionally categorize primary prostate adenocarcinomas may have failed to detect MSPC because of its infrequency (5-11%) (Cancer Genome Atlas Research, 2015; You et al., 2016; Zhao et al., 2017). In striking contrast, analysis of clinical samples from a neoadjuvant trial with abiraterone and enzalutamide indicated that the large majority of primary tumors, which did not respond to treatment, consists of MSPC, suggesting that ARPC morphs into MSPC in response to AR blockade. In addition, recent studies have shown that the nonresponders to an enzalutamide trial share a transcriptional program of low AR activity and stemness reminiscent of the one described here, but no distinctive oncogenic mutation (Alumkal et al., 2020). These findings suggest that MSPC constitutes a form of adaptive resistance to second-generation AR inhibitors and suggest that agents that exhibit activity against MSPC may improve the efficacy of such inhibitors in ARPC or mixed ARPC/MSPC cases.

Remarkably, exposure to enzalutamide and the ensuing blockade of the AR induced AR-dependent prostate adenocarcinoma cells to acquire hybrid E/M traits and become tolerant to the drug in a matter of days in the absence of proliferation. Although largely quiescent, the persister tumor cells formed a large number of tumorspheres in suspension culture and exhibited tumor initiating capacity in castrated mice, consistent with the observation that the hybrid E/M state is associated with increased stemness (Bierie et al., 2017; Pastushenko et al., 2018). Integrated RNA-seq and ATAC-seq followed by ChIP-seq provided evidence that the LNCaP cells acquire mesenchymal and stem-like traits and become tolerant to enzalutamide through epigenetic reprogramming, as it has been proposed for melanoma cells exposed to vemurafenib (Shaffer et al., 2018; Sun et al., 2014). During this process, the enhancers and promoters driving the AR and luminal differentiation program were deactivated and those controlling EMT and stemness genes became active. Further analysis revealed that the MSPC transcriptional program is dominated by the oncogenic TFs ETS1 and NFκB1/2, which have been previously linked to overproliferation, EMT, and immunosuppression (Massague, 2004; Taniguchi and Karin, 2018). Finally, LNCaP cells treated *in vitro* with enzalutamide transitioned from the ARPC to the MSPC transcriptional subtype, which we had defined in metastatic samples, validating the model and illustrating the pathological relevance of the findings. These findings indicate that enzalutamide can induce reprogramming and adaptive resistance very rapidly.

Single cell analysis of head and neck tumors has indicated that hybrid E/M tumor cells are localized at the invasive edges and generated at least in part in response to local cues (Puram et al., 2017). A similar analysis of mammary and skin tumors arising in mouse models has linked the hybrid E/M transcriptional state to increased invasion and metastasis (Pastushenko et al., 2018). In addition to corroborating the epigenetic and transcriptional trajectory of LNCaP cells undergoing reprogramming *in vitro*, single cell RNA-seq and pseudotime analysis revealed that the majority of tumor cells harboring hybrid E/M traits (cluster 3) were potentially sensitive to a variety of HER kinase inhibitors. In agreement with the observation that enzalutamide can induce expression of HER2/3 (Shiota et al., 2015), the reprogrammed LNCaP cells exhibited elevated levels of HER2/3 and, in response to NRG1, activated ERK and AKT and re-entered the cell cycle. Importantly, upon injection into castrated mice, these cells generated liver and adrenal gland metastases, indicating that dedifferentiation to a hybrid E/M and stem-like state is linked to metastatic capacity in prostate cancer. We infer that the metastatic expansion of reprogrammed cells is fueled by NRG produced by stromal cells and adjacent normal epithelial cells, which we documented with FISH analysis in mouse models. The existence of such a crosstalk suggests that the reprogrammed cells have hijacked a stromal dependency recently identified in luminal progenitors in the normal prostate (Karthaus et al., 2020). In agreement with the potential clinical significance of our findings, RPPA and GSEA demonstrated that HER2/3 signaling is significantly activated in MSPC/cluster 3 patient samples. In studies complementary to ours, it has been shown that microenvironment-derived NRG1 promotes antiandrogen resistance in ARPC models (Zhang et al., 2020). It is possible that the efficacy of combinations of HER2/3 and AR inhibitors in these models arises at least in part from AR blockade-induced reprogramming to MSPC.

Notably, we found that *TP53* is broadly inactivated in MSPC as a consequence of either mutation or transcriptional silencing. Combined *TP53* and *RB1* mutations are instead prevalent in NEPC, consistent with their inferred function in transdifferentiation to the neuroendocrine fate (Ku et al., 2017; Mu et al., 2017; Zou et al., 2017). By modeling the transition from ARPC to MSPC *in vitro*, we found that enzalutamide rapidly suppresses the expression of E2F1 and BRCA1 and thereby profoundly downregulates TP53 and TP53 target genes in *PTEN* mutant cells. Biochemical and functional analysis revealed that E2F1 and BRCA1 coordinately bind to the TP53 promoter and induce TP53 expression in control cells (Figure S8K). This indicates that E2F1 not only increases TP53 stability by inducing expression of ARF (Kastenhuber and Lowe, 2017) but also directly induces expression of TP53 by binding to canonical E2F1 sites in the TP53 promoter. In contrast, BRCA1 is not a sequence specific TF but can regulate gene expression either by promoting chromatin remodeling or by binding to sequence specific TFs (Silver and Livingston, 2012). We hypothesize that BRCA1 cooperates with E2F1 at the TP53 promoter through either or both of these mechanisms (Figure S8K). In addition, it has been proposed that BRCA1 directs the expression of a subset of TP53 target genes by binding to TP53 at their promoters (Mullan et al., 2006). These results suggest that E2F1 and BRCA1 induce TP53 by multiple mechanisms, potentially explaining why overactivation of TP53 is limited by strong feedback mechanisms, which inhibit E2F1 and hence alleviates overproliferation and oncogenic stress (Kastenhuber and Lowe, 2017). Although we have not examined the mechanism through which AR signaling induces expression of BRCA1, we postulate that E2F1-driven overproliferation causes replication stress and thereby induces and activates BRCA1. AR blockade reverses this process, enabling the accumulation of DNA damage, as observed previously (Li et al., 2017). Irrespective of mechanism, our study suggests that treatment with enzalutamide and presumably other AR inhibitors induces growth arrest and apoptosis of a large fraction of prostate cancer cells, but renders the remainder functionally deficient of BRCA1 and TP53 activity.

Prior studies have firmly implicated the inactivation of BRCA1 in lineage plasticity in breast cancer. Inactivation of BRCA1 commonly occurs in luminal progenitors but it leads to the emergence of basal-like Triple-Negative Breast Cancer (TNBC) (Turner and Reis, 2006). Recently, it was shown that BRCA1 stabilizes the differentiated state of primary mammary epithelial cells by promoting interstrand crosslink DNA repair (Wang et al., 2016). Intriguingly, BRCA1 participates in this process in association with NUMB1 and HES1, and disruption of the complex between these proteins induces persistent DNA damage and transdifferentiation to basal-like TNBC (Wang et al., 2019). In contrast, we found that, although BRCA1 is transcriptionally downregulated at the onset of reprogramming in prostate adenocarcinoma cells, it is the subsequent inactivation of TP53 that induces hybrid E/M and stem-like traits. In fact, silencing of BRCA1 suppressed the capacity of TP53-silenced prostate cancer cells to manifest these traits. Ostensibly, the signaling and transcriptional networks controlled by BRCA1 and TP53 are arranged in distinctive manners in breast and prostate epithelial cells. It is also of significance that silencing of TP53 alone was not sufficient to suppress the expression of the AR or promote enzalutamide resistance. Accordingly, the TP53-silenced cells were able to generate metastasis but only in noncastrated mice.

BMP signaling restricts the expansion of the basal compartment of the prostate and promotes luminal differentiation and AR expression (Mou et al., 2016; Omori et al., 2014). We found that canonical BMP signaling controls the expression of the AR and, conversely, the AR controls the expression of BMP pathway components and hence BMP signaling, in prostate adenocarcinoma (Figure S8K). Consistent with this model, AR blockade induced inactivation of BMP signaling and decreased AR expression and signaling in LNCaP cells. However, inhibition of BMP signaling did not promote the expression of hybrid E/M traits, presumably because it did not result in a complete suppression of AR signaling. Notably, LNCaP cells that were pretreated with the BMP-R1 inhibitor LDN193189 resisted treatment with enzalutamide, and concurrent inactivation of TP53 enabled them to expand *in vitro* and form tumorspheres and organoids in response to NRG1. Whether BMP signaling promotes anti-androgen resistance through the coordinated action of the target genes identified here awaits further studies. These findings reveal an unanticipated function of inhibition of BMP signaling in anti-androgen resistance.

Although androgen deprivation elevates the expression of HER2 in prostate cancer and HER2 signaling promotes AR stability and DNA binding (Mellinghoff et al., 2004), lapatinib or trastuzumab as monotherapy or in combination with chemotherapy or AR signaling inhibitors, such as dutasteride or ketoconazole, have not demonstrated clinical activity in unselected patients (Orme and Huang, 2020). Our preclinical studies suggest that the more potent pan-HER inhibitor neratinib cooperates with enzalutamide in curbing the expansion of metastases generated by enzalutamide resistant and reprogrammed LNCaP cells. Since these cells contain a fraction of AR^+^ cells, which seemingly outgrow in response to adrenal androgens in castrated mice, the combination of neratinib with abiraterone or enzalutamide may be efficacious in cases of mixed ARPC/MSPC with elevated HER2/3 activity. Moreover, the combination of neratinib with the mTOR kinase inhibitor MLN0226 effectively inhibited both ERK and PI3K-mTOR signaling and suppressed the outgrowth of metastases generated by *PTEN* mutant DU145 cells. These findings suggest that rational neratinib combinations may be efficacious not only in ARPC but also in mixed ARPC/MSPC and MSPC.

In sum, the findings in this paper identify MSPC as a novel intrinsic subtype of M-CRPC and trace its origin to therapy-induced plasticity. Rather than favoring the outgrowth of tumor clones that have acquired certain oncogenic mutations, AR blockade is shown to directly induce reprogramming to a dedifferentiated stem-like state characterized by hybrid E/M traits and clinical aggressivity. Our results support a model in which transcriptional inactivation of TP53 induces hybrid E/M and stemness traits and inactivation of BMP signaling results in anti-androgen resistance, indicating that dedifferentiation and drug tolerance are mediated by separate mechanisms. Finally, this study identifies HER2/3 signaling as a major pathway prostate cancer cells rely on to emerge from drug-induced tolerance, an observation that deserves to be explored in biomarker-driven clinical trials.

## Acknowledgements

This work was supported by NIH grants R35 CA197566 (Outstanding Investigator Award to F.G.G.), U01 CA224044 (to Nora Navone, Y.C. and Andy Futtreal), and P30 CA016672 (MDACC Cancer Center Support Grant), by CPRIT Recruitment of Established Investigators Award RR160031 (to F.G.G.), by the generous philanthropic contributions to The University of Texas MD Anderson Moon Shots Program, and by the Korea Health Industry Development Institute (KHIDI) grant for research under the Biomedical Global Talent Nurturing Program HI19C0723 (to H.H.). We thank Peter Shepherd for assistance with the propagation of PDX models and members of the Giancotti laboratory for discussions. We thank the CPRIT Single Cell Genomics Core (RP180684) for support with single cell sequencing experiments.

## Author Contributions

F.G.G. conceived and led the study. F.G.G., H.H., and Y.W. conceived the hypotheses and designed and analyzed the experiments. H.H. and F.G.G. wrote the manuscript. H.H. and Y.W. performed and analyzed most of the experiments. H.H. generated and analyzed the prostate cancer xenograft, cell line and organoids transcriptome dataset and the human datasets. Y.W. performed and analyzed genome-wide RNA sequencing, ChIP sequencing and ATAC sequencing experiments. H.H., Y.W., S.G., X.W., and E.C. performed experiments with the LNCaP cells enzalutamide treatment. S.G., A.R., and X.W. performed experiments with the LNCaP TP53-KD and BRCA1-KD cells. H.H. and B.S. performed single cell RNA sequencing. J.C. performed the pharmacological inhibition experiments on the LNCaP and DU-145 cells. G.C., Y.W. and S.G. performed experiments on the role of BMP signaling. A.R., S.L., and Y.J. performed *in vitro* functional assays. Y.W., H. C., and M.H. performed bioinformatics analyses and Nick Navin provided guidance over the performance and analysis of single cell RNA sequencing data. D.K., B.Z. and S.G. participated in the characterization of the PDX model of reprogramming. S.N.M. performed histological analysis of the PDXs. A.R., S.L., and S.G. participated in the characterization of Pten^pc−/−^ mouse prostate cancer cells. X.W. helped with ChIP sequencing. Y.J. A.R. and B.Z. provided provision of reagents, materials, lab instruments and animals used in the manuscript. A.A. provided clinical expertise on AR-independent disease. Nora Navone, S.N.M. and A.A. provided PDXs and their transcriptome data. E.E. and C.J.L. provided access to the transcriptome data of prostate cancer tissues derived from the abiraterone-enzalutamide neoadjuvant clinical trial. Y.C. provided the prostate cancer patient-derived organoids and critical insights during the study.

## Methods

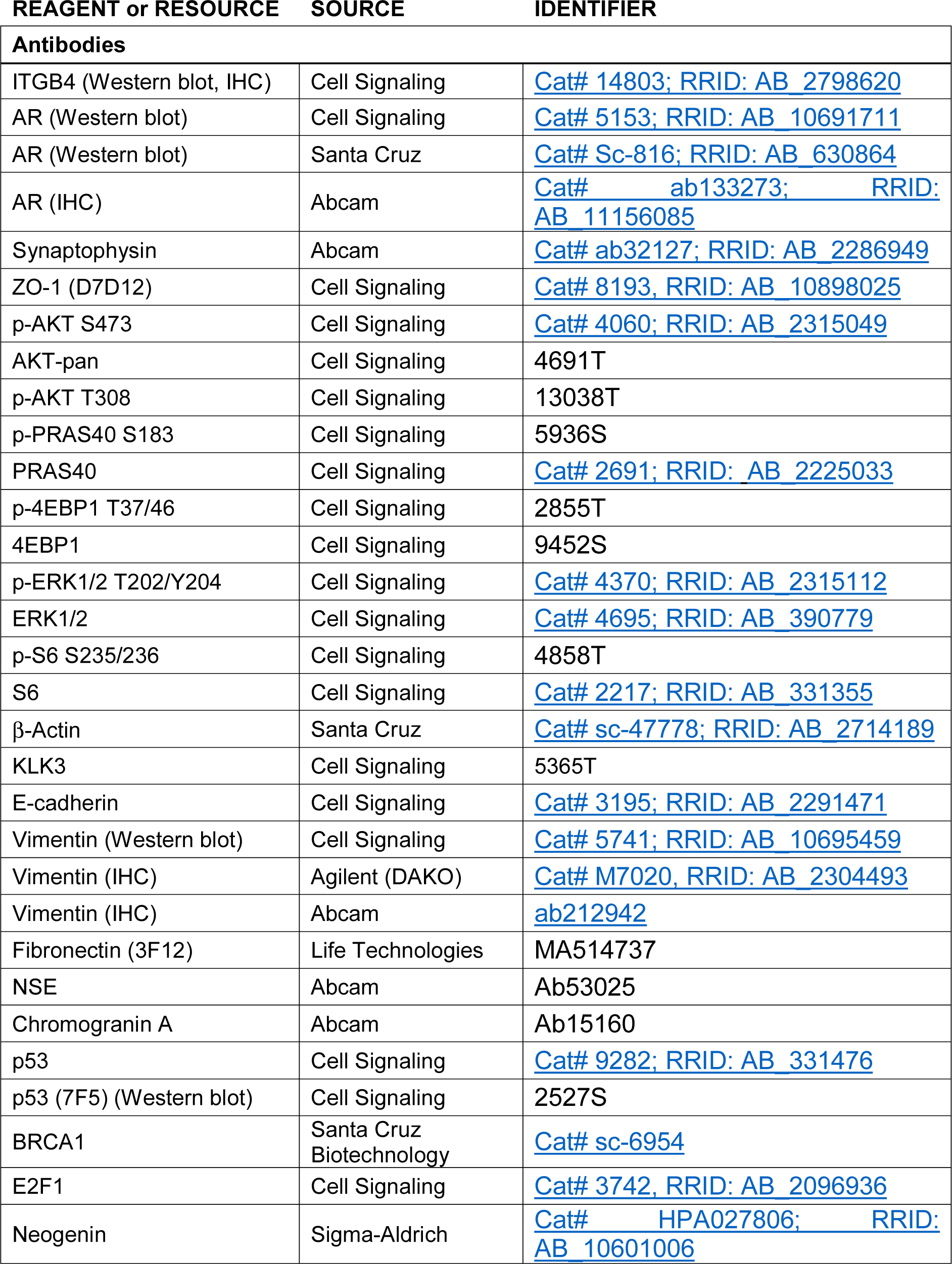
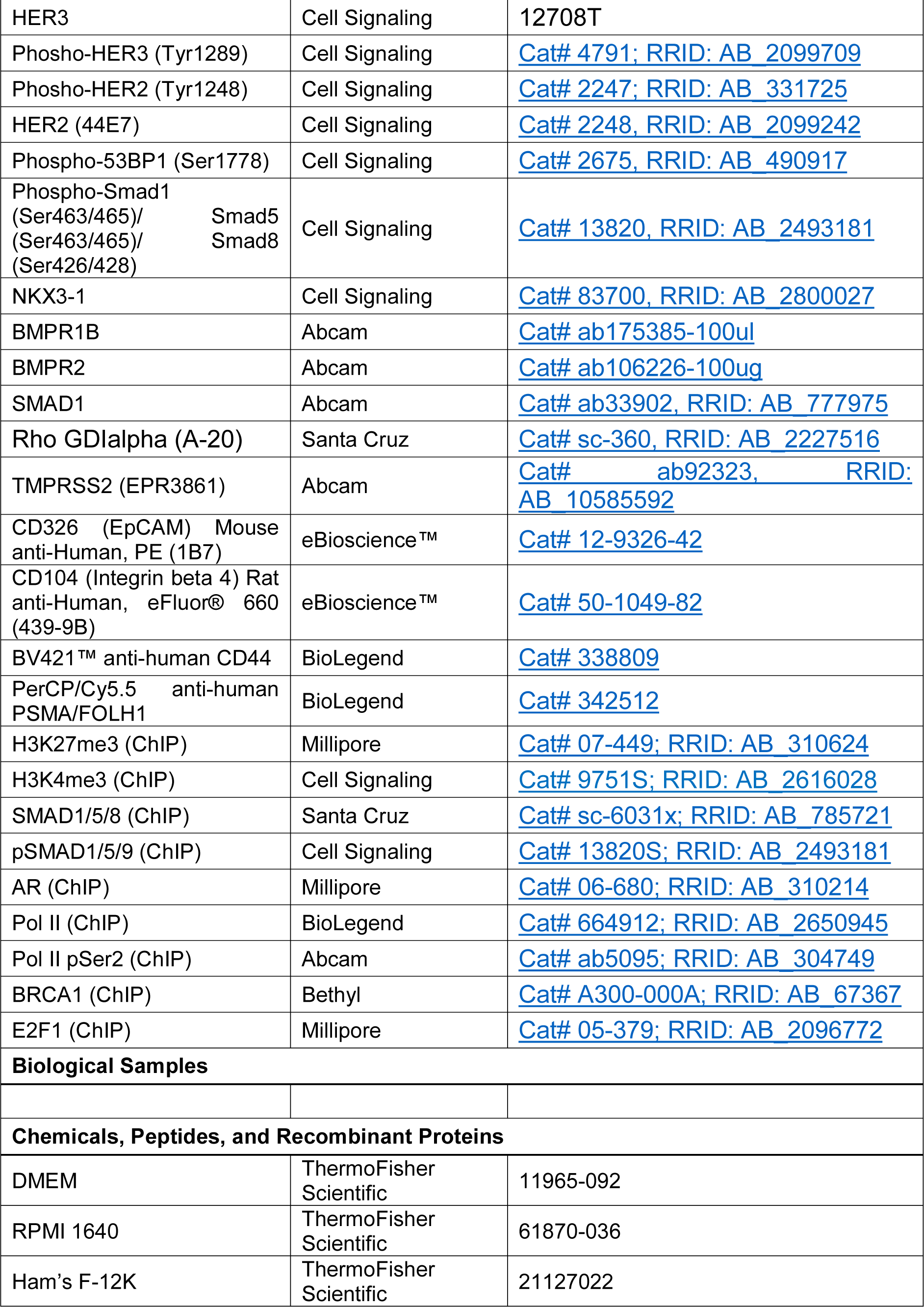
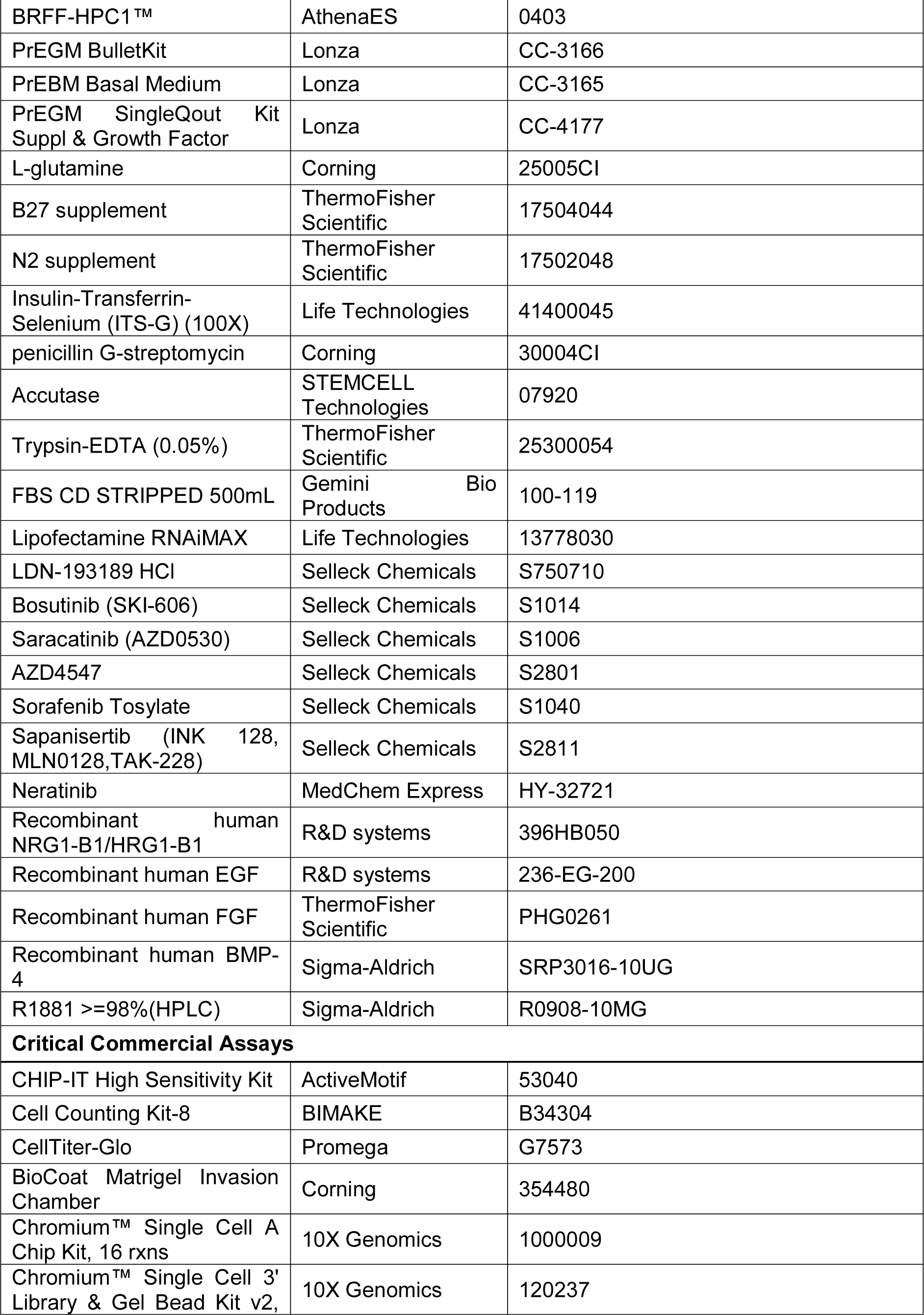
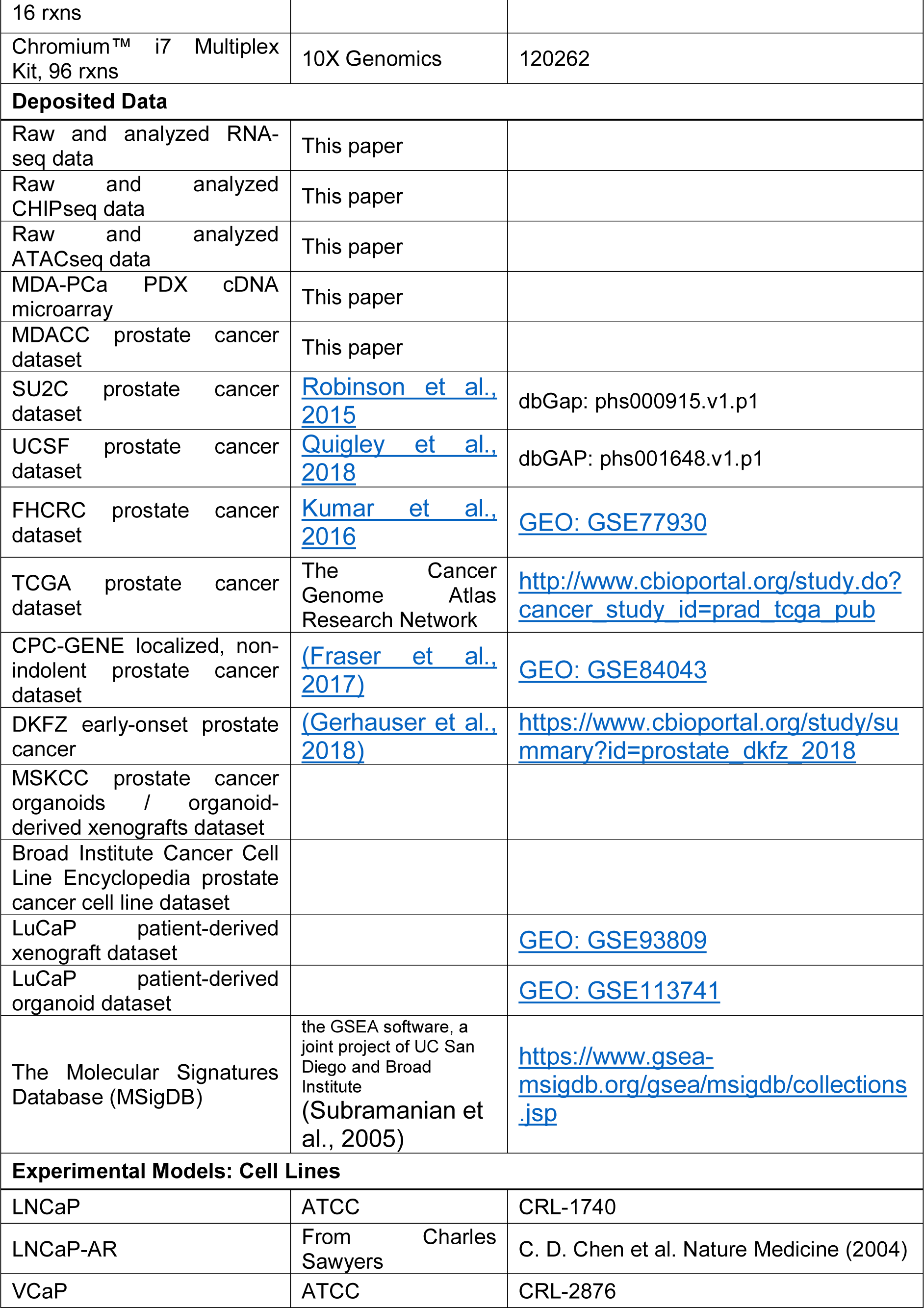
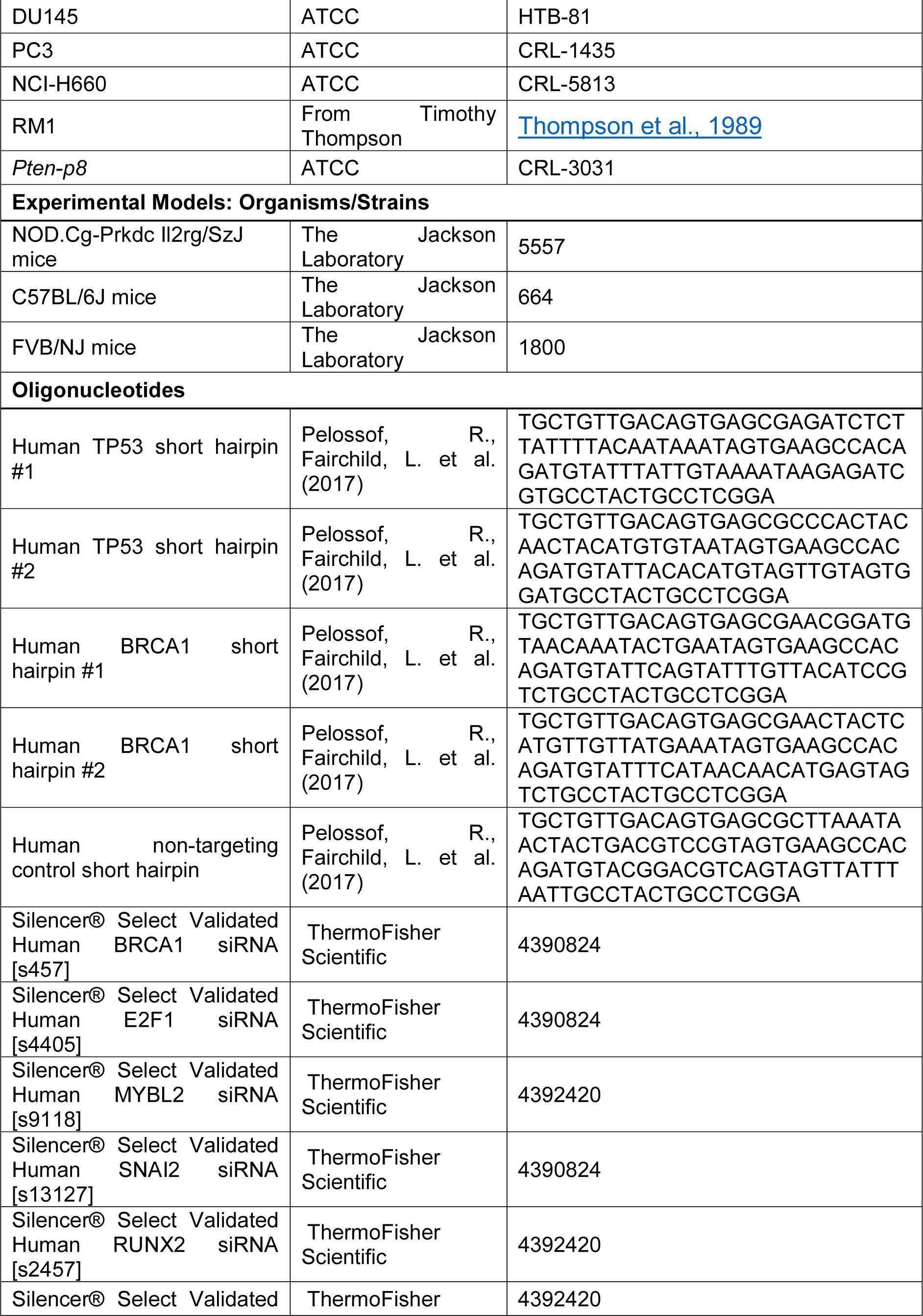
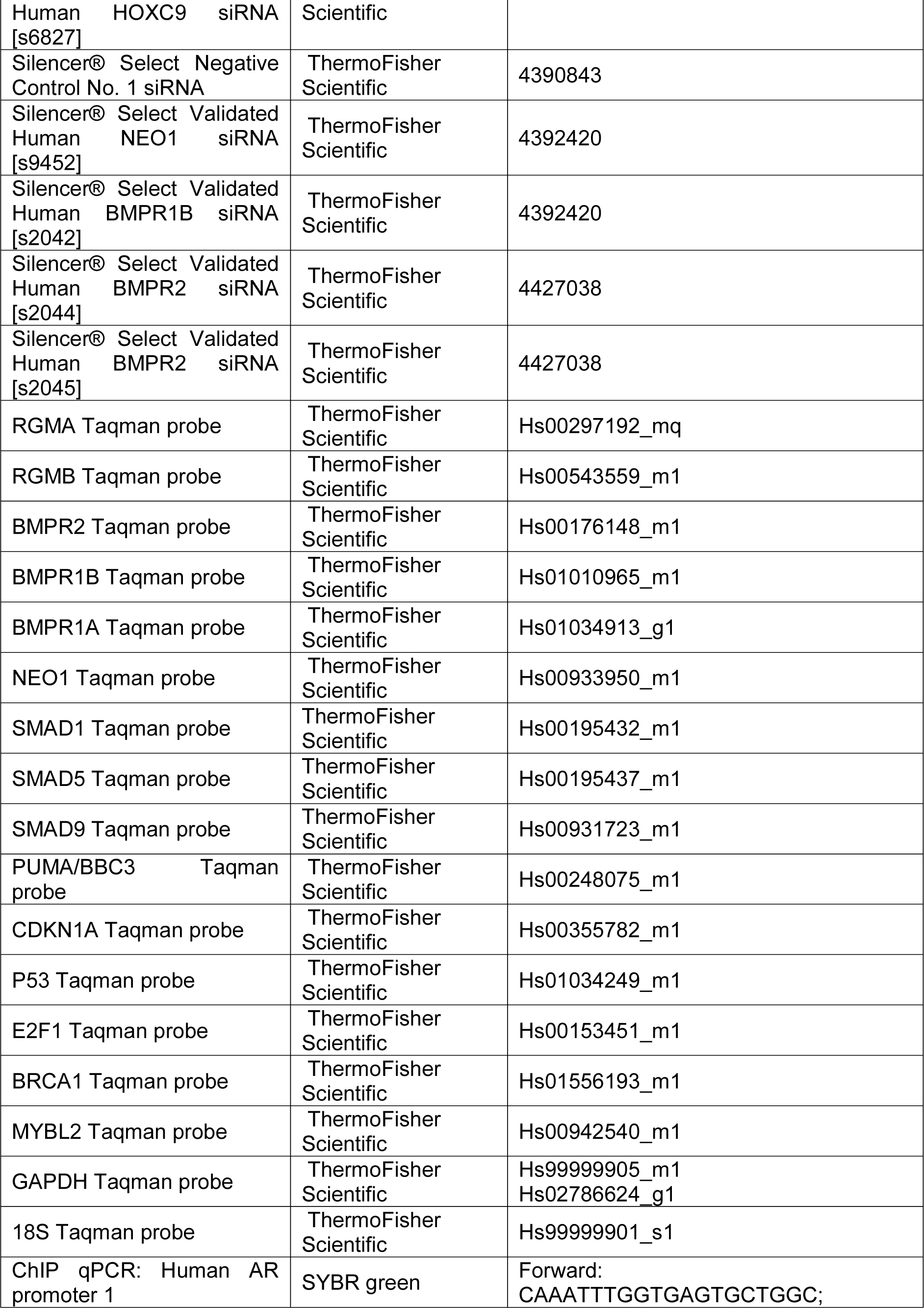
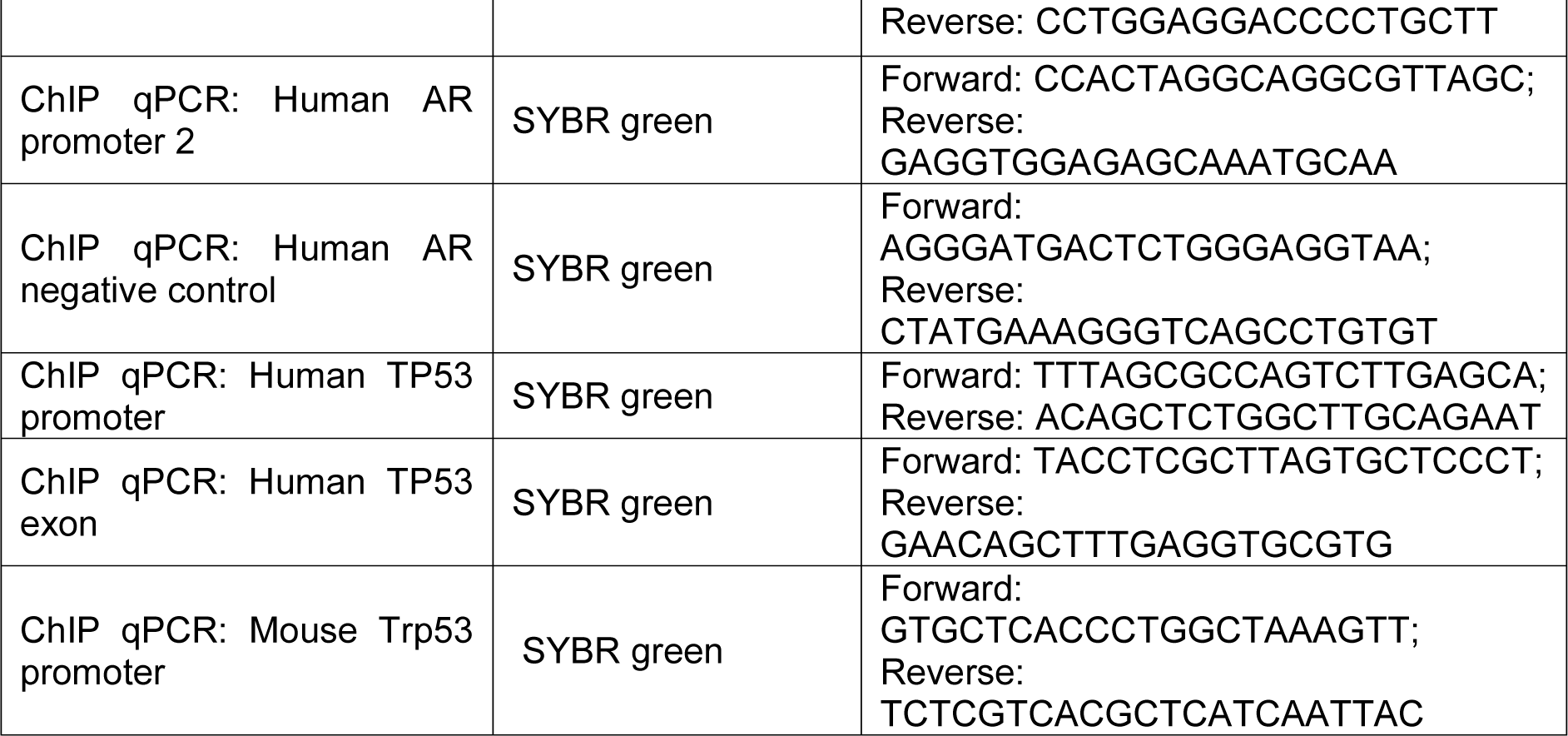

## EXPERIMENTAL MODEL AND SUBJECT DETAILS

### Cell Lines and Reagents

LNCaP, VCaP, DU145, PC3, NCI-H660 cells were obtained from ATCC, 293FT packaging cells were from Invitrogen and cultured according to manufacturers’ instructions. MDA-PCa-2B cells were obtained from Dr. Nora Navone’s laboratory and cultured in BRFF-HPC1 (AthenaES, MD) supplemented with 10% FBS and Gentamycin at 37°C in 5% CO2.

### Generation of *Pten*-p8^(-/-)^ Cell Line

The murine *Pten*-p8^(-/-)^ cell line was established by infecting the previously described *Pten*-p8^(-/+)^ cells (Jiao et al., 2007)with pLV-EGFP-Cre vector (Plasmid #86805). After transduction, EGFP-positive cells were selected by fluorescence activated cell sorting.

### Patient-derived Xenograft Models

Prostate cancer patient-derived xenograft (PDX) models were developed in the Prostate Cancer PDX program, Genitourinary medical oncology, MD Anderson Cancer Center accordingly (Li et al., 2008; Palanisamy et al., 2020). Partial characterization of some prostate cancer PDXs utilized in this work were published in: (MDA-PCa-118b, (Li et al., 2008)); (MDA-PCa-144-4, MDA-PCa-163-A, and MDA-PCa-177-B, (Aparicio et al., 2016)); (MDA-PCa-180-30, (Tzelepi et al., 2012)); (MDA-PCa-149-1, (Sircar et al., 2012)); (MDA-PCa-133, (Lee et al., 2011)). Fresh tumor chunks of PDX passage less than ten serial were provided from the MDA prostate cancer PDX program. Upon arrival, the specimens were placed in cold, sterile alpha-MEM (Gibco; Invitrogen), and small pieces were then implanted into subcutaneous pockets of 6- to 8-week-old male NOD SCID gamma mice (The Jackson Laboratory). The wound was closed either by Reflex 7mm Wound Clips (Roboz Surgical Instrument Co.) or 3M™ Vetbond™ Tissue Adhesive. Mice were monitored weekly for tumor growth. Once the initial implanted tumor grew in the mouse and reach certain size, tumor was collected and the extracts were prepared using T-PER tissue protein extraction regent (Thermo Fisher Scientific, Waltham, MA, USA) supplemented with protease and phosphatase inhibitors cocktails (Roche). Frozen tumor tissues were ground with mortar and pestle, incubated with the extraction buffer (2 mL of buffer per 0.1 g of tissue) in ice for 30 min, sonicated for 10 sec for 3 times in ice, centrifuged at 12,000 rpm for 5 min at 4°C, and the supernatant was collected and used for Western blot analysis. Tissue sections (4 μm) from formalin-fixed, paraffin-embedded (FFPE) PDX tumor tissue blocks were analyzed by immunohistochemical staining of AR (1:50, Dako), VIM (1:200, Hi pH, Dako) and ITGB4 (1:50, Cell Signaling) with use of an Autostainer Plus (Dako North America, Inc. Carpinteria, CA).

### Patient-derived Organoids

Prostate cancer patient-derived organoid culture was performed as described earlier (Gao et al., 2014).

### Animals

For all the animal studies in the present study, the study protocols were approved by the Institutional Animal Care and Use Committee (IACUC) of UT MD Anderson Cancer Center. Male BALB/c nude mice, male NOD SCID gamma mice, male C57BL/6J mice, and male FVB/NJ mice (aged 4-6 weeks) were obtained from The Jackson Laboratory. For localized tumor growth assay, cells were resuspended in 100 μL PBS with Matrigel in 1:1 ratio and subcutaneously injected into both rear flanks. The volume of the s.c. xenograft was calculated as V = L × W2/2, where L and W stand for tumor length and width, respectively. For experimental metastasis assays, cells were resuspended in 100 μL PBS and intracardially injected into the left ventricle with a 26G tuberculin syringe. For drug treatment, drug solution was delivered either intraperitonially or by oral gavage using 20G reuseable feeding needle (Roboz Surgical Co.). Metastatic burden was detected through noninvasive bioluminescence imaging of experimental animals using an IVIS Spectrum and Biospec 7T MRI instruments at the Small Animal Imaging Facility (SAIF). To investigate the effect of drug treatment, compounds were delivered daily through p.o. Bioluminescence signal was measured using the ROI tool in Living Image software (Xenogen).

## METHOD DETAILS

### Stable and Conditional Knockdown of Gene Expression

shRNAs were designed using the SplashRNA algorithm (Pelossof et al., 2017). Optimized lentiviral miR-E expression backbone system was used (Fellmann et al., 2013). Constitutive - SREP (Red, Puromycin); inducible - LT3RENIR (Red,Neomycin).

### Cell Proliferation Assay

Cells (5 × 10^3^ for LNCaP and PCa-2B, 2 × 10^3^ for all other cells) were seeded in a 96-well plate for 24 hours. After 24 hours, cells were cultured in DMEM or RPMI without phenol red containing 2% (vol/vol) FBS in the presence of given concentration of the compound(s). Viable cell numbers were measured by formazan formation using a Cell Counting Kit 8 (Dojindo). Apoptotic cells were detected by a standard TUNEL assay using an *in-situ* Cell Detection kit (Roche).

### Tumorsphere Assay

Single cells suspensions of tumor cells (1,000 cells/mL) were plated on ultra-low attachment plates and cultured in serum-free PrEGM (Lonza) supplemented with 1:50 B27, 20 ng/mL basic fibroblast growth factor (bFGF) and 40 ng/mL epidermal growth factor (EGF) for 10 days. Tumorspheres were visualized under phase contrast microscope, photographed, and counted. For serial passage, tumorspheres were collected using 70-μm cell strainers and dissociated with Accutase (Stem Cell Technologies) for 30 min at 37°C to obtain single cell suspensions.

### Cell Invasion Assay

Cell invasion was assayed using Matrigel coated BioCoat Cell Culture Inserts (24-well plates, Corning). After Matrigel was rehydrated at room temperature, 2 x 10^5^ cells suspended in 0.5 mL RPMI medium were plated into each insert. 0.5 mL medium with 15% FBS or CSS were added into the bottom of each well. Noninvading cells were removed after 48 hours culture. The cells on the lower surface of the membrane were stained with crystal violet.

### Matrigel 3D Culture

Dissociated cells were incubated in PrEGM medium (Lonza) supplemented with 1:50 B27, 20 ng/mL basic fibroblast growth factor (bFGF) and 40 ng/mL epidermal growth factor (EGF, Corning). Matrigel bed was made in 6 well plate by putting 4 separate drops of Matrigel per well (50 μL Matrigel per drop). Plates were placed in 37°C CO_2_ incubator for 30 min to allow the Matrigel to solidify. For each sample, 100 μL of cell suspension was mixed with 100 μL cold Matrigel, and pipetted on top of the Matrigel min. Warm L each). The plates were then incubated at 37°C for another 30L PrEGM (2.5 mL) was then added to each well. The cells were cultured and monitored for 10-14 days with 50% medium change every 3 days. For immunostaining experiments, the cells were cultured in 8 well chamber slide. Cells were fixed with 4% paraformaldehyde for 20 minutes and standard immunostaining protocol was then followed.

### Prostate Organoid Culture

Mouse and human prostate cancer cell organoid forming assay (embedding method) was performed as described earlier (Chua et al., 2014). Prostate cancer cells were resuspended in prostate organoid culture medium, consisting of: hepatocyte medium supplemented with 10 ng mL^−1^ EGF, 10 μM Y-27632 (STEMCELL Technologies), 1 × Glutamax (Gibco), 5% Matrigel (Corning) and 5% charcoal-stripped FBS, which had been heat-inactivated at 55 °C for 1 h. After resuspension in prostate organoid medium, the resulting cell suspension containing 500–3,000 dissociated cells was mixed with 60 μL of Matrigel, and the mixture was pipetted around the rim of wells in a 24-well plate. The mixture was allowed to solidify for 30 min at 37 °C, before addition of 400 μL organoid culture medium to each well. The culture medium was changed every other day, and organoids were counted after 8-10 days. The efficiency of organoid formation was calculated by averaging the number of organoids visible using a 10X objective. For statistical analyses, efficiency percentages were arcsine converted to perform unpaired two-tailed Student’s t-test.

### Bioluminescence and X-ray Imaging

For bioluminescent imaging, mice were anesthetized and injected with 1.5 mg of D-luciferin intraperitoneally at the indicated times. Animals were imaged in an IVIS 100 chamber within 5 min after D-luciferin injection, and data were recorded using Living Image software (Xenogen). Photon flux was calculated by using the ROI tool in Living Image software. Bone metastases were further confirmed by X-Ray imaging using IVIS Lumina XR equipped with X-ray and Optical Overlay.

### Immunohistochemistry Staining

Immunohistochemistry on paraffin-embedded sections and immunofluorescent staining were performed using a Discovery XT processor (Ventana Medical Systems). The tissue sections were deparaffinized with EZPrep buffer (Ventana Medical Systems), antigen retrieval was performed with CC1 buffer (Ventana Medical Systems). Sections were blocked for 30 minutes with Background Buster solution (Innovex), followed by avidin-biotin blocking for 8 minutes (Ventana Medical Systems). Sections were incubated with anti-AR (Abcam, ab133273 1 μg/mL); ITGB4 (Cell Signaling, cat# 14803, μg /mL); Vimentin (Cell Signaling, cat# 5741, 0.5 μg /mL); anti-Synaptophysin μg/mL) for 5 hours, followed by 60 minutes incubation with (Abcam, ab32127, 1 μ biotinylated horse anti-rabbit (Vector Labs, cat# PK6101) at 1:200 dilution (for AR, ITGB4, Vimentin), HRP-conjugated goat anti-rabbit (PI-1000) at 1:250 dilution (for synaptophysin). The detection was performed with DAB detection kit (Ventana Medical Systems) according to manufacturer instruction. Slides were counterstained with hematoxylin and cover-slipped with Permount (Fisher Scientific).

### Western Blot

For immunoblotting, cells were washed twice with PBS and lysed in RIPA buffer (50 mM Tris-HCl pH 7.4, 150 mM NaCl, 1 mM EDTA, 1% Triton X-100, 1% sodium deoxycholate, and 0.1% SDS) supplemented with protease inhibitors (Protease/Phosphatase Inhibitor Cocktail (100X), Cell Signaling #5872). Total protein concentrations were determined by using Pierce™ BCA Protein Assay Kit (ThermoScientific™, 23225). Cell extracts concentrations were brought down to 1ug/uL with Sample Buffer 4X and boiled for 5min before gel loading.

### NRG1 FISH labeling experiment

For NRG1 RNA FISH experiment, the procedure was carried out according to the manufacturer’s instructions (Stellaris® RNA FISH Protocol for Frozen Tissue; Biosearch Technologies, Inc., CA). Mouse NRG1 probe (5 nM total, labeled with CAL Fluor® Red 610 Dye) was used in this study: Sequences of custom probe sets are listed in Supplementary. All hybridizations were done overnight in the dark at 37°C in a humidifying chamber.

### Immunofluorescence Staining

Cell were plated on Falcon™ Chambered Cell Culture Slides (Corning Inc) and cultured (specific condition and duration indicated in each experiment figure). Cells were fixed, washed and stained for antibodies (primary and secondary) and visualized.

### Prostate Cancer Patient Datasets

Prostate cancer patient sample gene expression and amplification data were acquired from the cBioportal database. Additionally, the UCSF metastatic prostate cancer patient dataset was kindly provided by the authors (Quigley et al., 2018). Z-score 2.0 was used as cut-off value to determine mRNA up/downregulation in a given sample. For the UCSF dataset, copy number alteration was called using following log2 ratio bounds, as used in the original paper: - chr1-chr22 Gain / shallow loss / deep loss: 3 / 1.65 / 0.6-chrX, chrY Gain / loss: 1.4, 0.6. Morpheus was used for clustering and heatmap generation (https://software.broadinstitute.org/morpheus).

### Prostate Cancer Patient-derived Xenografts, Cell Lines and Organoids (PCO-94) Dataset Generation and Processing

Five gene expression datasets of castration-resistant prostate cancer patient-derived xenografts, cell lines and organoids were merged into a single data table by HUGO gene symbol as reference. MDA PCa PDX: microarray data of MDA PCa PDX, including tumors of same origin but grown in castrated and uncastrated hosts (133-4_cas1,2; 180-30_cas1,2) (Tzelepi et al., 2012); MSKCC PCa organoid/ODX (organoid-derived xenograft): mRNA expression (RNA Seq FPKM) data available for 10 of 12 PCa organoids (Gao et al., 2014). CCLE PCa Cell lines gene expression (RNA Seq RPKM) data (Release date: 14-Feb-2018. Broad Institute). LuCaP M-CRPC PDXs custom Agilent 44k whole genome expression microarray (includes early/late and castration-resistant passages. GSE93809. Nguyen et al., 2017). LuCaP PDX-derived organoids (RNA Seq TPM) data (includes two repeats. GSE113741. Beshiri et al., 2018). Microarray data were transformed to non-logarithmic scale. ASAP v1 “Automated Single Cell Analysis Pipeline” (http://asap-old.epfl.ch) was utilized for following data processing.

1. Log2 conversion and batch correction using “ComBat” method (Johnson et al., 2007)
2. Plotting by classical MultiDimensional Scaling (MDS) or t-SNE.
3. Cluster Identification (see details in “PCO-94 Dataset Clustering”)
4. Retrieve Differentially Expressed Genes (DEGs) among clusters by using Limma method (Ritchie et al., 2015)
5. Geneset enrichment analysis of the DEGs

### PCO-94 Dataset Clustering and Determination of Optimal K

As preliminary, we tried three independent clustering methods (PAM “Partitioning Around Medoids”, K-Means and Hierarchical clustering), three data sources (normalized data, MDS values and t-SNE values after dimension-reduction) and n of cluster as either two or three (suggested by silhouette analysis), total 18 combinations (3x3x2). The largest cluster containing 50 samples (later determined as ARPC) showed 17 of 18 concordance rate across clustering combinations. The smallest cluster containing 13 samples (later determined as NEPC) showed 18 of 18 concordance across clustering combinations. Among the remaining 31 samples, 21 samples showed more than 6 of 9 concordance rate (later determined as MSPC) when n of cluster was three, and the remaining 10 samples showed inconsistent results across the combinations (later determined as mixed). Overall, about 80% of the samples were consistently clustered together in the test. In this manuscript, we used PAM as final representative clustering method, and the MDS values as data source for clustering. To confirm the optimal n of cluster (K), we performed Consensus clustering (clustering.algorithm = SOM; cluster.by = columns; distance.measure = Euclidean; resample = subsample; merge.type = average; descent.iterations = 2,000; normalize.type = row-wise; normalization.iterations = 0), and calculated PAC (proportion of ambiguous clustering). Final optimal K was determined as three by the lowest PAC.

### CIBERSORT Tumor Deconvolution and Estimation of Subtype Abundances

To compute intratumoral heterogeneity, a deconvolution method “CIBERSORT” (Newman et al., 2015) was applied. We followed “Custom Signature Genes File” tutorial-mixture file: full PCO-94 dataset after batch correction (gene n: 14,968); reference sample file: reduced PCO-94 dataset (gene n: 2,424); phenotype class file: annotation of clusters determined by PAM clustering of full PCO-94 dataset. Specifically, the gene signature was defined by the average expression values of 2.4K differently expressed genes from the PCO-94 clustering results. RNA-seq read-normalized gene expression values (RSEM, RPKM, and FPKM for TCGA, CPC-GENE and DKFZ, and SU2C-PCF and UCSF datasets, respectively) or microarray data (FHCRC dataset) with Entrez gene ID and HUGO gene symbol annotations were loaded as a “mixture” file. Purity was defined by the estimated abundances of each cluster types in a sample. Samples were designated as “mixed” if the largest component < 75% (cell lines, organoids, PDXs) or < 60% (human tissues).

### Bulk RNA-seq Analysis

Total RNA were extracted from samples using TRIzol (Invitrogen, USA), then 1 µg was then sent to BGI for quality testing and library construction. Libraries were sequenced on a BGISEQ-500. RNA-seq reads were aligned to the human reference genome (hg19) using Tophat (v2.1.1; https://ccb.jhu.edu/software/tophat/index.shtml)(Kim et al., 2013). Gene models of Refgene were downloaded from the Illumina’s iGenomes project (https://support.illumina.com/sequencing/sequencing_software/igenome.html). FPKM (Fragments Per Kilobase of transcript per Million mapped reads) values were generated using cufflinks (v2.2.1; http://cole-trapnell-lab.github.io/cufflinks/)(Trapnell et al., 2013). Further differential expression analysis was using cuffdiff function which is in cufflinks, and considered genes with log2 fold change > 4 or < -2 and false discovery rate (FDR) < 0.05 as significantly differentially expressed.

### Single Cell RNA-seq Analysis

For single cell RNA sequencing, the Chromium Single Cell 3 Library and Gel Bead Kit v2 (10X Genomics Inc) were applied according to the manuf^′^acturer’s protocol. Briefly, single cell suspensions of LNCaP cells were counted and loaded on individual lanes of a Single Cell A Chip with appropriate reagents. The chip was ran in the Chromium™ Single Cell Controller to generate single cell gel bead-in-emulsions (GEMs) for sample and cell barcoding. Libraries were generated using 10x Genomics’ protocol, pooled and sequenced by Illumina in Hiseq 4000 sequencer. Five samples (Day 0, Day 1, Day 3, Day 5, Day 7) were aligned with human genome GRCh38 using STAR version 2.5.1b (Dobin et al., 2013) and further aggregated using Cell Ranger v2.0.2 for analysis, resulting 6,072 cells of 205,881 mean reads per cell (post-normalization) and 6,164 median genes per cell.

For clustering of the aggregated data, we used the graph-based clustering algorithm implemented in the Cell Ranger pipeline, which consists of building a sparse nearest-neighbor graph (where cells are linked if they among the k nearest Euclidean neighbors of one another), followed by Louvain Modularity Optimization (LMO) (Blondel et al., 2008). The value of k is set to scale logarithmically with the number of cells. Additional cluster-merging approaches included performing hierarchical clustering on the cluster-medoids in PCA space and merging pairs of sibling clusters if there are no genes differentially expressed between them (with B-H adjusted p-value below 0.05). The hierarchical clustering and merging is repeated until there are no more cluster pairs to merge. From the resulting 8 clusters, two of them showed significant low UMIs than the rest and enriched by mitochontrial genes or ribosomal genes. They were excluded from further analysis in this manuscript.

### Single Cell Trajectory Inference

The reconstruction of single cell trajectory was done with Monocle 3 (Cao et al., 2019; Qiu et al., 2017; Trapnell et al., 2014). First the 10X Genomics Cell Ranger output was loaded into Monocle 3 using load_cellranger_data function, and the data was pre- processed using PCA method. Here we chose 100 principal components (PCs; num_dim = 100) to make sure most of the variation in gene expression across all the cells was captured. The dimensionality of the data was reduced with UMAP algorithm (McInnes et al., 2018) and mutually similar cells were grouped into clusters using a technique called Louvain community detection. Each cell is assigned not only to a cluster but also to a partition. Next, we fitted a principal graph within each partition using the learn_graph() function. And cells were ordered according to their progress along the learned trajectory. In our time series experiments, we chose cells in the UMAP space from early time point (here Control0) as "roots" of the trajectory. We mainly focused on the trajectory within the large partition.

### Chromatin immunoprecipitation (ChIP)

1x10^6^ Cells were crosslinked by using 2mM DSG for 45 min, then by using 1% formaldehyde for 15 min, which were both performed at room temperature (RT). To stop crosslink, glycine was added to final concentration of 0.125M, then incubate at RT for 5 min. Cells were collected by scraping from dishes, then washed with PBS 3 times. Resuspend pellets in 0.5ml of SDS lysis buffer (1% SDS, 10mM EDTA, 50mM Tris-HCl, pH8.0)/PIC/PMSF/Sodium butyrate mix, then incubate on ice for 10 min. Sonicate the crosslinked cellular lysate with Diagnode sonicator. After sonication, aliquot samples into a 1.7ml tube. Centrifuge at max speed for 10 min at 4. Transfer supernatant to a new 1.7ml tube. To prepare chromatin immunoprecipitation sample, per 0.1ml of sonicated sample, add 0.9ml of dilution buffer (50mM Tris-HCl, pH8.0, 0.167M NaCl, 1.1% Triton X-100, 0.11% sodium deoxycholate)/PIC/PMSF/Sodium butyrate mix, and then add antibody bound Dynabeads. Gently mix and place on rocker O/N at 4 Place tube in magnetic stand. Invert several times. Allow beads to clump. Discard supernatant. Perform the following wash steps with 0.8ml of cold buffer. Flick tubes to resuspend beads and incubate each wash for 5min on rocker at 4. Place tube in magnetic stand. Invert several times. Allow beads to clump and discard supernatant. 1 time with RIPA-150, 1 time with RIPA-500, 1 time with RIPA-LiCl, 2 times with 1xTE Buffer, pH8.0. Resuspend beads in 200ul freshly made Direct Elution Buffer (10mM Tris-HCl pH8.0, 0.3M NaCl, 5mM EDTA, 0.5% SDS). Add 1ul of RNase A and incubate O/N at 65 to reverse crosslink. Quick spin sample. Place in magnetic stand. Allow beads to clump and transfer supernatant to a new low-bind tube. Add 3ul of Proteinase K and incubate for 1-2hrs at 55. Purify the reverse-crosslinked ChIP DNA sample using phase lock tubes and EtOH precipitation. Resuspend sample in 25ul of Qiagen elution buffer. DNA was amplified by real-time PCR (ABI Power SYBR Green PCR mix).

### Immune Cell Subset Deconvolution Analysis

Intratumoral immune cell subsets from the SU2C, FHCRC and UCSF M-CRPC datasets were analyzed by using CIBERSORT bulk transcriptome deconvolution technique (Newman et al., 2015). We used the LM22 signature genes file consisting of 547 genes that accurately distinguish 22 mature human hematopoietic populations and activation states, including seven T cell types, naïve and memory B cells, plasma cells, NK cells, and myeloid subsets.

### Gene Set Enrichment Analysis

We used v3.0 of java GSEA program (Subramanian et al., 2005).

### Single Sample GSEA Projections and Visualizations

We carried out ssGSEA (Barbie et al., 2009) using the GenePattern module ssGSEA Projection (v9) (www.genepattern.org). We used Prism (v8) for data visualization and related statistical analysis. Genesets used for the analysis are from the Molecular Signature Database, including their hallmark genesets (Liberzon et al., 2015).

### Sample and library preparation for ChIP-seq and ATAC-seq

ChIP-seq sample preparations were performed as ChIP experiments. Libraries were prepared according to standard illumina protocol. Libraries were sequenced at Sequencing and Microarray Facility at MDACC. For ATAC-seq, 5x10^5^ LNCaP cells were prepared and collected. Cells were then washed once with cold 1xPBS and spinned down at 500g for 5min at 4. Cells were kept on ice and subsequently resuspended in 25ul 2xTD buffer (Illumina Nextera kit), 2.5ul Transposase enzyme (Illumina Nextera kit, 15028252) and 22.5ul Nuclease-free water in total of 50ul reaction for 1hr at 37. DNA was then purified using Qiangen MinElute PCR purification kit (28004) in a final volume of 10ul. ATAC-seq Libraries were prepared following the Buenrostro protocol (http://www.ncbi.nlm.gov/pmc/articles/PMC4374986/) and ATAC-seq libraries were sequenced as 50 base paired-end reads on the DNBseq platform at the BGI Americas.

### Analysis of ATAC-seq Data

We utilized cutadapt (v1.18) (Martin, 2011) for the raw reads to remove the adaptor sequence or the reads shorter than 35bp and then aligned those trimmed reads to the human reference genome (hg19) using default parameters in Bowtie2 (v2.4.1) (Langmead and Salzberg, 2012). The aligned reads were subsequently filtered for quality and uniquely mappable reads were retained for further analysis using Samtools (v1.10) (Li et al., 2009). Relaxed peaks were called using MACS2 (v2.1.2) (Feng et al., 2012) with a p value of 1X10-2. Consensus peaks were calculated by taking the overlap of peaks for samples. Genome-wide read coverage was calculated by BEDTools (Quinlan and Hall, 2010). In order to calculate the ATAC-seq read density at the promoters and the enhancers, normalized read densities (RPKM, Reads Per Kilobase per Million mapped reads) were calculated across the gene promoter regions and the enhancer regions, coordinately. The promoters used in this study were defined as 1 kb upstream and 1kb downstream of the transcription start site determined based on the UCSC gene annotation. The annotation of the enhancers was from the FANTOM5 and the GenHancer. The annotation of the indicated transcription factors binding motif was from the HOMER (Heinz et al., 2010). Identification of significantly over-represented functional categories was done using function of “Investigate Gene Sets” from GSEA (http://software.broadinstitute.org/gsea/msigdb/annotate.jsp)(Mootha et al., 2003).

### Analysis of ChIP-seq Data

Sequencing reads from H3K4me3 and H3K27me3 ChIP-seq were trimmed by using trimmomatic (v0.39)(Bolger et al., 2014). The trimmed reads were then aligned to the human genome (hg19) using the Bowtie2 software. PRC duplication reads were removed by Samtools. The following up peaks calling and reads density were calculated by using the same methods as we did for ATAC-seq. The promoters used in this study were defined as 5 kb upstream and 5kb downstream of the transcription start site determined based on the UCSC gene annotation. To visualize ChIP-seq signal at individual genomic regions, we used the UCSC Genome Browser (https://genome.ucsc.edu)(Kent et al., 2002).

### FACS Analysis

Cells were detached with Accutase and washed in blocking solution (HBSS supplemented with 10% FBS). Cell suspensions were incubated with the indicated antibodies for 45 minutes at 4°C and analyzed by FACS. At the end point of the *in vivo* experiment, blood and bone marrow cells were collected from each mouse and treated with Red Blood Cell lysis buffer for 5 minutes. Cells were then washed once with RPMI supplemented 2% FBS, stained with indicated antibodies for 45 minutes and analyzed by FACS.

### Analysis of Protein and mRNA Expression

For immunoblotting, cells were washed with PBS and lysed in RIPA buffer (50 mM Tris-HCl pH 7.4, 150 mM NaCl, 1 mM EDTA, 1% Triton X-100, 1% sodium deoxycholate, and 0.1% SDS) supplemented with protease inhibitors (Calbiochem) and phosphatase inhibitors (PhosSTOP, Roche Life Science). Protein concentrations were measured by using the DC Protein Assay. Total RNA was extracted using the RNeasy Mini kit coupled with RNase-free DNase set (Qiagen) and reverse transcribed with SuperScript™ IV VILO™ Master Mix with ezDNase™ Enzyme (Invitrogen). cDNA corresponding to approximately 10 ng of starting RNA was used for one reaction. qPCR was performed with Taqman Gene Expression Assay (Applied Biosystems). All quantifications were normalized to endogenous GAPDH.

## QUANTIFICATION AND STATISTICAL ANALYSIS

Statistical analyses used R and GraphPad Prism 8 software, with a minimum of three biologically independent samples for significance. For animal experiments with subcutaneous injections, each subcutaneous tumor was an independent sample. For intracardiac injection and survival analysis, each mouse was counted as a biologically independent sample. Results are reported as mean ± SD or mean ± SEM. Comparisons between two groups were performed using an unpaired two-sided Student’s t-test (p < 0.05 was considered significant). Comparison of multiple conditions was done with one- or two-way ANOVA test. For correlation analysis, the Spearman coefficient and Pearson coefficient were used. All experiments were reproduced at least three times, unless otherwise indicated.

---

**Suppl. Fig. 1.**
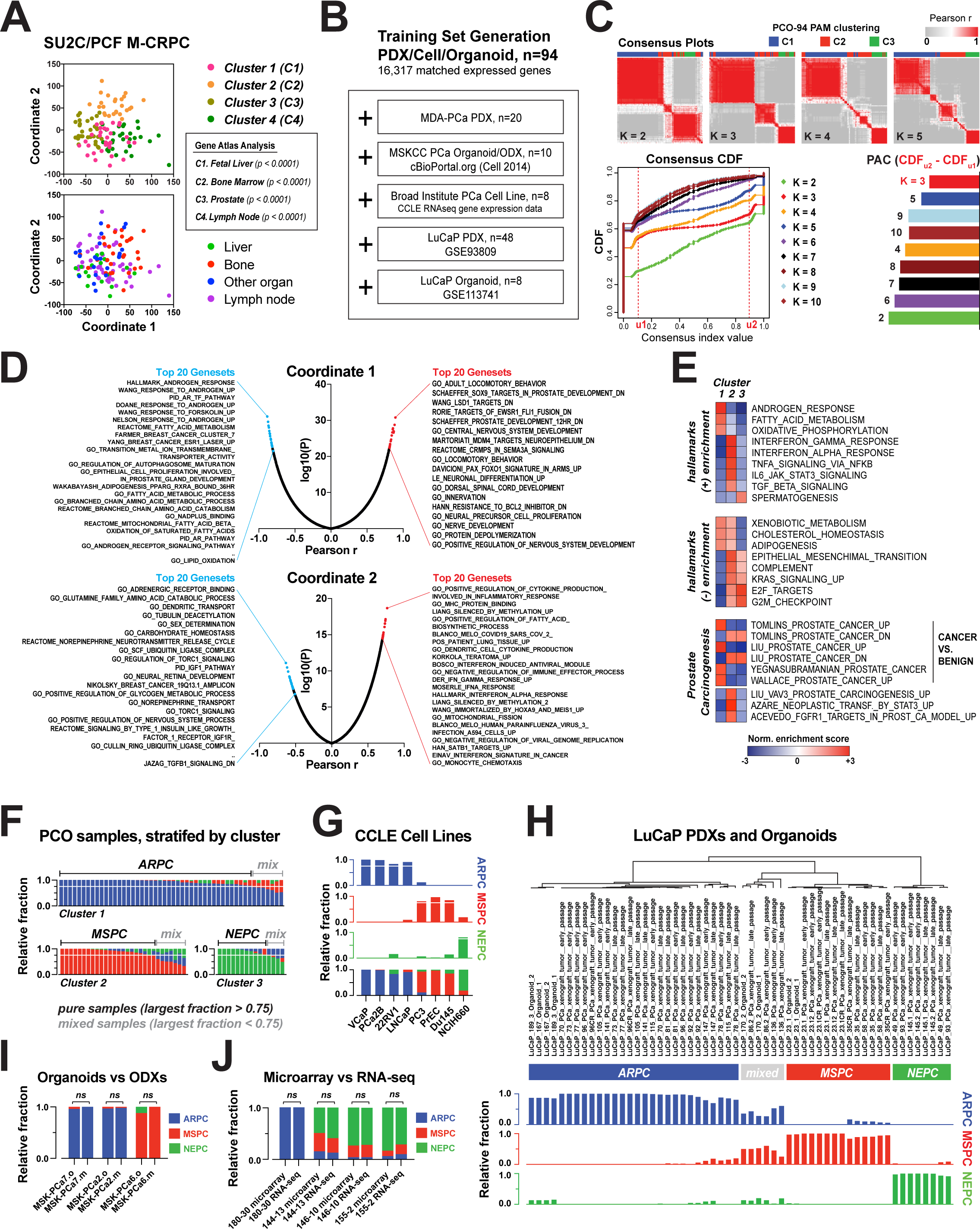

**Suppl. Fig. 2.**
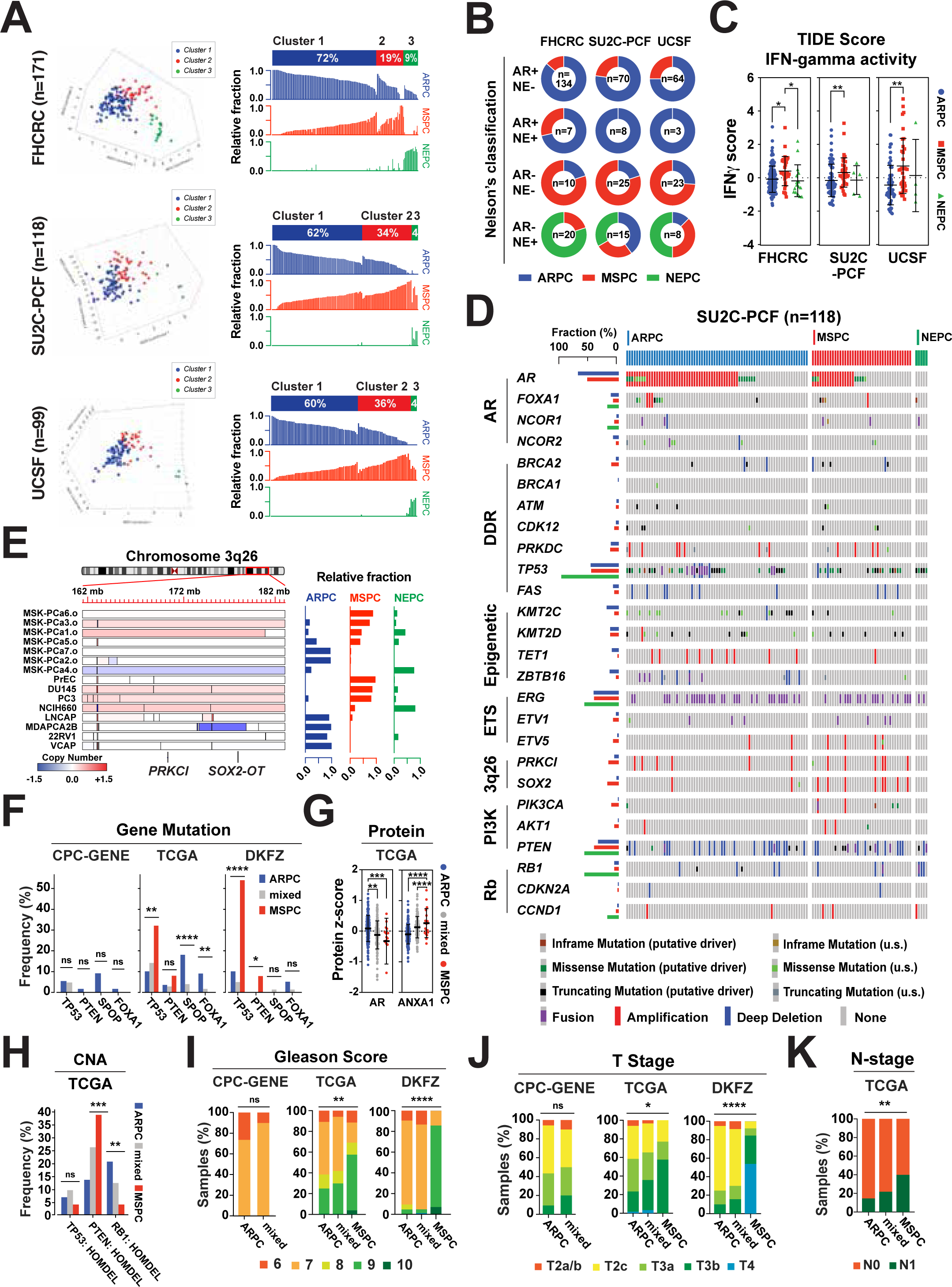

**Suppl. Fig. 3.**
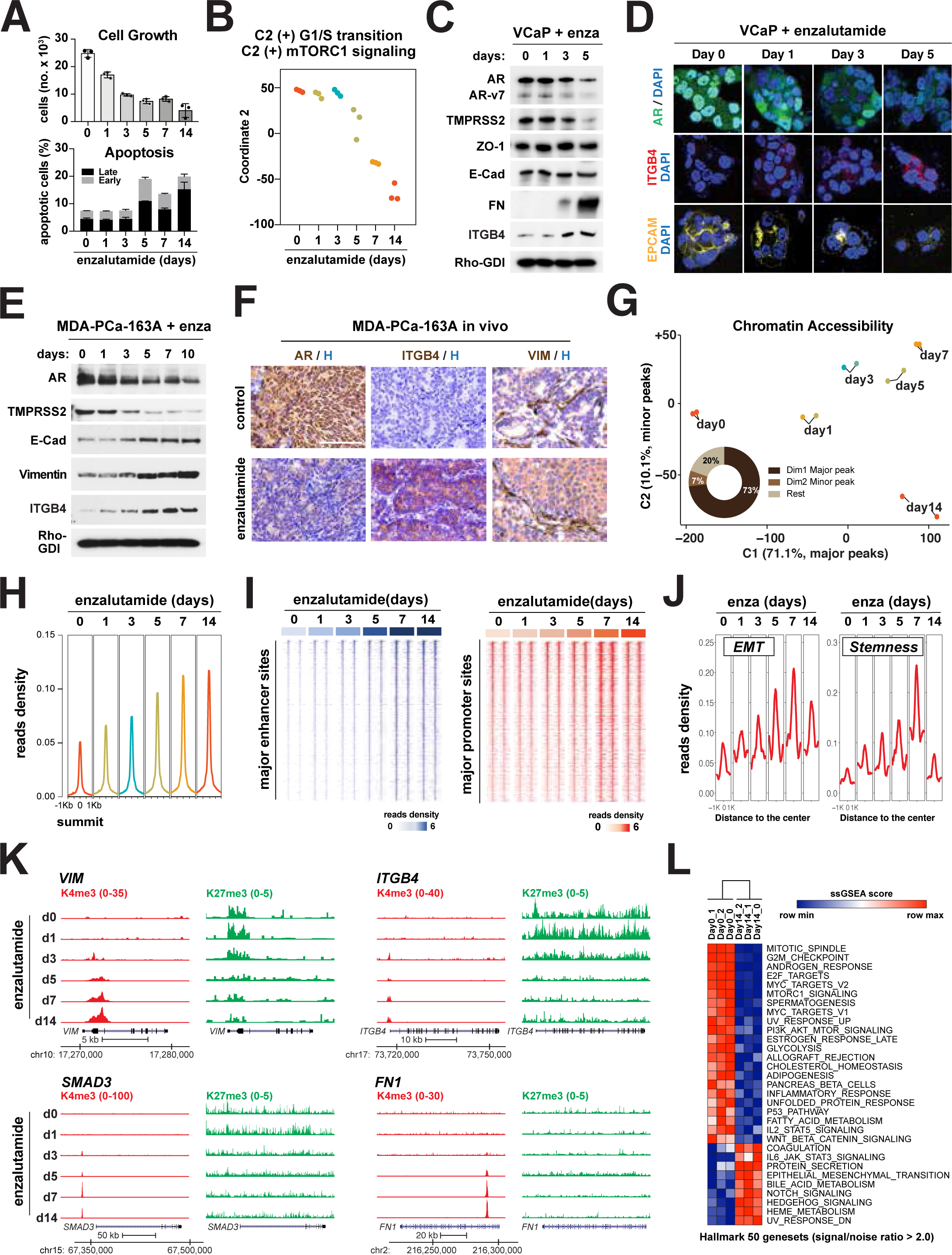

**Suppl. Fig. 4.**
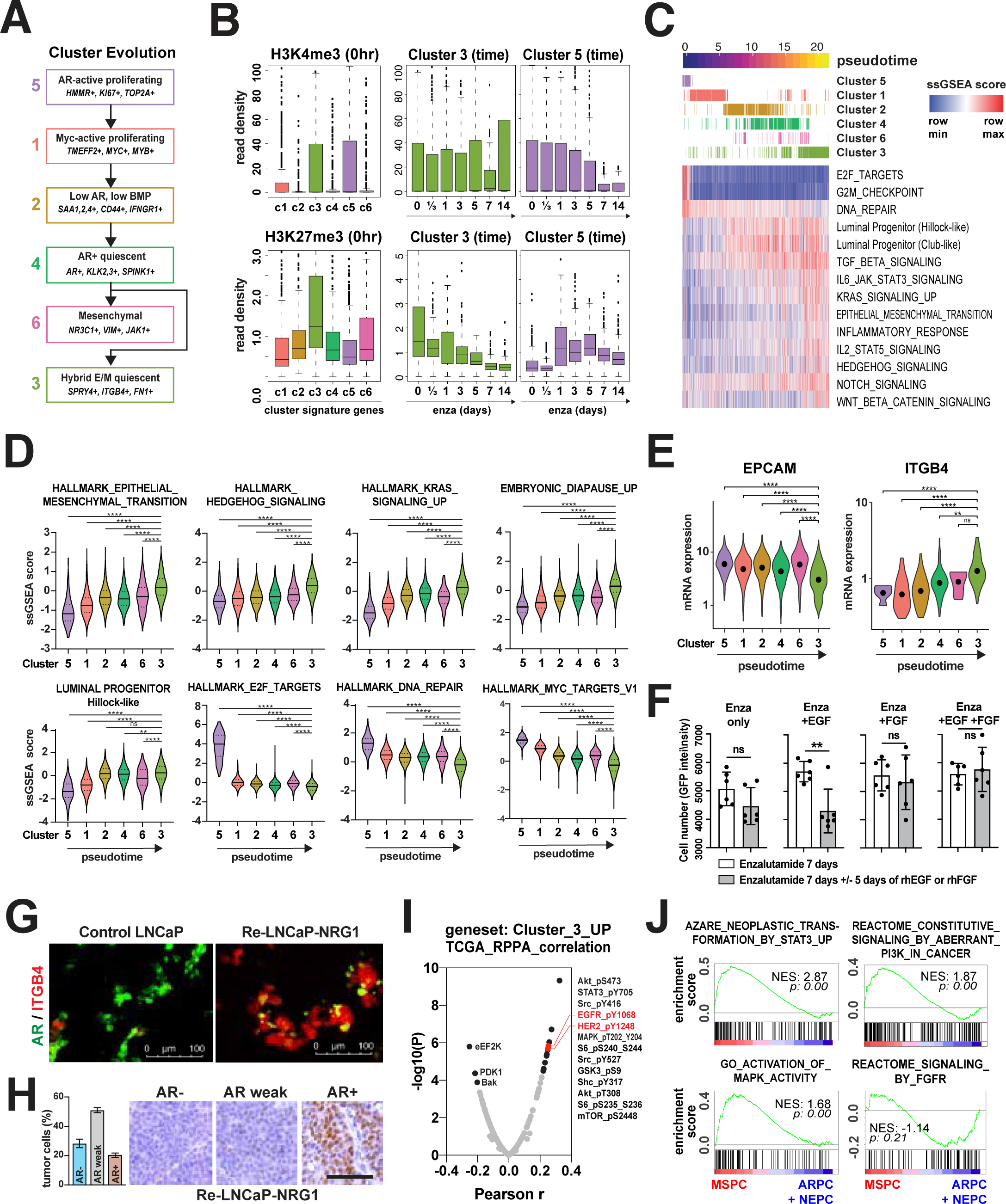

**Suppl. Fig. 5.**
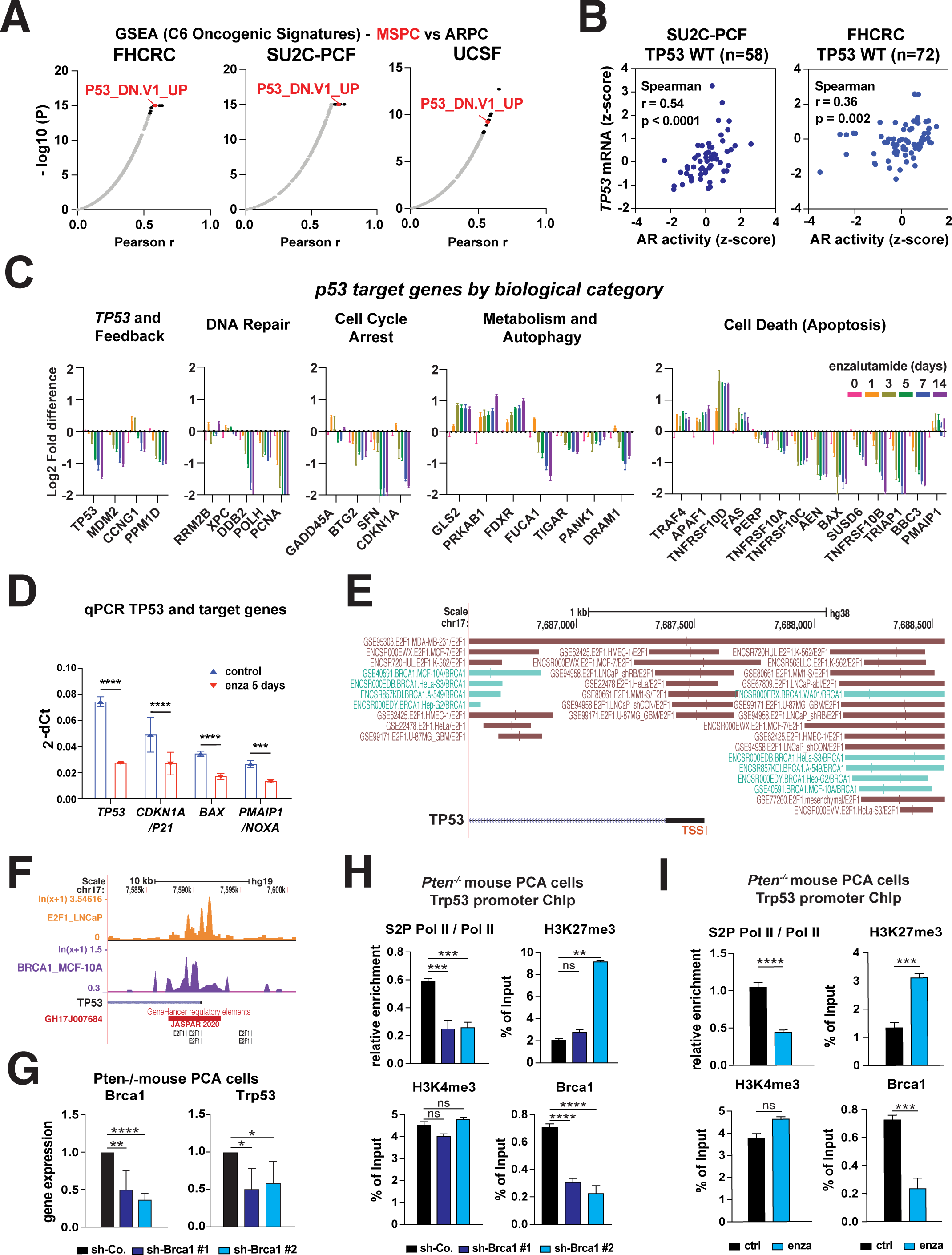

**Suppl. Fig. 6.**
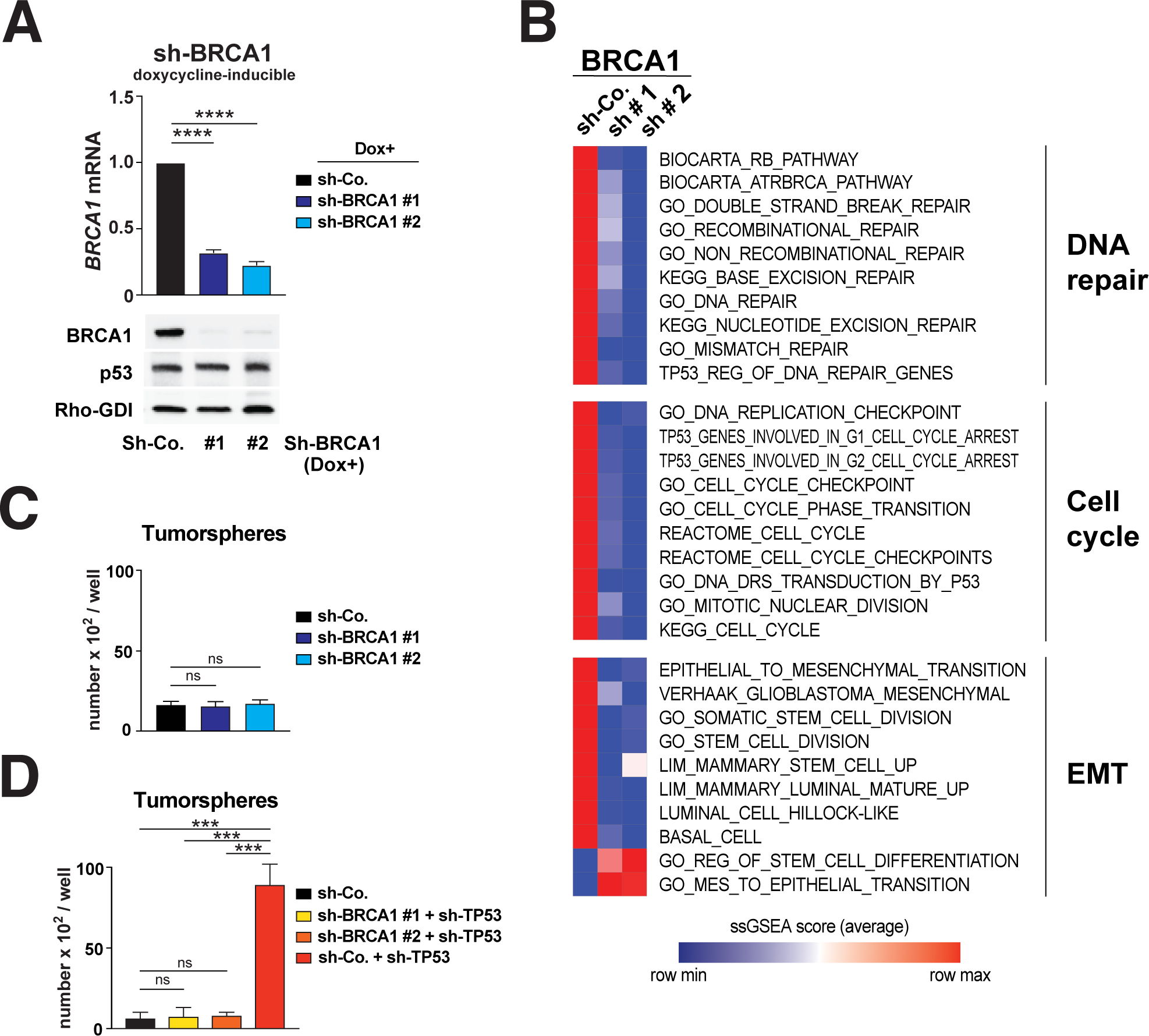

**Suppl. Fig. 7.**
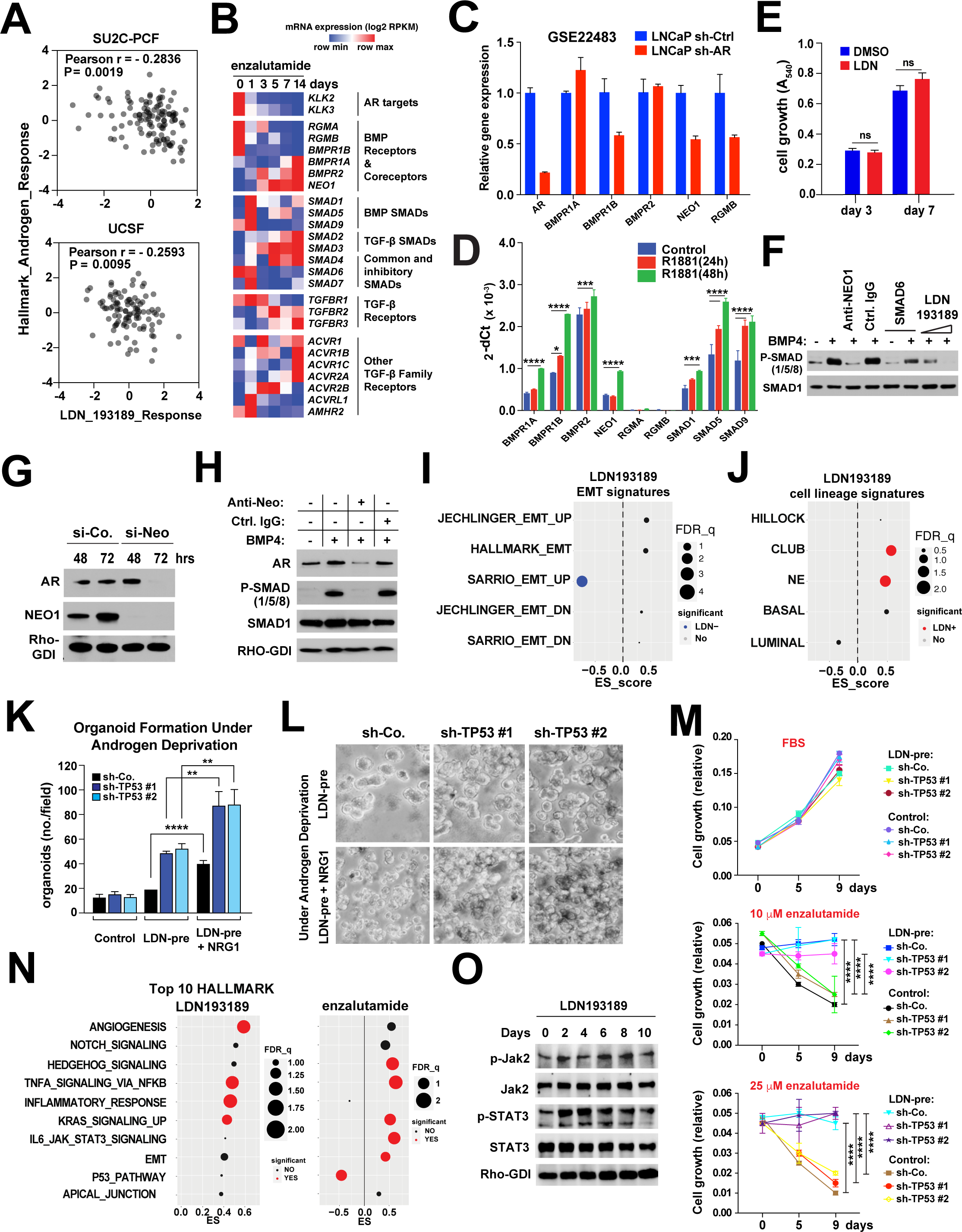

**Suppl. Fig. 8.**
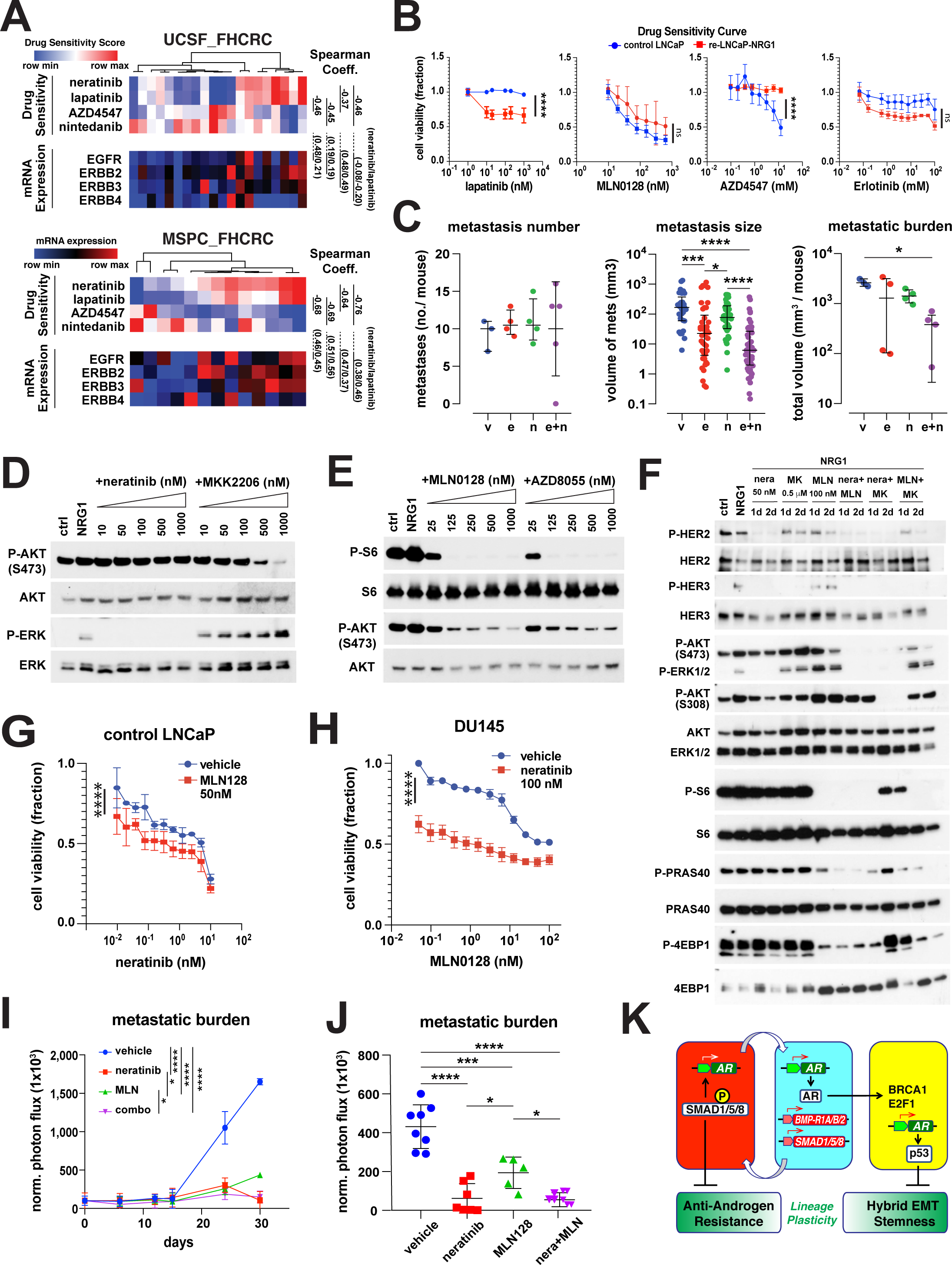

