## Supplementary Figure legend for "Transcriptional Inactivation of TP53 and the BMP Pathway Mediates Therapy-induced Dedifferentiation and Metastasis in Prostate Cancer"

**Supplementary Figure 1.**

**A**. SU2C/PCF M-CRPC whole transcriptome dataset (n of sample: 118, n of transcript: 50,331, Robinson et al., 2015) was downloaded from cBioPortal.org. Batch correction was performed (batch 1: BROAD; batch 2: U MICHIGAN). Unsupervised clustering using Partition Around Medoids (PAM) method resulted in 4 clusters (upper dot plot). Top upregulated genes include: DEFB1, C2orf70, AKR1B10, METTL7B, SERPINA1 in Cluster 1; AC020663.1, AL590710.1, RP11-83M16.5, RP11-136F16.1, AC004815.1 in Cluster 2; C9orf91, ERGIC1, STK39, HOMER2, ACACA in Cluster 3; CXCR3, CD79A, IGHV4-61, IGLV3-25 in Cluster 4. Gene Atlas Analysis was performed by “Gene Enrichment” command of ASAP (Automated Single-cell Analysis Pipeline,<https://asap-old.epfl.ch/>. Gardeux et al., 2017), computing overlap of reference datasets from BioGPS Human Cell Type and Tissue Gene Expression Profiles (Su et al., 2004). Cluster 1: Fetal liver (Odd Ratio (OR): 30.0, adj. p = 0). Cluster 2: Bone marrow (OR: 68.3, adj. p = 7.5e-5). Cluster 3: Prostate (OR: 24.7, adj. p = 1.2e-9). Cluster 4: Lymph node (OR: 545.1 adj. p = 9.7e-10). Actual tumor sites of the samples are shown below (lower dot plot). C# = Cluster #.

**B**. Box lists of five datasets merged into PCO-94. Dataset names, n of samples and sources are indicated. MDA PCa PDX: microarray data of MDA PCa PDX, including tumors of same origin but grown in castrated and uncastrated hosts (133-4_cas1,2; 180-30_cas1,2) (Tzelepi et al., 2012); MSKCC PCa organoid/ODX (organoid-derived xenograft): mRNA expression (RNA Seq FPKM) data available for 10 of 12 PCa organoids (Gao et al., 2014). CCLE PCa Cell lines gene expression (RNA Seq RPKM) data (Release date: 14-Feb-2018. Broad Institute). LuCaP M-CRPC PDXs custom Agilent 44k whole genome expression microarray (includes early/late and castration-resistant passages. GSE93809. Nguyen et al., 2017). LuCaP PDX-derived organoids (RNA Seq TPM) data (includes two repeats. GSE113741. Beshiri et al., 2018).

**C**. (top) PCO-94 dataset consensus plots, comparing PAM clustering results (K=3) and consensus clustering (K=2-5). Algorithm: SOM (self-organizing map); distance measure: Pearson. (bottom left) Consensus Cumulative Distribution Function (CDF) from K=2 to K=10. The proportion of ambiguous clustering (PAC) score was measured by the ΔCDF from u1(0.1) to u2(0.9). (bottom right) PAC score bar graph sorted by the lowest PAC, indicating K=3 as optimal n of clusters.

**D**. Representative top 20 positively (right, red) and negatively (left, blue) correlating gene sets of principal coordinates 1 and 2, shown in Figure 1A. Pearson correlation coefficients of ssGSEA scores and the coordinate values were calculated and presented as volcano plot. The C2 “curated”, the C5 “Gene Ontology” and the H “hallmark” gene sets (from the mSigDB collections) were used in the analysis.

**E**. Normalized Enrichment Score (NES) heatmap of three clusters (continued from Figure 1C). Hallmarks (+) enrichment: NES > +1.0 in one cluster only. Hallmarks (-) enrichment: NES < -1.0 in one cluster only. Prostate Carcinogenesis: (top) Genes up/down-regulated in PCa vs benign prostate tissue (Tomlin et al., 2007; Liu et al., 2006; Wallace et al., 2008) or in PCa cell line but not in normal prostate epithelial or stromal cells (Yegnasubramanian et al., 2008). (bottom) Genes up-regulated in Oncogene activated PCa tumor models (Liu et al., 2008; Azare et al., 2007; Acevedo et al,. 2007).

**F**. Relative fraction of ARPC, MSPC and NEPC in PCO samples, stratified by PAM cluster, sorted by largest fraction size. Samples assigned as “pure” if the largest fraction > 0.75 (white horizontal line). Otherwise assigned as “mixed”.

**G**. Relative fraction of ARPC, MSPC and NEPC in CCLE prostate cancer cell lines. Note that all samples are assigned as “pure”.

**H**. Hierarchical clustering and dendrogram of LuCaP PDXs and Organoids (n=56), aligned with relative fraction prediction from CIBERSORT analysis (bar graph). Note that all repeat samples, early and late passage pairs and castration-resistant derivatives are clustered together in each category. In “mixed” category, LuCaP PDX 86.2 mosaicism has been suggested, as the “ERG expression appeared in in later passages but not in the first four passages” (Nguyen et al., 2017).

**I, J.** Comparison of relative fraction prediction from pairs of tumor organoids and ODXs (organoid-derived xenografts) (**I**) and microarray data versus RNA-seq data of same PDXs (**I**). No statistically significant difference was noted (chi-square test). ns = not significant. MSK-PCa#.o represents “organoid”, MSK-PCa#.m represents “organoid-derived mouse xenograft”.

**Supplementary Figure 2.**

**A.** Left: three-dimensional plots of M-CRPC datasets PCoA analysis. Clustering by Partition Around Medoids (PAM) (K=3). Right: CIBERSORT ARPC, MSPC and NEPC relative fractions calculated from in each cluster. n% indicates proportions of clusters in each dataset. Clusters were numbered by their size. Note that cluster 1 of each dataset has the largest ARPC, cluster 2 MSPC, cluster 3 NEPC. The separations between cluster 1 and 2 were less clear than cluster 3 versus the rest in PCoA plot, consistent with CIBERSORT analysis.

**B**. Comparison of clustering results (shown in Supplementary Figure 2A) to previously proposed M-CRPC classification from Dr. Nelson’s group (Bluemn et al., 2017; Su et al., 2019).

**C**. TIDE analysis (continued from Figure 2D). IFN-gamma activity score of each subtype is shown. P value was calculated by One-way ANOVA. Multiple comparison by Dunnett’s test.

**D**. Common genetic alterations of ARPC, MSPC and NEPC, in SU2C-PCF M-CRPC dataset. Oncoprint was generated in cBioPortal.org (Cerami et al., 2012). DDR = DNA damage repair; u.s.=unknown significance.

**E**. Copy number alterations of chromosome 3q26 in MSK-PCa Organoids and CCLE PCa cell lines. Segment view by IGV (hg19). Right: matching relative fraction data is shown.

**F**. Gene mutation frequencies (%) in primary adenocarcinoma molecular subtypes. Chi-square test.

**G**. AR and ANXA1 protein expressions in TCGA primary adenocarcinoma molecular subtypes. Expression z-score from RPPA. One-way ANOVA. Multiple comparison by Dunnett’s test.

**H**. Copy number alteration (CNA) frequencies (%) of TP53, PTEN and RB1 in TCGA primary adenocarcinoma molecular subtypes. Chi-square test.

**I-K**. Distributions of Gleason score (I), Tumor stage (J) and Nodal stage (K) in primary adenocarcinoma molecular subtypes. If radical prostatectomy data was available, then surgical specimen Gleason score and pathologic T/N stage were used. Chi-square test.

**Supplementary Figure 3.**

**A.** Cell proliferation rate (average n of cells per condition, upper) and apoptotic cell rate (early apoptosis (annexin v+ PI+) and late apoptosis (annexin v+ PI-) by flow cytometry, lower) in LNCaP cells treated with enzalutamide for 0 to 14 days.

**B.** Principal Coordinate Analysis (PCoA) plot of LNCaP enzalutamide time-series RNA-seq data merged with “PCO-94” original dataset. X axis: drug incubation time. Y axis: coordinate 2 value. Three samples per time point. ssGSEA scores of G1/S transition and mTORC1 signaling were top-2 gene sets correlation with coordinate 2.

**C, D.** VCaP enzalutamide time-series in vitro. **C**. Western blot of AR, AR-v7, AR target gene TMPRSS2, EMT and stemness markers. **D**. immunofluorescence staining of AR, ITGB4 and EpCAM in enzalutamide-treated VCaP cells.

**E, F.** MDA-PCa-163-A enzalutamide time-series in vitro (**E**) an in vivo (**F**). **E**. Western blot of AR, target gene, EMT and stemness markers. **F**. immunohistochemistry staining images of indicated proteins in enzalutamide-treated or control (untreated) tumors. H = Hematoxylin.

**G-J.** Analysis of LNCaP enzalutamide time-series ATAC-seq data. **G.** Principal Component Analysis (PCA) plot. Duplicates per time point. X axis component 1 called as “Major” peak, and Y axis component 2 called as “Minor” peak in following analysis. Inner donut plot: total variability explained (%). “Major” peak was defined by those variability primarily explained by PCA component 1. **H.** “Major” peak sites summit. Average of duplicates. **I.** Heatmap of ATAC peaks in the -1 to +1 kb around “Major” enhancer sites (left) and “Major” promoter sites (right). **J.** ATAC-seq peak summits of EMT-related genes and stemness-related genes.

**K**. H3K4me3 and H3K27me3 signals over the promoters of EMT and stemness-related genes in LNCaP cells treated with enzalutamide (0 to 14 days).

**L**. LNCaP cells treated with enzalutamide ssGSEA score heatmap, comparing Day 0 samples and Day 14 samples. The hallmark gene sets were used.

**Supplementary Figure 4.**

**A.** LNCaP cell state clusters gene expression characteristics with cell-fate evolutionary trajectory order.

**B**. Pre and post-enzalutamide treatment promoter histone marks state. H3K4me3 (upper) and H3K27me3 (lower) peaks average of cluster 1 to 6 upregulated genes in LNCaP enzalutamide time-series assessed by histone marker Chip-sequencing. Daily average peak level changes of cluster 3 (green) and 5 (purple) are shown (right).

**C.** Heatmap of ssGSEA scores (overlapping in panel D) in each cluster. Columns aligned by pseudotime. Prostate luminal progenitor genesets from Henry et al., 2018. The others: Hallmark genesets.

**D**. Violin plots of ssGSEA scores – hallmarks EMT, Hedeghog, Ras, embryonic diapause, prostate luminal progenitor, hallmarks E2F target, DNA-repair and Myc target genesets in each cluster. P value was calculated by One-way ANOVA. Multiple comparison by Dunnett’s test.

**E**. Violin plots of *EPCAM* and *ITGB4* gene expressions in cluster 1 to 6. P value was calculated by One-way ANOVA. Multiple comparison by Dunnett’s test.

**F.** Cell proliferation (GFP intensity) of LNCaP^GFP^ cells of 7-day enzalutamide treatment only or followed by 5 days of recombinant human EGF 20ng/ml or FGF 40ng/ml. Welch’s t-test. **p < 0.01; ns = not significant.

**G.** Co-immunofluorescence staining images of AR and ITGB4 in LNCaP cells or re-LNCaP-NRG1 cells.

**H.** AR and Integrin β4 expressions in reprogrammed LNCaP cell adrenal gland metastases. Representative areas from AR- (mice #181) and AR+ tumors (#182) are shown.

I. Volcano plot of TCGA prostate adenocarcinoma samples cluster_3_up geneset ssGSEA score and RPPA protein expression z-score.

**J**. Enrichment plots of STAT3, PI3K, MAPK or FGFR activation signatures in SU2C-PCF MSPC subtype versus the rest. NES = normalized enrichment score.

**Supplementary Figure 5.**

**A.** Top 10 genesets (black dots) of which the ssGSEA score positively correlate with MSPC proportion estimates (%) from the FHCRC, SU2C-PCF and UCSF metastatic CRPC datasets. P53_DN.V1_UP (red dot) was ranked within top 10 in all three datasets. Parental gene sets are the C6 Oncogenic Signatures (n=189) of the MSigDB Collections.

**B.** Scatter plot of *TP53* mRNA expression z-score and AR activity z-score of samples of wild type *TP5*3 (no mutation, diploid, n=58) from the SU2C-PCF dataset and FHCRC dataset. AR activity was measured by ssGSEA of the HALLMARK_ANDROGEN_RESPONSE geneset.

**C**. Log2 transformed fold differences of the TP53 and target genes FPKM values from RNA-seq of LNCaP cells treated with 10uM enzalutamide for 0, 1, 3, 5, 7 or 14 days. Target gene list acquired from Fischer M. Census and evaluation of p53 target genes. Oncogene. 2017 Jul 13;36(28):3943-3956.

**D.** TP53 and its regulatory target genes mRNA levels by qRT-PCR in LNCaP cell treated with or without enzalutamide, 10uM, for 5 days. error bars indicate mean ± SD. **p<0.01, ***p<0.005, ****p<0.001.

**E, F.** BRCA1 and E2F1 ChIPseq peak profile on TP53 promoter region. ChIPseq data acquired from in Remap 2020 of the ENCODE project were aligned on around TP53 transcriptional start site (TSS) (**E**). Representative peak data presented in **F**.

**G**. Relative gene expression of *Brca1* (left panel) and *Trp53* (right panel) mRNAs in Pten-P8^(-/-)^ cells stably transduced with the indicated hairpins. Error bars, mean±SD of triplicate experiments, **p<0.01, ***p<0.005, **** p<0.0001 two-tailed Student t test.

**H, I**. Relative occupancy of RNA Pol II Ser2p and enrichment of H3K4me3, H3K27me3, *Brca1* on the *Trp53* promoter from ChIP-qPCR in *Brca1*-KD (**H**) or enzalutamide treated (**I**) Pten-P8^(-/-)^ cells; error bars indicate mean ± SD. **p<0.01, ***p<0.005, ****p<0.001.

**Supplementary Figure 6.**

**A**. Relative gene expression of BRCA1 mRNAs (upper) and western blot of indicated antibodies (lower) of BRCA1-stably silenced LNCaP cells. Error bars, mean±SD of triplicate experiments, **** p<0.0001 two-tailed Student t test.

**B**. Ranks ssGSEA heatmap of Sh-Ctrl and Sh-*BRCA1* LNCaP cells. Genes signatures were categorized in relevance to DNA repair, cell cycle, EMT and stemness, respectively.

**C, D.** Tumor sphere formation ability was compared among control (Sh-Co.), BRCA1 knockdown LNCall cells (**C**), or control, P53 knockdown LNCaP cells (Sh-TP53), and P53- BRCA1- double knockdown (sh-TP53+ BRCA1-#1; sh-TP53+ BRCA1-#2) (**D**). Cells were plated in triplicate in 24 well ultra-low attachment plates at a seeding density of 3000 cells/well (sphere assay). Sphere numbers were counted after 10 days. Error bars, mean ± SE. For statistical analyses, unpaired two-tailed Student’s t-tests was performed.

**Supplementary Figure 7.**

**A.** Scatter plots showing correlation of ssGSEA scores of the hallmark androgen response and LDN193189 response in the SU2C-PCF dataset (upper) and UCSF dataset (lower).

**B**. Heatmap of BMP-SMAD and TGF-SMAD signaling components gene expression averages in LNCaP cells treated with enzalutamide 10uM for 0 to 14 days. Three replicates per condition.

**C.** BMP-SMAD signaling components mRNA expression levels measured by cDNA microarray of late passage LNCaP cells with sh-Ctrl and sh-AR. Data from GSE22483 (Gonit et al., Mol Endocrinol, 2011).

**D.** BMP-SMAD signaling components mRNA expression levels measured by qPCR in LNCaP cells treated with DMSO or R1881 100pM for 48 hrs. ns=not significant; * p<0.05. Student’s *t-*test.

**E.** Cell proliferation assay of LNCaP cells treated with or without LDN193189 for the indicated time points (days).

**F**. Phosphorylated SMAD1/5/8 immunoblots in serum starved parental LNCaP cells or SMAD6 overexpressed LNCaP cells either treated with BMP4 (100 ng/mL, 30 mins) or pretreated with anti-NEO1 blocking antibody (5ug/mL) or control IgG or LDN193189 for indicated concentration for 2 hrs followed by treated with BMP4. Total SMAD 1 was used as internal control.

**G**. AR protein immunoblots in total lysates of LNCaP cells transfected with control or human Neo1 specific siRNA (SMART pool) at the indicated time point.

**H**. AR protein immunoblots in total lysates of LNCaP cells pretreated with anti-NEO1 blocking antibody (5ug/mL) or control IgG and treated with BMP4 (100 ng/mL, 30 mins).

**I, J.** ClusterProfiler dot plot showing EMT signatures enriched (**I**) or stemness signatures enriched (**J**) using gene set enrichment analysis (GSEA) for genes differentially expressed in control compared to LDN193189 treatment condition.

**K, L.** Quantification (**K**) and representative images (**L**) of control and TP53-silenced LNCaP cells subjected to organoid formation assay. The indicated cells were pretreated with or without LDN193189 50nM and/or rh-NRG1 for 8 days. Then 100 –10,000 dissociated cells from each condition were plated into wells of ultra-low attachment 96 well plates. Organoid number per field (left) and diameter in micrometer (right) were counted and measured at day 10. Scale bar=100 μm. Error bars, mean±SD of triplicate experiments, **** p<0.0001 two-tailed Student t test.

**M.** Relative cell growth of LNCaP sh-Co., sh-TP53#1, sh-TP53#2 cells grown in control or LDN-193189 (pretreated for 8 days) and in the conditions of CSS alone, CSS+ENZA (10uM) and CSS+ENZA (25uM) for additional 9 days.

**N.** Clusterprofiler dot plot showing the top10 enriched signatures from Hallmark gene sets using gene set enrichment analysis (GSEA) for genes differentially expressed in control compared to LDN193189 treatment condition (left), or in control compared to enzalutamide 7days treatment condition (right). X axis title “ES” represents the GSEA enrichment score. Y axis represents the name of the signatures. Dot size represents the -log10 (FDR_q_value + 0.001). Dot color represents the significance.

**O.** Immunoblotting of JAK2, STAT3 proteins and their phosphorylated form in LNCaP cells treated with LDN193189 100nM for 0-10 days.

**Supplementary Figure 8.**

**A.** Predicted Drug sensitivity heatmaps of MSPC samples from the FHCRC and the UCSF datasets. mRNA expressions of ERBB1-4 are shown together. Spearman correlation coefficients between neratinib/lapatinib and FGF inhibitors AZD4547/nintedanib sensitivity scores or the mRNA expressions are shown (right).

**B.** Cell viability dose-response curves of LNCaP control and re-LNCaP-NRG1 cells to lapatinib, MLN0128 or AZD4547. Error bars, mean ±SD of 8 replicates, **** p<0.0001 Welch's unequal variances t-test of AUC.

**C.** Tumor numbers, sizes and burden (total sum of sizes) of re-LNCaP-NRG1 metastases in castrated male NSG mice treated with daily enzalutamide 10mg/kg and/or neratinib 40mg/kg for 4 weeks. v = vehicle; e = enzalutamide; n = neratinib.

**D.** Immunoblotting of AKT and ERK1/2 protein phosphorylation upon increasing neratinib or MKK2206 drug concentration in re-LNCaP-NRG1 cells.

**E.** Immunoblotting of AKT and S6 protein phosphorylation upon different dosage of mTOR kinase inhibitors MLN0128 and AZD8055 in re-LNCaP-NRG1 cells.

**F.** Immunoblotting of HER2/3, ERK1/2, AKT and mTOR kinase pathway protein phosphorylation upon long-term (1day or 2days) combined treatments of neratinib (nera) 50nM, MK2206 (MK) 0.5uM or MLN0128 (MLN) 100nM in re-LNCaP-NRG1 cells.

**G.** Cell viability dose response curve of LNCaP cells upon MLN1028 50nM and/or varying dose of neratinib. Error bars, mean ±SD of 8 replicates, **** p<0.0001 Welch's unequal variances t-test of AUC.

**H.** Cell viability dose response curve of DU145 cells upon neratinib 100nM and/or varying dose of MLN1028. Error bars, mean ±SD of 8 replicates, **** p<0.0001 Welch's unequal variances t-test of AUC.

**I.** Changes of normalized photon flux from DU145 metastatic tumors in castrated male NSG mice treated with daily neratinib 40mg/kg and/or MLN0128 0.3mg/kg for 4 weeks.

**J.** Normalized photon flux from bioluminescence images of week 4 DU145 metastatic tumors in castrated male NSG mice treated with daily neratinib 40mg/kg and/or MLN0128 0.3mg/kg for 4 weeks.

**K**. Graphical Model.

**Supplementary Figure References**

**Acevedo VD**, Gangula RD, Freeman KW, Li R, Zhang Y, Wang F, Ayala GE, Peterson LE, Ittmann M, Spencer DM. Inducible FGFR-1 activation leads to irreversible prostate adenocarcinoma and an epithelial-to-mesenchymal transition. Cancer Cell. 2007 Dec;12(6):559-71.

**Azare J**, Leslie K, Al-Ahmadie H, Gerald W, Weinreb PH, Violette SM, Bromberg J. Constitutively activated Stat3 induces tumorigenesis and enhances cell motility of prostate epithelial cells through integrin beta 6. Mol Cell Biol. 2007 Jun;27(12):4444-53. Epub 2007 Apr 16.

**Beshiri ML**, Tice CM, Tran C, Nguyen HM et al. A PDX/Organoid Biobank of Advanced Prostate Cancers Captures Genomic and Phenotypic Heterogeneity for Disease Modeling and Therapeutic Screening. Clin Cancer Res 2018 Sep 1;24(17):4332-4345.

**Bluemn EG**, Coleman IM, Lucas JM, Coleman RT, Hernandez-Lopez S, Tharakan R, Bianchi-Frias D, Dumpit RF, Kaipainen A, Corella AN, Yang YC, Nyquist MD, Mostaghel E, Hsieh AC, Zhang X, Corey E, Brown LG, Nguyen HM, Pienta K, Ittmann M, Schweizer M, True LD, Wise D, Rennie PS, Vessella RL, Morrissey C, Nelson PS. Androgen Receptor Pathway-Independent Prostate Cancer Is Sustained through FGF Signaling. Cancer Cell. 2017 Oct 9;32(4):474-489.e6.

**Cerami E**, Gao J, Dogrusoz U, Gross BE, Sumer SO, Aksoy BA, Jacobsen A, Byrne CJ, Heuer ML, Larsson E, Antipin Y, Reva B, Goldberg AP, Sander C, Schultz N. The cBio cancer genomics portal: an open platform for exploring multidimensional cancer genomics data. Cancer Discov. 2012 May;2(5):401-4.

**Gardeux V**, David FPA, Shajkofci A, Schwalie PC, Deplancke B. ASAP: a web-based platform for the analysis and interactive visualization of single-cell RNA-seq data. Bioinformatics. 2017 Oct 1;33(19):3123-3125.

**Gao D**, Vela I, Sboner A, Iaquinta PJ, Karthaus WR, Gopalan A, Dowling C, Wanjala JN, Undvall EA, Arora VK, Wongvipat J, Kossai M, Ramazanoglu S, Barboza LP, Di W, Cao Z, Zhang QF, Sirota I, Ran L, MacDonald TY, Beltran H, Mosquera JM, Touijer KA, Scardino PT, Laudone VP, Curtis KR, Rathkopf DE, Morris MJ, Danila DC, Slovin SF, Solomon SB, Eastham JA, Chi P, Carver B, Rubin MA, Scher HI, Clevers H, Sawyers CL, Chen Y. Organoid cultures derived from patients with advanced prostate cancer. Cell. 2014 Sep 25;159(1):176-187.

**Henry GH**, Malewska A, Joseph DB, Malladi VS, Lee J, Torrealba J, Mauck RJ, Gahan JC, Raj GV, Roehrborn CG, Hon GC, MacConmara MP, Reese JC, Hutchinson RC, Vezina CM, Strand DW. A Cellular Anatomy of the Normal Adult Human Prostate and Prostatic Urethra. Cell Rep. 2018 Dec 18;25(12):3530-3542.e5.

**Liu Y**, Mo JQ, Hu Q, Boivin G, Levin L, Lu S, Yang D, Dong Z, Lu S. Targeted overexpression of vav3 oncogene in prostatic epithelium induces nonbacterial prostatitis and prostate cancer. Cancer Res. 2008 Aug 1;68(15):6396-406.

**Nguyen HM**, Vessella RL, Morrissey C, Brown LG, Coleman IM, Higano CS, Mostaghel EA, Zhang X, True LD, Lam HM, Roudier M, Lange PH, Nelson PS, Corey E. LuCaP Prostate Cancer Patient-Derived Xenografts Reflect the Molecular Heterogeneity of Advanced Disease and Serve as Models for Evaluating Cancer Therapeutics. Prostate. 2017 May;77(6):654-671.

**Su AI**, Wiltshire T, Batalov S, Lapp H, Ching KA, Block D, Zhang J, Soden R, Hayakawa M, Kreiman G, Cooke MP, Walker JR, Hogenesch JB. A gene atlas of the mouse and human protein-encoding transcriptomes. Proc Natl Acad Sci U S A. 2004 Apr 20;101(16):6062-7.

**Su W**, Han HH, Wang Y, Zhang B, Zhou B, Cheng Y, Rumandla A, Gurrapu S, Chakraborty G, Su J, Yang G, Liang X, Wang G, Rosen N, Scher HI, Ouerfelli O, Giancotti FG. The Polycomb Repressor Complex 1 Drives Double-Negative Prostate Cancer Metastasis by Coordinating Stemness and Immune Suppression. Cancer Cell. 2019 Aug 12;36(2):139-155.e10.

**Tomlins SA**, Mehra R, Rhodes DR, Cao X, Wang L, Dhanasekaran SM, Kalyana-Sundaram S, Wei JT, Rubin MA, Pienta KJ, Shah RB, Chinnaiyan AM. Integrative molecular concept modeling of prostate cancer progression. Nat Genet. 2007 Jan;39(1):41-51.

**Tzelepi V**, Zhang J, Lu JF, Kleb B, Wu G, Wan X, Hoang A, Efstathiou E, Sircar K, Navone NM, Troncoso P, Liang S, Logothetis CJ, Maity SN, Aparicio AM. Modeling a lethal prostate cancer variant with small-cell carcinoma features. Clin Cancer Res. 2012 Feb 1;18(3):666-77.

**Yegnasubramanian S**, Haffner MC, Zhang Y, Gurel B, Cornish TC, Wu Z, Irizarry RA, Morgan J, Hicks J, DeWeese TL, Isaacs WB, Bova GS, De Marzo AM, Nelson WG. DNA hypomethylation arises later in prostate cancer progression than CpG island hypermethylation and contributes to metastatic tumor heterogeneity. Cancer Res. 2008 Nov 1;68(21):8954-67.

**Wallace TA**, Prueitt RL, Yi M, Howe TM, Gillespie JW, Yfantis HG, Stephens RM, Caporaso NE, Loffredo CA, Ambs S. Tumor immunobiological differences in prostate cancer between African-American and European-American men. Cancer Res. 2008 Feb 1;68(3):927-36.
